# Uncovering the Neural Correlates of the Urge-to-Blink: A Study Utilising Subjective Urge Ratings and Paradigm Free Mapping

**DOI:** 10.1101/2024.07.19.603913

**Authors:** Mairi S. Houlgreave, Eneko Uruñuela, César Caballero-Gaudes, Penny Gowland, Katherine Dyke, Valerie Brandt, Imaan Mohammed, Rosa Sanchez Panchuelo, Stephen Jackson

**Affiliations:** School of Psychology, University of Nottingham, University Park, Nottingham, NG7 2RD, UK; Sir Peter Mansfield Imaging Centre, School of Physics and Astronomy, University of Nottingham, University Park, Nottingham, NG7 2RD, UK; Basque Centre on Cognition, Brain and Language; Department of Psychology, Centre for Innovation in Mental Health, University of Southampton; Clinic of Psychiatry, Social Psychiatry and Psychotherapy, Hannover Medical School, Hannover, Germany; Institute of Mental Health, School of Medicine, University of Nottingham, University Park, Nottingham, NG7 2RD, UK

**Author notes:** Correspondence to: Mairi Houlgreave, School of Psychology, University of Nottingham, University Park, Nottingham, NG7 2RD, UK. E-mail address (M.S. Houlgreave).

**Keywords:** Urge, Blinking, Suppression, functional Magnetic Resonance Imaging, Paradigm free mapping, Insula

## Abstract

Neuroimaging plays a significant role in understanding the neurophysiology of Tourette syndrome (TS), in particular the main symptom, tics, and the urges associated with them. Premonitory urge is thought to be a negative reinforcer of tic expression in TS. Tic expression during neuroimaging is most often required as an overt marker of increased urge-to-tic, which can lead to considerable head movement, and thus data loss. This study aims to identify the brain regions involved in urge in healthy subjects using multi-echo functional MRI and a timing- free approach to localise the BOLD response associated with the urge-to-act without information of when these events occur. Blink suppression is an analogous behaviour that can be expressed overtly in the MRI scanner which gives rise to an urge like those described by individuals with TS.

We examined the urge-to-blink in 20 healthy volunteers with an experimental paradigm including two conditions, “Okay to blink” and “Suppress blinking”, to identify brain regions involved in blink suppression. Multi-echo functional MRI data was analysed using a novel approach to investigate the BOLD signal correlated with the build-up of the urge-to-blink that participants continuously reported using a rollerball device. In addition, we used the method of multi-echo paradigm free mapping (MESPFM) to identify these regions without prior specification of task timings.

Subjective urge scores were correlated with activity in the right posterior and ventral-anterior insula as well as the mid-cingulate and occipital cortices. Furthermore, blink suppression was associated with activation in the dorsolateral prefrontal cortex, cerebellum, right dorsal-anterior insula, mid-cingulate cortex and thalamus. These findings illustrate that different insula subregions contribute to the urge-for-action and suppression networks. The MESPFM approach showed co-activation of the right insula and cingulate cortex. The MESPFM activation maps showed the highest overlap with activation associated with blink suppression, as identified using general linear model analysis, demonstrating that activity associated with suppression can be determined without prior knowledge of task timings.

## 1.1 Introduction

In contrast to other movement disorders, many individuals with Tourette syndrome (TS) can temporarily suppress their tics (Robertson, 2011). However, the majority experience unpleasant sensations that build up in intensity until the tic is released (Kwak et al., 2003; Leckman et al., 1993). These urges can manifest as sensations such as pressure, itching, numbness, or aching (Kwak et al., 2003; Woods et al., 2005), and are often used in behavioural therapies to predict and pre-empt tics (Azrin & Nunn, 1973). One key mechanistic question is whether tics are voluntary and function to alleviate premonitory urge (PU) (Leckman et al., 1993), which could act as a negative reinforcer of tic behaviour (Capriotti et al., 2014), or whether urges arise due to the act of suppression, much like the sensation experienced when suppressing a yawn (Jackson et al., 2011).

Previous research into the generation of tics and PU has suggested the involvement of separate networks. A functional magnetic resonance imaging (fMRI) study by Bohlhalter and colleagues showed that the primary sensorimotor cortex and the cerebellum are active at tic onset, whereas the insula and premotor regions are active just before a tic, suggesting either an involvement in PU or in movement preparation (Bohlhalter et al., 2006).

It has been theorised that the urge-to-act may involve a loop comprising the anterior insula, the mid-cingulate cortex (MCC) and the mid-insula (Jackson et al., 2011), where activation of this pathway would lead to urge sensation, initiation of an action in response to the urge and finally assessment of whether the urge has been fulfilled. Research into addictive behaviours such as smoking has shown that patients with brain injuries involving the insula were more likely to report a reduction in the urge-to-smoke compared to smokers with damage in other loci (Naqvi et al., 2007). Furthermore, sensations such as scratching, numbness, and warmth in distinct body parts can be elicited with direct stimulation of the contralateral insula (Penfield & Faulk, 1955). A recent study found that the grey matter value of voxels in the posterior right insula showed a negative association with motor tic severity scores, whereas a region in the anterior dorsal/mid insula was positively correlated with PU scores, suggesting that different portions of the insula may have different roles in tics and urges (Jackson et al., 2020). The anterior insula is known to be involved in interoceptive processing; thus, PU may manifest due to increased awareness of internal sensations (Craig, 2002, 2009). Similarly, it has been proposed that the mid-insula has a role in subjective feelings relating to movement and therefore could establish whether the urge-to-act has been fulfilled (Craig, 2009; Jackson et al., 2011). On the other hand, complex motor responses can be evoked by stimulation of the anterior MCC, which demonstrates that the region could have a role in the execution of actions performed in response to an urge (Caruana et al., 2018; Jackson et al., 2011).

The neural correlates of the urge-to-move have also been investigated in healthy participants with experimental paradigms involving the suppression of common behaviours such as blinking and yawning (Berman et al., 2012; Lerner et al., 2009; Mazzone et al., 2010; Nahab et al., 2009; Yoon et al., 2005). These behaviours give rise to an urge similar to those described by TS patients (Berman et al., 2012; Botteron et al., 2019). A variety of areas including the cingulate cortex, insulae, prefrontal cortex (PFC) and temporal gyri have shown activation associated with urges (Berman et al., 2012; Lerner et al., 2009; Mazzone et al., 2010; Nahab et al., 2009; Yoon et al., 2005). Using a meta-analytic approach, Jackson and colleagues revealed that there is an overlap in activity in the MCC and the right insula during the urge-to-act in healthy participants for a variety of behaviours and the urge-to-tic in patients (Jackson et al., 2011). Therefore, when investigating the network involved in PU, blinking can be used for analogous investigation in healthy controls (Jackson et al., 2011).

The issue with investigating PU is their temporal correlation with motor preparation. Usually in fMRI studies looking at the neural correlates of TS, tics are identified post-hoc using video recordings (Bohlhalter et al., 2006; Neuner et al., 2014), which is subjective and time-consuming. Regions involved in the urge-to- tic can then be identified by looking at regions that are active just before a tic, but this will also identify regions involved in tic generation (Bohlhalter et al., 2006; Neuner et al., 2014). Furthermore, a high proportion of fMRI data are lost during tics, for example due to concomitant head jerks (Bohlhalter et al., 2006; Neuner et al., 2014), however if participants are asked to suppress their tics there would be no overt marker of increased urge-to-tic and, mechanisms involved in tic suppression will be present in the results.

To separate the networks involved in urge and action suppression, we investigated the urge-to-blink in healthy controls performing a blink suppression paradigm. Subjects were asked to continuously rate feelings of urge so that the blood-oxygen level-dependent (BOLD) signal could be modelled with a general linear model (GLM) based on these subjective ratings, which will allow us to identify a network associated with the urge. We also compared ‘Okay to blink’ and blink suppression blocks to highlight regions involved in action suppression, where we expected to show activation in the right inferior frontal gyrus (IFG) (Aron et al., 2004, 2014).

Nevertheless, using a conventional GLM approach will involve averaging across many trials to increase the signal-to-noise ratio (SNR). This assumes that the response is the same for every trial and that the timings are known a priori to establish the hypothesised model for the fMRI signal. In practice, events such as tics and urges are spontaneous and vary in duration as well as in phenotype across time and between participants.

To overcome these assumptions, we also analysed fMRI data with a Paradigm Free Mapping (PFM) approach where the neuronal activity underlying single-trial BOLD events is estimated without prior knowledge of event timings or durations by solving a hemodynamic deconvolution (inverse) problem (Caballero Gaudes et al., 2013; Uruñuela et al., 2023).

It is expected that both the conventional and PFM analyses will detect regions previously identified as being part of the urge network including the MCC and right insula (Jackson et al., 2011). If the same regions can be identified without specification of task timings, this would validate the use of PFM in fMRI studies that aim to characterise urge networks in disorders such as TS. This is important for TS research as, due to the caveats of movement during conventional neuroimaging, moments of heightened urge cannot be identified, and networks involved in the urge-to-tic and tic suppression cannot be disentangled. PFM could allow these networks to be separated without the need for continuous urge ratings.

The primary aim of this study was to identify the BOLD signal correlated with the build-up of the urge-to-blink that participants continuously reported using a rollerball device. The secondary aim of this study was to validate the use of a multi-echo sparse paradigm free mapping (MESPFM) algorithm (Caballero-Gaudes et al., 2019) to identify activation during a blink suppression paradigm, before applying it to covert responses such as the urge-to-tic.

## 1.2 Methods

### 1.2.1 Participants

Twenty-two healthy participants were screened for counterindications for MRI, use of medication and history of neurological or psychiatric disorders. One participant (male, 21 years old, right-handed) was excluded before data analysis due to a technical issue which led to the loss of the fMRI data, and one participant (female, 28 years old, right-handed) was excluded during analysis due to excessive movement. Handedness for the remaining twenty subjects (13 female, mean age (± standard deviation (SD)) = 28 ± 5.2 years) was determined using the Edinburgh Handedness Inventory (18 right-handed, 2 ambidextrous; mean = (± standard deviation (SD)) = 80 ± 31.7, range = -35 to 100) (Oldfield, 1971). Subjects gave informed consent and the study received local ethics committee approval.

### 1.2.2 fMRI Task

All subjects underwent three 7-minute fMRI runs of the same task. The experimental task was based on a previous study by Brandt and colleagues which recorded real-time urge ratings (Brandt et al., 2016) and was implemented in Psychopy2 (1.83.04) (Peirce et al., 2019). Eyeblinks during each run were captured with an MR-compatible camera "12M-i" with integrated LED light mounted on the head coil (MRC systems GmbH) (half frame rate=60Hz). A projected screen displaying the task was visible by a mirror positioned above the participants’ eyes (Figure 2). For the first 30 seconds an instruction was displayed to move an MR-compatible trackball (Cambridge Research Systems) (sampling rate of 10Hz) randomly using their right-hand (‘Random’). This was followed by alternating 60-second runs of ‘Okay to blink’ and ‘Suppress’. During these conditions, participants continuously rated their urge-to-blink on a scale of 0-100 while following instructions to either blink normally or to suppress their blinks, respectively. The ‘Random’ baseline was repeated during the last 30 seconds of the run. Participants were instructed to pay attention to the instructions displayed at the top of the screen and during the ‘Suppress’ condition to return to suppressing their blinks should any escape blinks occur (Berman et al., 2012; Lerner et al., 2009; Stern et al., 2020). Previous studies have shown that 60 seconds of action suppression is achievable and induces feelings of urge (Lerner et al., 2009; Stern et al., 2020). The order of ‘Okay to blink’ and ‘Suppress’ blocks was randomly counterbalanced to reduce order effects, with 50% of participants starting with suppression following the initial baseline. All participants moved the trackball using their right hand regardless of hand dominance.

**Figure 1.**
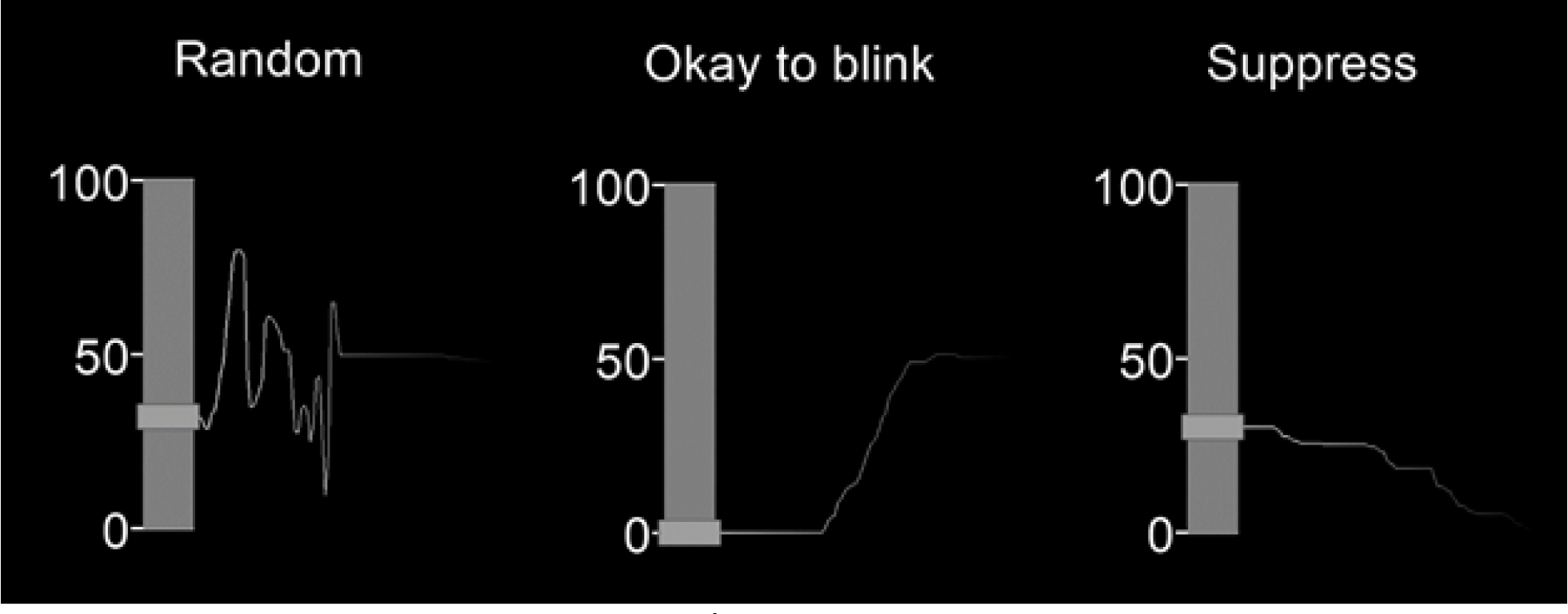
The real-time urge task display. A figure displaying the real-time urge monitor, with urge rated on a scale of 0-100 and nstructions for each condition displayed above.

**Figure 2.**
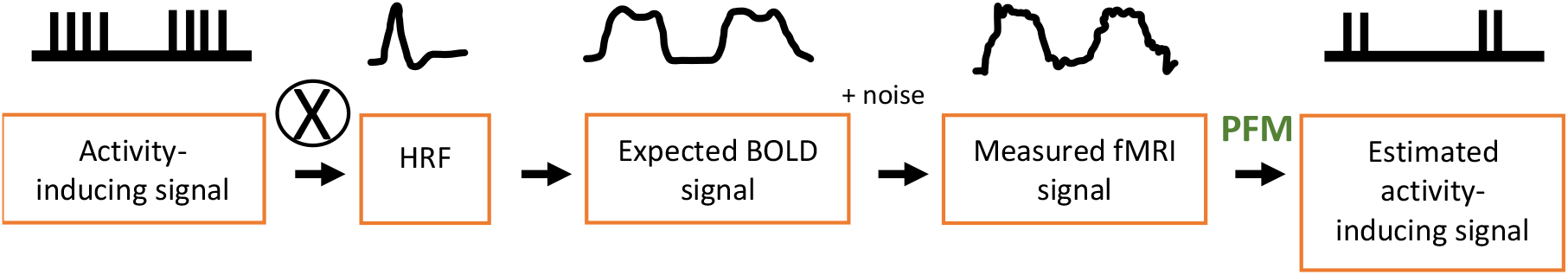
Estimation of the activation timeseries. Paradigm free mapping (PFM) involves deconvolving the measured fMRI signal to estimate the activity-inducing signal using a haemodynamic response function (HRF) template (Uruñuela et al., 2021). (Figure based on flowchart from (Uruñuela et al., 2021)).

SPSS version 27.0 was used for statistical analysis of behavioural data. Differences between blocks were calculated using paired t-tests. The behavioural blink data did not meet the assumptions for parametric testing and therefore a Wilcoxon signed-rank test was used. The level for significance was one-tailed due to the directional hypothesis that suppression blocks would result in fewer blinks. Alpha level was set to p≤0.05.

Before image analysis, the urge data were down-sampled from 10 Hz to 1 Hz and then standardised to Z-scores, through mean subtraction and division by the standard deviation. This process was completed for the random and experimental conditions for each run in each subject separately.

### 1.2.3 Temporal Relationship Between Urge and Blinks

To investigate whether urge intensity was associated with the likelihood of blink occurrence, we followed a method similar to that of Brandt and colleagues (2016). The Z-scores were calculated using the urge data from each run separately after the data were down-sampled from 10 Hz to 1 Hz. For the binary logistic regression, the urge Z-scores were concatenated across participants into separate okay-to-blink and suppress timeseries. Blink occurrence per second was binarized such that the occurrence of a blink was recorded rather than the number of blinks.

To look at the changes in urge around a blink, we extracted 5 seconds before and after each blink. The blinks for the initial 5 seconds of each block were discarded to allow the level of urge to adjust and the last 5 seconds of blinks were discarded so that the average urge around blinks would not be affected by the change in the block. These data were averaged to give a single time-series for each participant for the suppression and okay to blink blocks separately. The peak latency, skewness and excess kurtosis of these distributions were investigated using two-tailed one-sample t-tests to investigate the temporal characteristics of urge using MATLAB (MATLAB R2020a, Mathworks, Natick, MA). Where data failed tests for normality a Wilcoxon signed-rank test was used. Curvilinear regression analysis was applied using SPSS version 27.0, to investigate whether the average urge intensities (Z-score) around the blink in each condition followed a quadratic relationship.

### 1.2.4 Image Acquisition

The fMRI data were acquired using a Philips 3T Ingenia MRI scanner (Philips Healthcare, Best, The Netherlands) with a 32-channel head coil situated in the Sir Peter Mansfield Imaging Centre, Nottingham UK. FMRI data was acquired with a T2*-weighted multi-echo gradient-echo echo-planar imaging sequence with the following parameters: matrix size = 64x64; FOV = 192x192x135 mm^3^; 45 slices; in-plane resolution= 3 mm; multiband factor = 3; SENSE reduction factor P = 1.8 in right-left direction; TR = 1800 ms; TEs = 12/35/58 ms; flip angle = 80°; bandwidth = 2150.8 kHz. The functional T2* weighted scan was followed by a structural T1-weighted MP2RAGE image scan acquired using matrix size = 256x256, FOV = 192x192x135 mm, 1x1x1 mm^3^ isotropic resolution, TR = 7.1ms, TE = 3.11ms, TI = 706/3061 ms, flip angle = 80°.

### 1.2.5 Image Preprocessing

Runs with an absolute mean displacement above 1.5 mm were discarded, resulting in all 3 runs from one participant (female, 28 years old, right-handed), 2 runs from one participant, and 1 run from five participants being removed from the analysis. One further run from one participant was removed due to loss of video data meaning that blink timings could not be defined. This left a total of 52 fMRI runs. The SNR of the fMRI timeseries (tSNR) for each run was calculated using in-house scripts (MATLAB R2018b, Mathworks, Natick, MA) to assess data quality (See Appendix A of the Supplementary material).

The first echo for each fMRI run was realigned to account for head motion using MCFLIRT (FMRIB’s linear image registration tool) using the middle volume as the reference (Jenkinson et al., 2002). This same transformation was then applied to datasets the second and third echo images. Subsequently, using Tedana (version *0.0.12*), all echoes were linearly combined with weights based on the voxelwise T2* parameters (Posse et al., 1999) and this ‘optimally combined’ dataset was input to multi-echo independent component analysis (ME-ICA) with the Akaike information criterion (AIC) method being used to select the number of independent components (DuPre et al., 2021; Kundu et al., 2012a, 2013). Rica was used to visualize and manually classify any components that had been misclassified or labelled as non-classified by Tedana (Uruñuela, 2021). After that Tedana was rerun using a list of the manually accepted components for denoising purposes.

The resulting individual denoised echo datasets were then pre-processed using FSL (FMRIB software library) (Jenkinson et al., 2012). Pre-processing involved the use of a high-pass filter to remove any signals below 0.0083Hz from the fMRI data. Images were spatially smoothed using a Gaussian kernel of 5mm FWHM (full width at half maximum) to increase the SNR and to account for any major anatomical differences between subjects. Following pre-processing, activation maps for the first echo were normalised to MNI152 space and the same transformation was then applied to the second and third echoes. Finally, a nuisance regression step was applied om AFNI (Cox, 1996) to each echo dataset and the optimally combined dataset to remove physiological fluctuations and low frequency trends that were not removed by tedana. Nuisance regressors included the first four Legendre polynomials and the first five principal components of CSF voxels within the lateral ventricles, which were identified after erosion of the corresponding tissue-segmented T1-w image (Behzadi et al., 2007), and computed prior to spatial transformation to MNI152 space and spatial smoothing (Caballero-Gaudes & Reynolds, 2017).

### 1.2.6 Standard General Linear Model Analyses

For the standard image analysis, the three echoes were combined with T2* weights to generate an optimally combined dataset (Kundu et al., 2012b; Posse et al., 1999). Within the FSL GLM design matrix, three boxcar models were used to define the onset and durations of each ‘Random’, ‘Suppress’ and ‘Okay to blink’ block. Parametric regressors were defined for the standardised (Z-score) urge scores for the random baseline and experimental periods separately. An additional regressor was used to define the onset times and durations for blinks. All regressors were convolved with a double-gamma haemodynamic response function (HRF). In the first-level analysis, data from each run for each subject were analysed separately. Contrasts were set up to compare the ‘Okay to blink’ and ‘Suppress’ blocks (‘Suppress’ > ’Okay’ & ‘Okay’ > ’Suppress’) and the experimental blocks where urge was rated continuously were compared to the baseline ‘Random’ condition to account for activity related to moving of the rollerball (‘Urge’ > ’Random’ & ‘Random’ > ‘Urge’). In addition, the activity relating to blinks and urge was compared to separate the activity relating to the blink from that of high urge (‘Urge’ > ‘Blink’ & ‘Blink’ > ‘Urge’). At the second-level, results from the first-level analysis were averaged across runs for each subject. Finally, at the third-level mixed effects analysis was used to average across subjects. The results were corrected at the cluster level with a Z threshold of 3.2 (p<0.05). Regions were identified using the Harvard-Oxford cortical and subcortical structural atlases, as well as the cerebellar atlas in MNI152 space after normalization with FLIRT (FMRIB’s linear image registration tool). Conjunction analysis was used to identify whether any voxels were overlapping in the thresholded Z-statistical maps for the ‘Urge’ > ’Random’ and ‘Suppress’ > ’Okay’ contrasts.

### 1.2.7 Multi-Echo Sparse Paradigm Free mapping

Assuming a linear time invariant system, the BOLD response is assumed to be the neuronal signal convolved with the HRF (+ noise) (Poldrack et al., 2011). PFM works by deconvolving the fMRI signal using the HRF via regularised least-squares estimation to estimate the neuronal-related signal at each voxel (Figure 2) (Caballero Gaudes et al., 2013; Uruñuela et al., 2020). In this work, a version of PFM (MESPFM) modified to take account of the linear dependence of the BOLD response with the echo times of the additional signals available from multi echo fMRI data (Caballero-Gaudes et al., 2019).

The MESPFM analysis was run using the 3dMEPFM command implemented in AFNI (Cox, 1996; Cox & Hyde, 1997). The signal percentage change for each echo was calculated by dividing the detrended data by the mean of the voxel data on a voxel-by-voxel basis. Prior to the analysis with MESPFM, the data relating to the random baseline at the beginning and end of each run were removed, so that only the six 1-minute blocks of alternating blink suppression and rest remained.

For MESPFM, the regularization parameter was selected using the Bayesian Information Criterion (BIC) (Caballero-Gaudes et al., 2019) according to the goodness of fit of the estimated model. Specifically, BIC will introduce an increasing penalty for more events being included in the model to prevent overfitting (Dziak et al., 2020). The HRF used for the deconvolution was the SPM canonical HRF (Penny et al., 2007), and the model only considered changes in the transverse relaxation rate (R2*).

A surrogate dataset was created by shuffling the data from the six 1-minute blocks of alternating blink suppression and rest, before the signal percentage change was calculated. This created a new dataset with the same temporal (and spectral) distribution as the original dataset but without the temporal relationships between the timepoints, which could act as a null distribution. This shuffled dataset was analyzed with the same MESPFM algorithm as the original dataset. If an activation event detected by MESPFM in the original dataset exhibits a larger amplitude than those seen in the surrogate dataset, then it is unlikely to have happened by chance and can be considered significant. This threshold was defined as the median amplitude of the surrogate activation timeseries for that run.

### 1.2.8 Activation Time Series

Following MESPFM, we removed the activity of spurious voxels via spatiotemporal clustering using a sliding window approach. The sliding window consisted of 3 datapoints: the current datapoint and those either side. The current data point at each voxel was then substituted as the value of the largest absolute value within that window. The *3dmerge -1clust* AFNI (Cox, 1996; Cox & Hyde, 1997) function was used to cluster neighbouring voxels, with a minimum cluster size of 10. The spatiotemporal clustering mask was then applied to the original data to remove spurious, isolated activations that are likely false activations.

Then, an activation time series (ATS) (Gaudes et al., 2011) was computed by counting the number of voxels with a negative estimated R2* signal (i.e., a positive BOLD response) at each TR in our selected region of interest (ROI), the right insula.

The right insula was selected as our ROI due to its frequent identification in studies exploring the neural correlates of urge and its hypothesized role in the urge-to- act (Berman et al., 2012; Jackson et al., 2011; Lerner et al., 2009; Mazzone et al., 2010; Nahab et al., 2009; Yoon et al., 2005). If there is contribution from different subregions within the right insula then we may be able to tease these subregions and any separable co-activations apart during clustering, following the MESPFM. Whilst the MCC is commonly identified as a region involved in the urge- to-act, its hypothesized role in the selection of an action in response to urge rather than in the urge sensation itself means it would be a less ideal candidate for ROI analysis (Jackson et al., 2011).

The mask of the right insula was created based on insula parcels from the Schaefer 1000 parcels 17 network atlas (Schaefer et al., 2018). Finally, we selected those peaks that had a higher number of activated voxels within the ROI compared with the shuffled dataset as any peaks higher than this are unlikely to have happened by chance.

### 1.2.9 K-Means Clustering

Clustering was used to identify any patterns in the activation maps associated with the selected ATS peaks. The input was the matrix of pairwise distances between the activation maps associated with the selected ATS peaks. The metric used for calculating these pairwise distances was the Euclidean distance. This would help us to group together coactive regions. K-means clustering aims to separate the data into k clusters, here k was chosen using consensus clustering (Wu et al., 2015). The selected ATS maps would be assigned to the cluster that minimised the distance between the data points and their cluster centroids.

For the consensus clustering, k-means clustering was applied to 80% of the data with k values in the range 2 to 15 with 100 iterations per k. The k with the highest consensus value was selected. The consensus value is the average proportion of times that any pair of data points were assigned to the same cluster across the runs, giving a value between 0 and 1.

The K-means algorithm was run 50 times with different centroid seeds with the number of clusters determined by the consensus clustering. Finally, the voxelwise Z-scores for the activation maps for each cluster were calculated using Z-normalization in space (i.e. subtracting the mean of the ΔR2* (change in 1/T2*) values across the brain and dividing by the corresponding standard deviation).

We then compared the MEPFM cluster maps with the urge, suppression and blink GLM-based maps to identify which they most closely represented. To do this, the Z-score maps of the identified MESPFM clusters were multiplied by -1 to account for the fact that the MESPFM estimates changes in R2* rather than the BOLD signal since negative changes in R2* generate a positive BOLD response, and vice versa. Next the MESPFM cluster Z-score maps were thresholded at Z = 3.2 to make them comparable to the GLM-based maps (‘Suppress’ > ‘Okay to blink’, ‘Blinks’, ‘Urge’ > ‘Random’). Then, conjunction analysis was used to identify whether any voxels were overlapping. The highest overlap between the GLM-based masks and the MESPFM-cluster mask was used to determine which GLM-based map the cluster represented the most. The percentage of overlapping voxels within the GLM-based masks is reported.

## 1.3 Behavioural Results

All blinks in each run were first annotated by one rater, then a random 60-second block from each run was annotated by a second rater using ELAN (MH, IM, KD), with an average agreement of 95.51% ± 10.13 (mean ± SD) (*ELAN*, 2019). Any blink discrepancies were discussed until agreement was achieved for all blink occurrences. The average number of blinks per minute in the ‘Okay to blink’ condition was 31.20 ± 3.63 (mean ± standard error of the mean (SEM)) while in the suppression condition this was significantly lower with 5.12 ± 0.81 blinks per minute (*t*(20) = -4.249, p<0.001). The average urge per minute was 22.79% ± 4.00 and 55.62% ± 3.42 for ‘Okay to blink’ and ‘Suppress’ blocks, respectively. The difference between the urge in the two conditions was highly significant (*t*(20) = -10.901, p<0.001). These findings indicate that participants successfully followed instructions to suppress blinks and that this was associated with an increased urge-to-blink.

Figure 4 shows examples of runs from two different representative subjects, where urge is shown to rise during the period of suppression, and suddenly decrease after ‘escape’ blinks. However, while for some subjects urge flattened throughout periods where blinking was okay (Figure 4B), others reported small increases in urge before the blinks (Figure 4A), although the magnitude this reached before a blink was released was lower than that seen in the suppression blocks.

**Figure 3.**
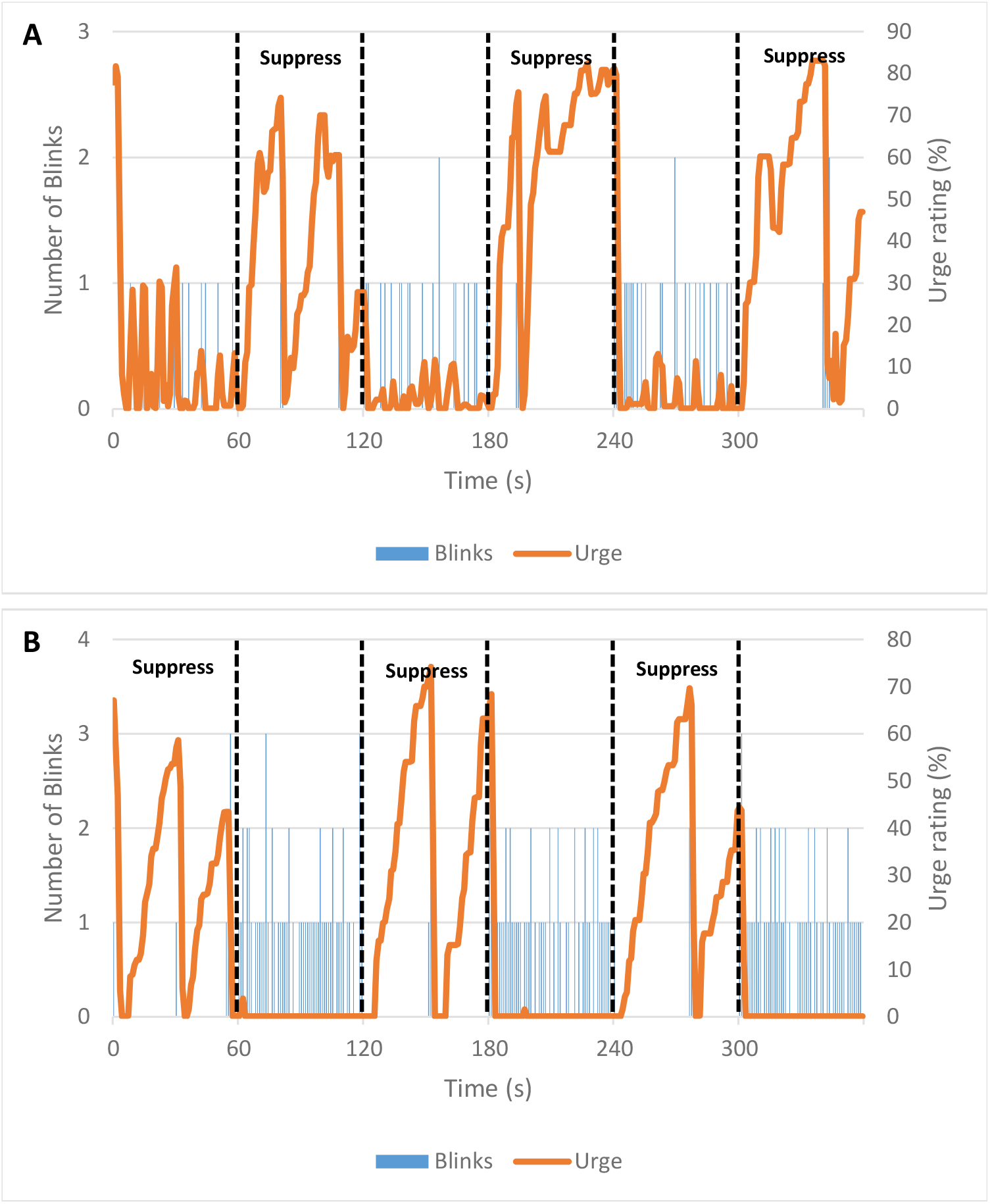
The association between the urge-to-blink and blinking. Graphs displaying blink timings for individual task runs from two representative participants alongside their subjective urge rating across time. Panel A) shows that some participants felt increases in urge even in ‘Okay to Blink’ blocks, whereas panel B) shows that some participants only felt urge during suppression blocks.

**Figure 4.**
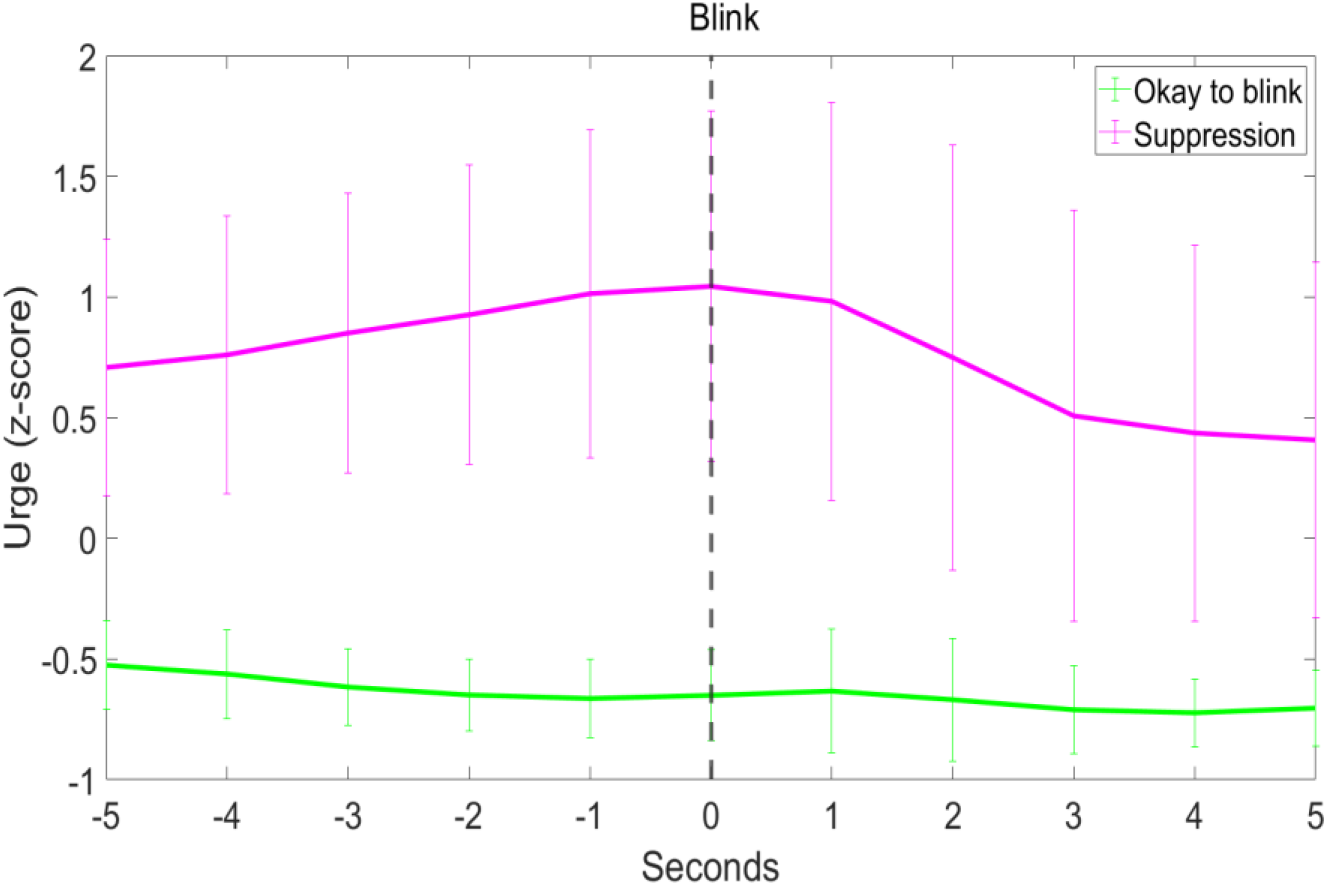
The distribution of mean urge per second around a blink at time 0. Error bars show the standard deviation.

### 1.3.1 Temporal Relationship Between Urge and Blinks

A binary logistic regression showed that only 0.6% of the variance in blink occurrence during ‘Okay to blink’ could be explained by changes in subjective urge ratings (Cox & Snell R^2^ = 0.006, χ^2^(1) = 53.667, p < 0.001; Exp(B) = 0.806, Wald(1) = 107.279, p < 0.001). Due to the scarcity of blinks in the ‘Suppress’ condition, all instances of blinks were classified as outliers by the model and so the data were not appropriate for this type of analysis.

A curvilinear regression showed that the mean urge around blinks followed a significant quadratic distribution over time in both the ‘Okay to blink’ (F(2,8) = 27.279, p < 0.001, Adjusted R^2^ = 0.840; Estimated urge = -0.661 - 0.017 * (time to blink) + 0.001 * (time to blink)^2^) and ‘Suppress’ conditions (F(2,8) = 26.192, p < 0.001, Adjusted R^2^ = 0.834; Estimated urge = 0.948 - 0.038 * (time to blink) – 0.019 * (time to blink)^2^) (Figure 5).

**Figure 5.**
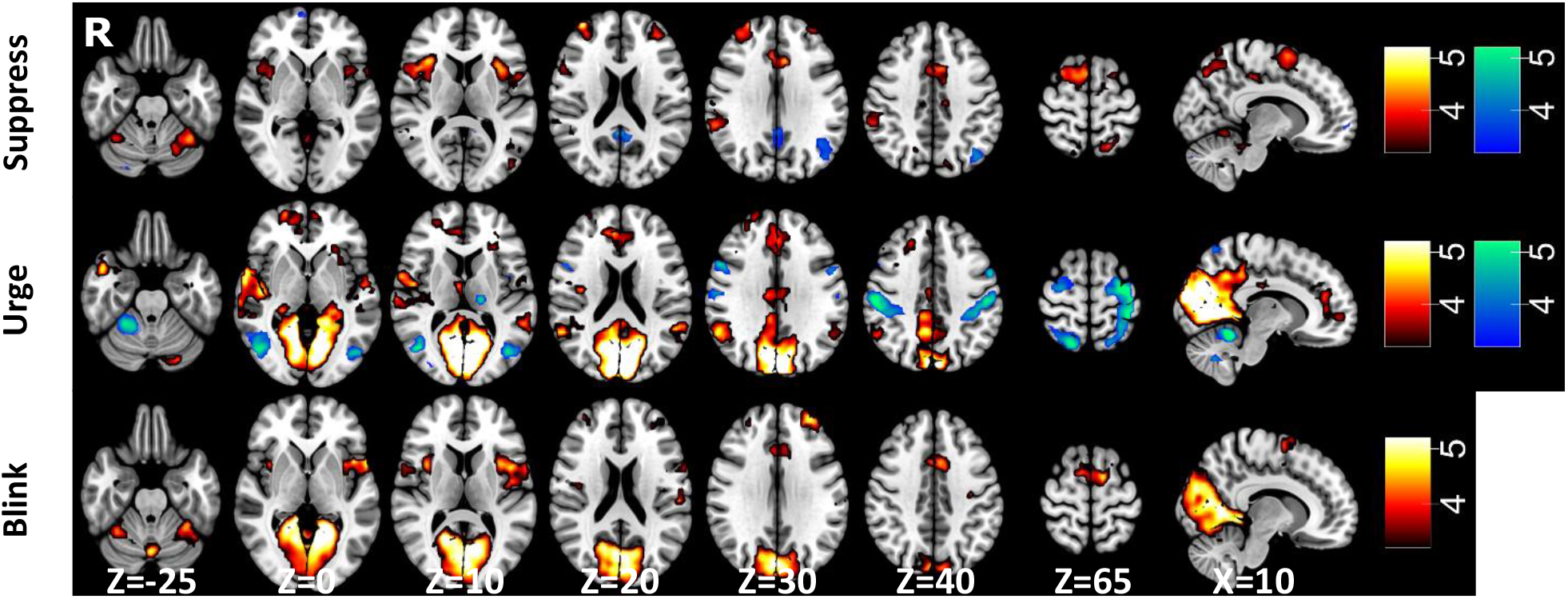
BOLD response associated with blink suppression, urge-to-blink and blinking. Statistical maps overlaid onto the MNI152 brain showing significant activations for the (top) ‘Suppress’ > ’Okay’ (red), ‘Okay’ > ’Suppress’ (blue); (middle) ‘Urge’ > ’Random’ (red), ‘Random’ > ‘Urge’ (blue); (bottom) ‘Blink’ > ‘Urge’; (red) contrast. Statistical maps were thresholded at Z=3.2 (p<0.05).

In the ‘Okay to blink’ condition, urge intensity peaked significantly before the blink (-3.55 s ± 2.52 (mean ± sd), z(19) = -3.68, p < 0.001), whereas in the ‘Suppress’ condition urge peaked at blink onset (0.56 s ± 2.87, z(17) = 0.89, p > 0.05) (Figure 5). There was no significant skew in the suppression condition (0.01 ± 0.75, t(17) = 0.06, p > 0.05), whereas in the free blinking condition, urges were slower to decrease than they were to increase before the peak (0.78 ± 0.62, z(19) = 3.58, p < 0.001). While there was no significant kurtosis in the ‘Okay to blink’ condition (2.84 ± 0.98, z(19) = -1.57, p > 0.05), the distribution of urge around the blink in the suppression condition was broader than that of a normal distribution (2.16 ± 1.27, z(17) = -2.90, p < 0.01). Two subjects were not included in the curvilinear regression and the temporal characteristics analysis for the ‘Suppress’ condition due to having no ‘escape’ blinks.

## 1.4 Standard General Linear Model Results

Locations of clusters local maxima for all GLM comparisons are defined within Appendix B of the Supplementary Material.

### 1.4.1 Block Analysis

For the contrast of ‘Suppress’ > ‘Okay to blink’, significant activations were identified with peaks in the dorsolateral prefrontal cortex (DLPFC), lateral occipital cortex, cerebellum, opercular cortices, supramarginal gyrus (SMG) and posterior cingulate (PCC) (Figure 6). Notably, significant activations were found in the left primary somatosensory cortex, MCC, supplementary motor area (SMA) and bilateral insulae. When contrasting ‘Okay to blink’ > ‘Suppress’, clusters were identified in the frontal orbital cortex, lateral occipital cortex, PCC, middle frontal gyrus and a small area in the cerebellum (Figure 6).

**Figure 6.**
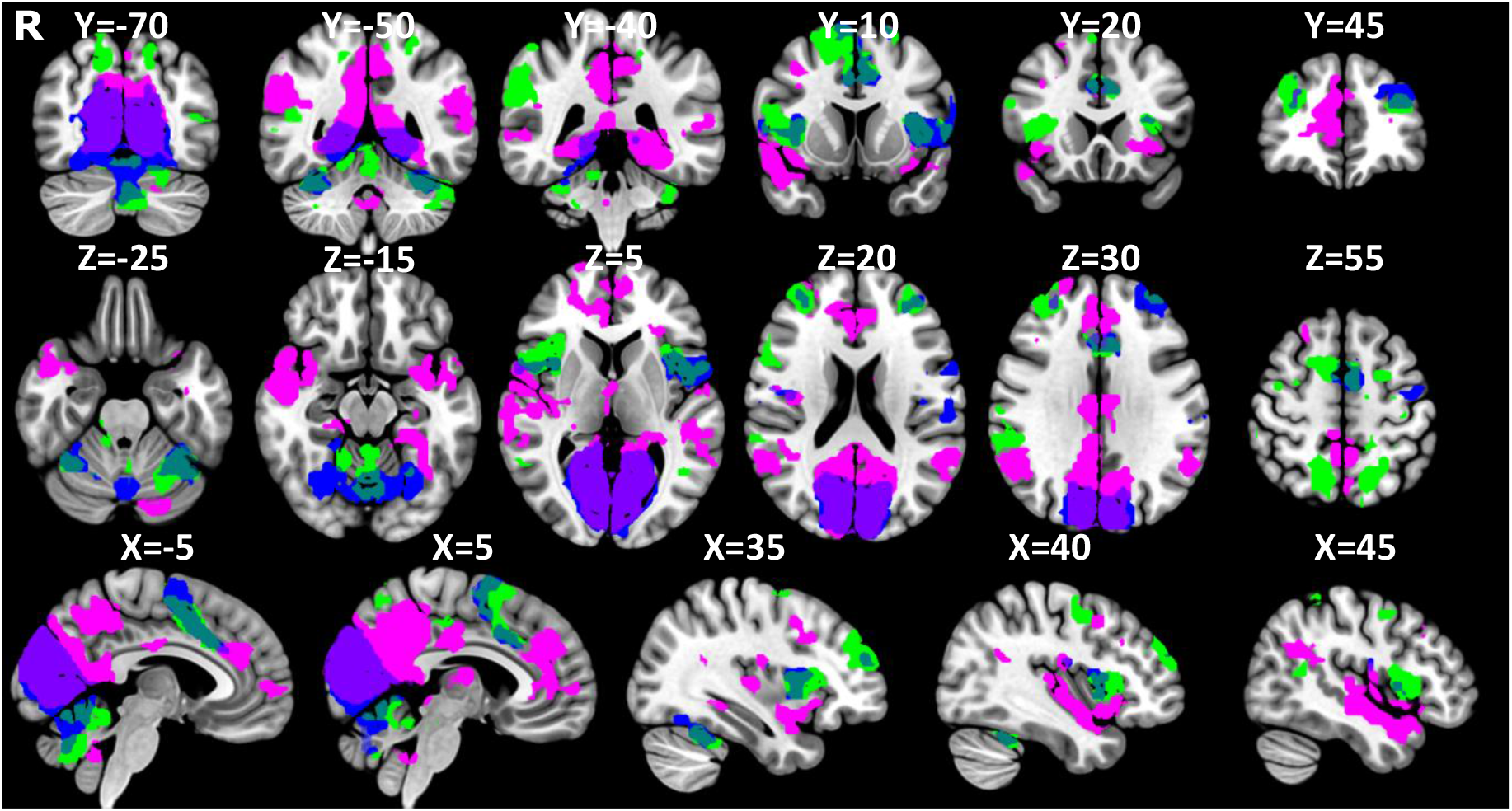
Separate networks for urge-to-act and action suppression. Masks of significant activation for the ‘Suppress’ > ‘Okay’ (green), ‘Urge’ > ‘Random’ (pink) and ‘Blink’ > ‘Urge’ (blue) contrasts overlaid onto the MNI152 brain.

### 1.4.2 Urge Analysis

For the contrast of ‘Urge’ > ‘Random’, significant activations were identified in the medial occipital cortex, opercular cortex, ACC, bilateral insulae and cerebellum (Figure 6). When contrasting ‘Random’ > ‘Urge’, clusters were identified in the bilateral sensorimotor cortices, lateral occipital cortex, cerebellum, left thalamus, opercular cortex and insulae (Figure 6).

In Figure 7, the activations associated with the contrast ‘Urge’ > ‘Random’ are visualised alongside those associated with ‘Suppress’ > ‘Okay to blink’ and ‘Blink’ > ‘Urge’ showing an overlap between blinking and suppression in the MCC and SMA, while the anterior cingulate cortex (ACC) is associated with the urge-to-blink. Notably, there is a differentiation in insula involvement with a dorsal-anterior portion involved in suppression and blinking, a central portion involvement in blinking and posterior and ventral-anterior regions being active during feelings of urge-to-blink (Figure 7).

**Figure 7.**
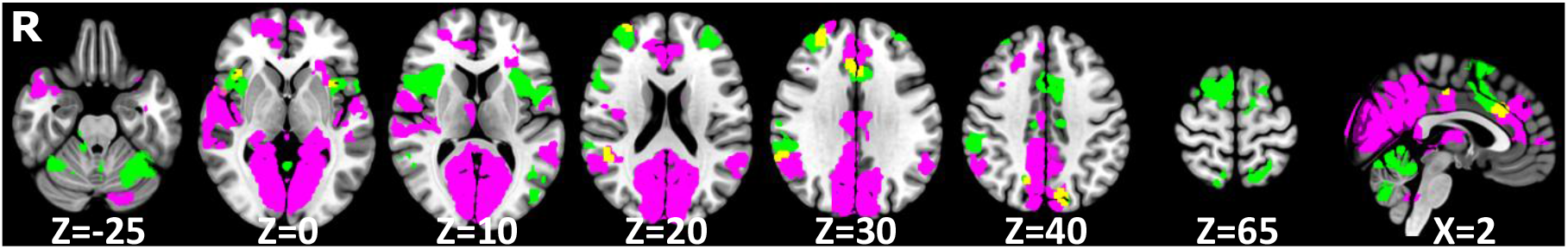
Voxels active during both the urge-to-blink and suppression. Masks of significant activation for the ‘Suppress’ > ‘Okay’ (green) and ‘Urge’ > ‘Random’ (pink) contrasts with overlapping voxels in yellow, overlaid onto the MNI152 brain.

### 1.4.3 Blink Analysis

For the contrast ‘Blinks’ > ‘Urge’, there were significant activations in the medial occipital cortex, MCC, opercular cortex, insulae, DLPFC, SMA and left primary sensorimotor cortex (Figure 6). No regions were identified by the ‘Urge’ > ‘Blinks’ contrast.

### 1.4.4 Conjunction Analysis

Figure 8 shows the overlap between the significant activations in the ‘Urge’ > ‘Random’ and ‘Suppress’ > ‘Okay to blink’ contrasts. Voxels were identified in the MCC, right DLPFC, right superior SMG, right angular gyrus, left postcentral gyrus, bilateral anterior insulae, right opercular cortex, right precuneous, left lateral occipital cortex and left VI in the cerebellum.

**Figure 8.**
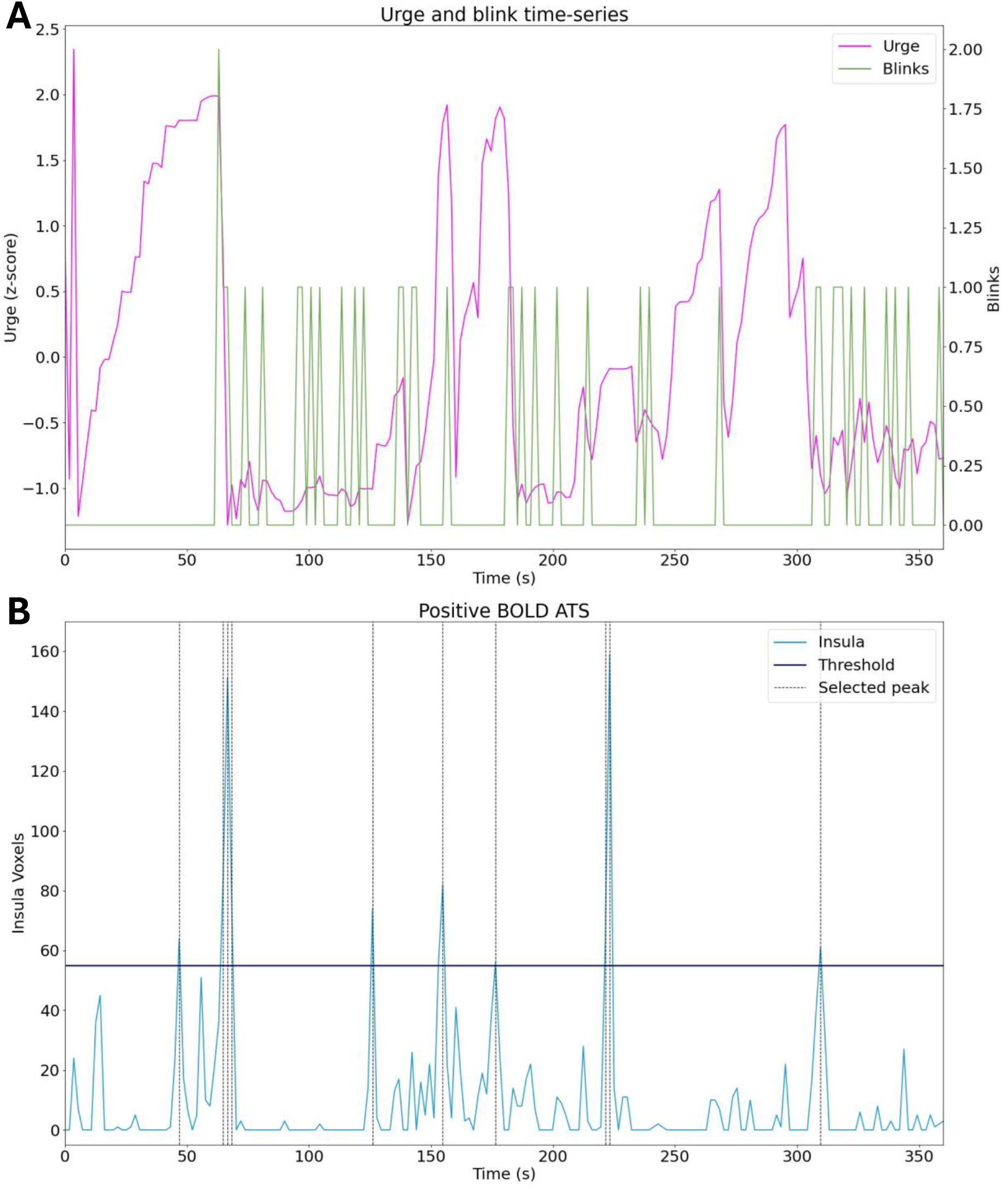
The activation timeseries from a representative subject. A) The interpolated urge scores and blink frequencies per TR.; B) All positive BOLD (negative R2*) activations within the right insula with the threshold set by the shuffled dataset.

## 1.5 Multi-Echo Sparse Paradigm Free Mapping Results

Figure 9 illustrates the detection of BOLD activation within the right insula obtained in a single run from a representative subject. Figure 9A shows the interpolated urge scores and blink frequencies per TR. Activation peaks within the right insula (selected ROI) which surpass the threshold are shown in Figure 9B. It is worth noting that not all the runs showed right insula activation surpassing the threshold set by the shuffled dataset (See Appendix C of the Supplementary Material).

**Figure 9.**
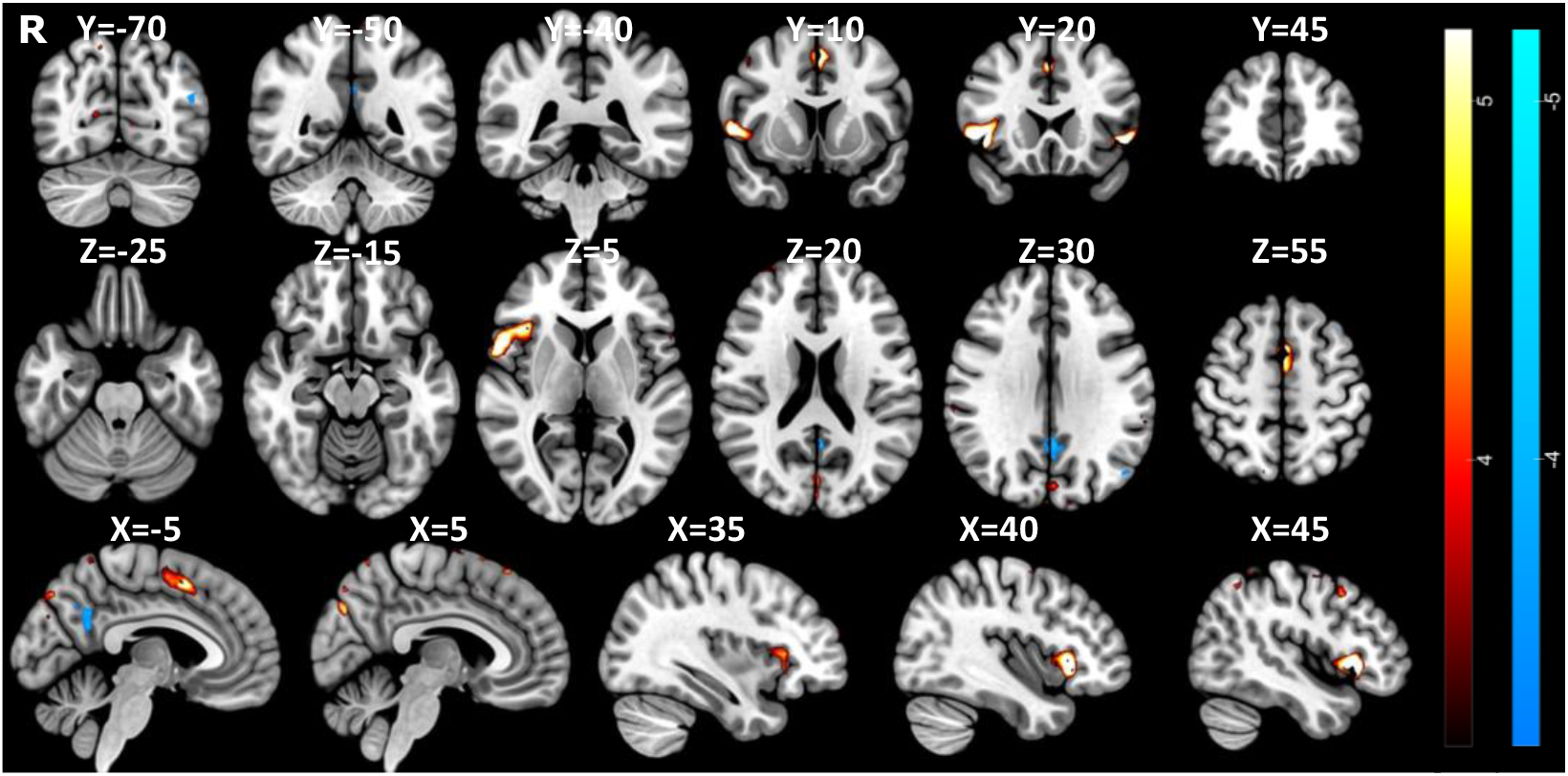
Cluster map 1 identified using multi-echo sparse paradigm free mapping. Positive activation was identified in the SMA, paracingulate cortex, ACC, bilateral insulae, and both medial and lateral occipital areas. Statistical maps were thresholded at Z= ±3.2 (p<0.05).

Consensus clustering determined that 3 clusters gave the most stable solution at the group level with a consensus value of 0.60140. The thresholded K-means cluster maps (k = 3) are shown in Figure 10, Figure 11 and Figure 12 (Z=±3.2). The positive, binarized K-means output maps are shown in Figure 13 (Z= 3.2).

**Figure 10.**
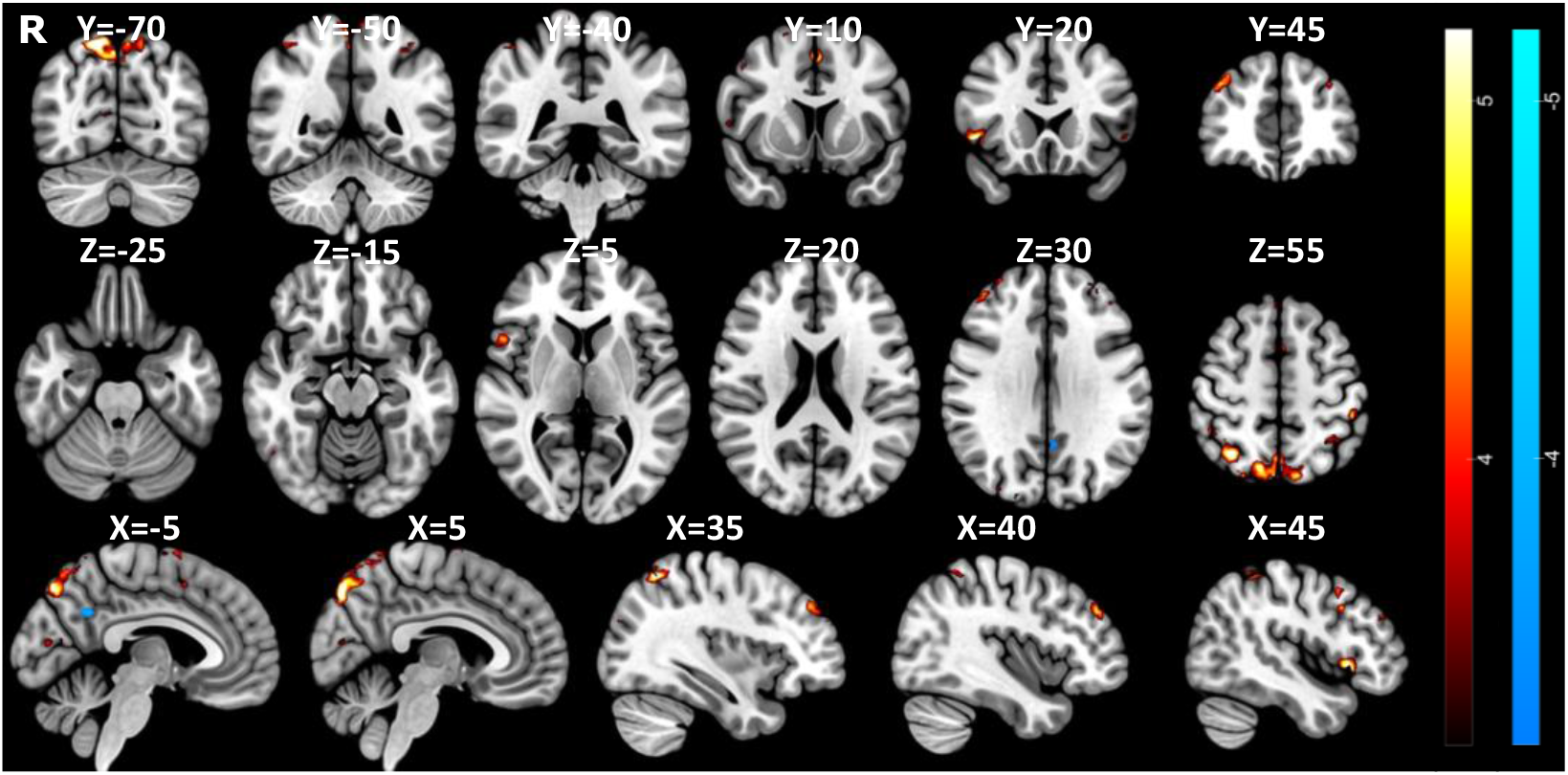
Cluster map 2 identified using multi-echo sparse paradigm free mapping. Positive activation was identified in the SMA, paracingulate cortex, right insula, bilateral DLPFC, and both medial and lateral occipital areas. Statistical maps were thresholded at Z= ±3.2 (p<0.05).

**Figure 11.**
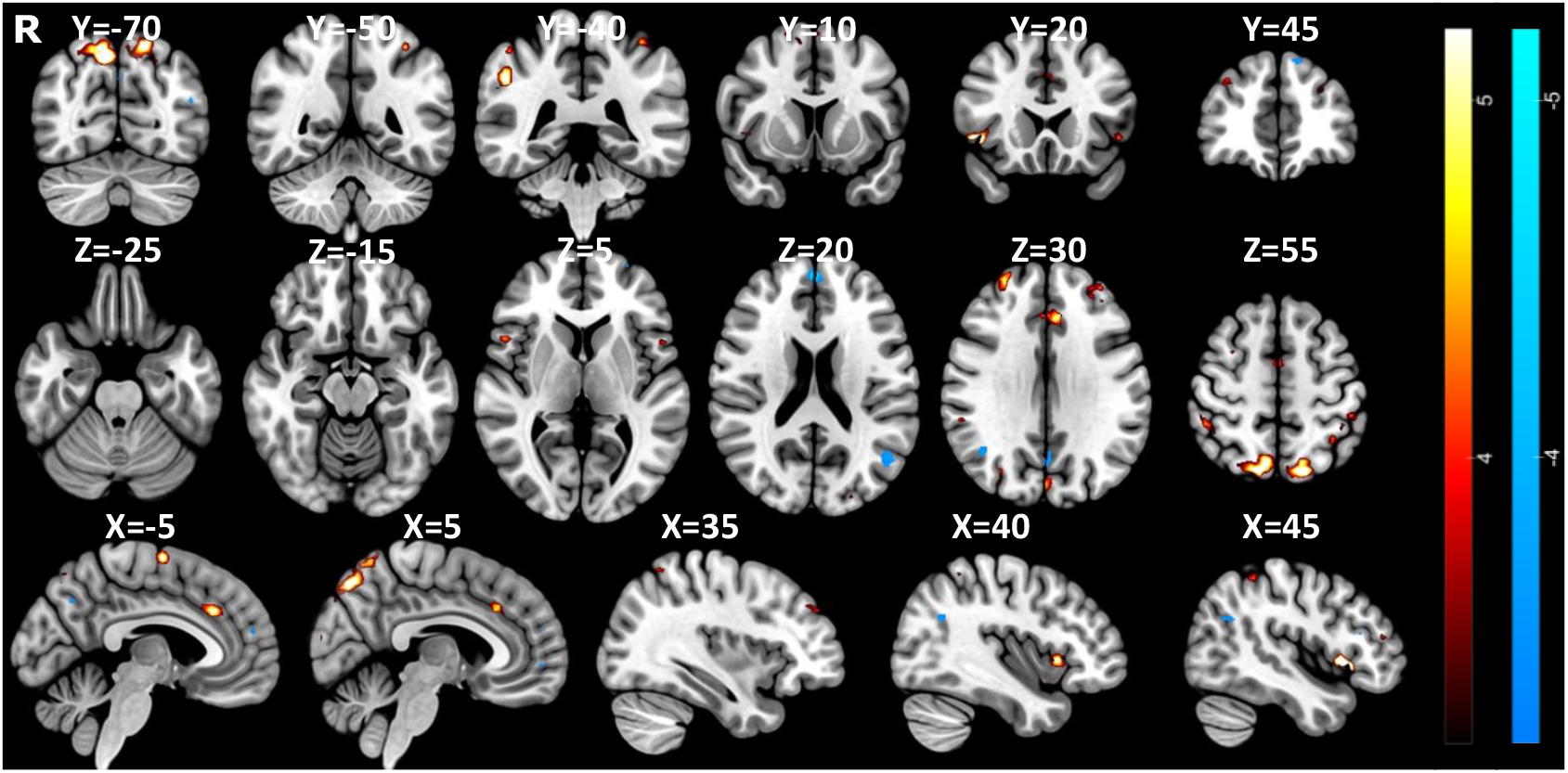
Cluster map 3 identified using multi-echo sparse paradigm free mapping. Positive activation was identified in the SMA, paracingulate cortex, ACC, right insula, bilateral DLPFC, and both medial and lateral occipital regions. Statistical maps were thresholded at Z= ±3.2 (p<0.05).

**Figure 12.**
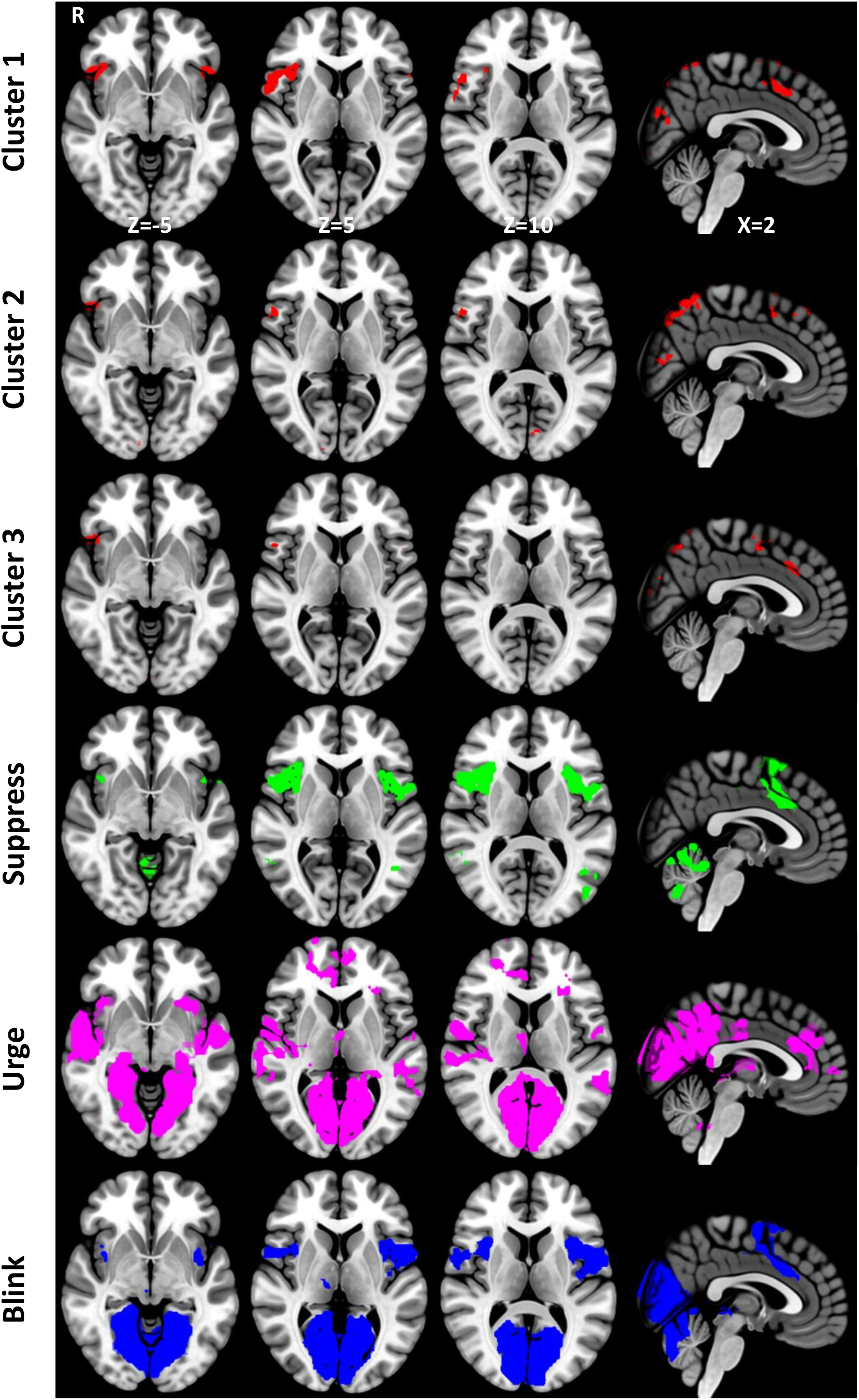
Comparison of the masks generated during the multi-echo sparse paradigm free mapping (Clusters 1-3) and the conventional general linear model analysis (Suppression, Urge, Blink) (thresholded at Z = 3.2).

The thresholded activation map for Cluster 1 reveals significant positive activation in the SMA, paracingulate cortex, ACC, bilateral insulae, bilateral frontal opercular cortices, right IFG pars opercularis, bilateral frontal orbital cortices, right postcentral gyrus, right superior parietal lobule, and both medial and lateral occipital areas. Significant negative activation localised to the left medial frontal gyrus (MFG), left lateral occipital cortex, precuneous and the PCC. Similarly, Cluster 2 involves positive activation of the SMA, paracingulate cortex, right insula, right frontal opercular cortex, bilateral IFG pars opercularis, right frontal orbital cortex, bilateral superior frontal gyri (SFG), right MFG, bilateral DLPFC, left postcentral gyrus, bilateral superior parietal lobules, and both medial and lateral occipital areas. Negative activation was seen within the left lateral occipital cortex, precuneous and PCC. Finally, Cluster 3 shows positive activation in the SMA, paracingulate cortex, ACC, right insula, bilateral frontal opercular cortices, bilateral SFG, bilateral DLPFC, left sensorimotor cortex, bilateral superior parietal lobule, and both medial and lateral occipital regions. Negative activation was seen within the precuneous, PCC, left lateral occipital cortex, left prefrontal gyrus and the SFG.

The three MESPFM cluster maps show positive activation within the right dorsal-anterior insula, paracingulate cortex, SMA, and medial and lateral occipital cortices (Figure 13). Figure 13 illustrates the results from both the MESPFM analysis and the conventional GLM analysis. The largest overlap between the three thresholded MESPFM cluster maps was with the regions shown to be active during suppression (Table 1), where the overlap was defined as the percentage of overlapping voxels within the GLM-based masks.

**Table 1.**
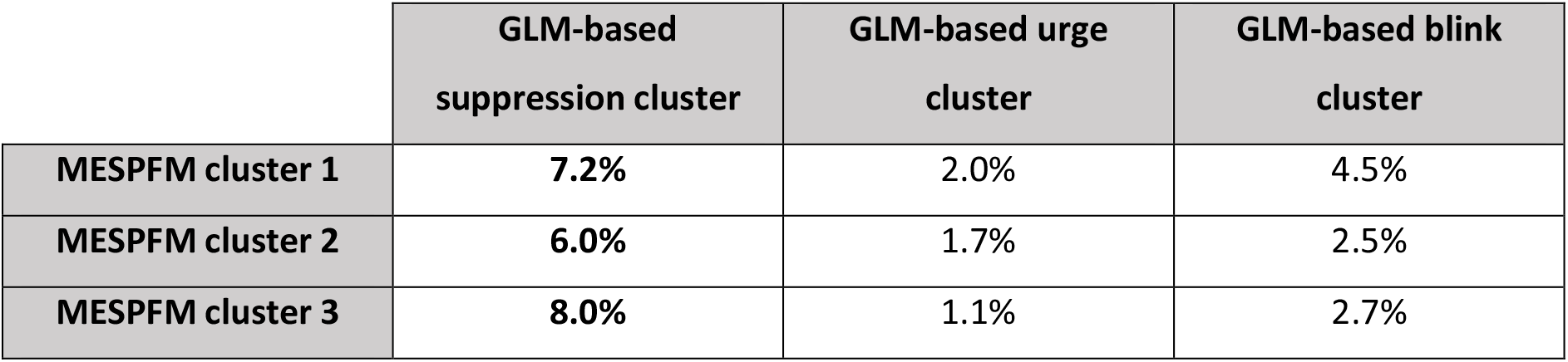
Percentage overlaps of the MESPFM-cluster masks with the GLM-based cluster masks (Chapter 3). The largest overlap for each MESPFM cluster is highlighted in bold.

|  | GLM-based<br>suppression cluster | GLM-based urge<br>cluster | GLM-based blink<br>cluster |
| --- | --- | --- | --- |
| MESPfM cluster 1 | **7.2%** | 2.0% | 4.5% |
| MESPfM cluster 2 | **6.0%** | 1.7% | 2.5% |
| MESPfM cluster 3 | **8.0%** | 1.1% | 2.7% |

## 1.6 Discussion

This fMRI study investigated the urge-to-blink using both a conventional general linear model analysis with a parametric model of subjective urge ratings and a MESPFM approach. The aim was to disentangle the anatomical correlates of the urge-to-blink from those of action suppression and to validate whether MESPFM can be used to identify neuronal activity in an action suppression paradigm without prior specification of urge timecourses.

### 1.6.1 Behavioural relationships between urge and blinks

Previous attempts to model the urge-to-blink have either employed a sawtooth model (Berman et al., 2012), where urge builds up linearly across the suppression block before decreasing at the end of the block, or an event-related model (Botteron et al., 2019), where urge decreases following escape blinks in the suppression block. Here, the representative examples of continuous urge ratings during the task show that blinking, particularly during suppression blocks, causes a temporary decrease in urge intensity. Therefore, although sawtooth models are likely better at approximating urge compared to a block analysis (Berman et al., 2012), they are still too simplistic as they do not capture the complex temporal characteristics of the urge, e.g. they do not consider escape blinks during suppression. More recent models that take account of these ‘escape’ blinks, such as the event-related approach suggested by Botteron and colleagues, more accurately represent real-time urge ratings (Botteron et al., 2019). If applied to fMRI data, the model could theoretically identify neural correlates of the urge-to-blink relatively well. However, this approach would not be appropriate in the analysis of the urge-to-tic where overt expression of the behaviour would be suppressed during scanning, highlighting the need for continuous urge rating or alternative modelling and analysis approaches.

Results from the curvilinear regression demonstrated a quadratic relationship between urge and blinks (see Figure 5) indicating that urge increases during suppression but diminishes after the blink. This is further supported by the urge peaking at blink onset. While the ‘Okay to blink’ blocks also showed a significant quadratic relationship, urge did not peak at blink onset. Furthermore, blink occurrence could not be predicted by the urge score in ‘Okay to blink’ blocks. This suggests that in the case of blinking in healthy participants, urge arises due to the act of suppression. Brandt and colleagues (2016) also found a significant quadratic distribution of urge and that the peak in urge was coincident with blinks in both the free to blink and suppress conditions.

### 1.6.2 Neural Correlates of the Urge-to-Blink

The regions identified using the urge parametric model included the insulae and ACC. These regions are commonly implicated in studies of urge; therefore, the right insula and cingulate cortex are thought to be key nodes in the urge network (Jackson et al., 2011).

Activation of the insula has been linked to various urge sensations, such as those related to ticcing (Bohlhalter et al., 2006; Neuner et al., 2014), blinking (Abi-Jaoude et al., 2018; Berman et al., 2012; Lerner et al., 2009) and yawning (Jackson et al., 2011). Patients with obsessive compulsive disorder (OCD) show increased insula activity during early blink suppression compared to controls (Stern et al., 2020). Furthermore, PU severity has shown a negative association with the volume of left insular grey matter thickness in TS patients (Draper et al., 2016).

Subregions of the insula are thought to have differing functions (Kelly et al., 2012; Kurth et al., 2010). The posterior insula has a role in the initial processing of both noxious and non-noxious somatosensory stimuli (Ostrowsky et al., 2002), whereas the anterior insula integrates information from several functional systems to bring about interoceptive awareness (Craig, 2009; Kurth et al., 2010). In agreement with this concept of a functional division, our data suggest that the posterior insula is involved in the processing of urge sensations as has been theorised previously (Tinaz et al., 2015). Information is thought to flow in a hierarchical fashion from the posterior insula to the anterior insula, with initial sensory processing in the posterior portion and progressive integration of information in the anterior portion to give a final representation that incorporates all the task information (Craig, 2009; Craig et al., 2000). Here, the ventral-anterior insula was also associated with urge, and this subregion has been shown to be linked with emotional processing (Kelly et al., 2012; Kurth et al., 2010). Similarly, stimulation of the pregenual ACC has been shown to induce emotional, interoceptive and autonomic experiences (Caruana et al., 2018). Previous analyses of the functional connectivity of the insula have indicated that the ventral-anterior subregion is connected to the rostral ACC within a limbic network that is associated with emotional salience detection (Cauda et al., 2011). On the other hand, the posterior insula is connected to sensorimotor regions within a network involved in response selection (Cauda et al., 2011). Therefore, it is possible that somatosensory urges are processed by the posterior insula, and through integration of information in the ventral-anterior insula and ACC, these urges become emotionally salient, which perhaps draws attention to their uncomfortable nature. Meanwhile, functional connections between the posterior insula and sensorimotor regions, including the MCC and SMA, may lead to either the continuation of suppression or to the release of a blink in response to the urge sensation.

Along with the previously described regions, the medial occipital cortex was also shown to be involved in feelings of urge and during blinks. We conjecture that this activation is specific to the urge-to-blink rather than the general urge network. Activation of the occipital cortex has been seen in previous studies looking at the urge-to-blink (Berman et al., 2012; Stern et al., 2020; Yoon et al., 2005), but it has not been described in relation to other forms of the urge-to-act (Bohlhalter et al., 2006; Jackson et al., 2011). This activation could be due to a loss of visual input during blinks (Nakano et al., 2013). However, as activation of this region is also seen when blinking in the dark (Golan et al., 2018), we suggest that there might be a combined effect of the medial occipital cortex receiving motor efferents when a blink is likely to occur, for instance, when the urge-to-blink is high (Bristow et al., 2005).

We assumed that the regions which showed greater activity in the ‘Random’ > ‘Urge’ contrast were associated purely with the movement of the trackball device. As such, this was used as an active baseline to tease apart activity related to urge from that of movement. However, participants moved the trackball more during the random condition than they did during the experimental blocks, and as such, this active baseline was not perfect. The higher activity seen in the cortical and cerebellar (lobules I-VI and VIII (Guell et al., 2018)) sensorimotor regions in the ‘Random’ > ‘Urge’ contrast was likely due to this increased movement of the trackball.

### 1.6.3 Neural Correlates of Action Suppression

A meta-analysis looking at the neural correlates of response inhibition identified the IFG (pars opercularis), SMG, SMA, MCC and bilateral insulae amongst other regions involved in action suppression (Zhang et al., 2017). These regions were also found to be active in our ‘Suppress’ > ‘Okay’ contrast, and the network bears a striking resemblance to the executive control network (Beckmann et al., 2005). The ‘Suppress’ > ‘Okay’ contrast also identified the dorsolateral PFC, which is thought to be involved in cognitive control (Miller & Cohen, 2001) and has previously been shown to be active to a higher degree in TS patients compared to healthy controls during blink inhibition (Mazzone et al., 2010). Therefore, this area may coordinate regions in a top-down manner to achieve the goal of blink suppression (Miller & Cohen, 2001). We also see that the activation of the insula/operculum extends into the IFG (pars opercularis), which is not surprising given its central role in the motor response inhibition network (Aron et al., 2004, 2014). More recently, Abi-Jaoude and colleagues found that the left DLPFC and left IFG showed higher activity in participants with fewer ‘escape’ blinks suggesting the regions play a role in successful suppression (Abi-Jaoude et al., 2018).

In addition, the cerebellum is hypothesised to have a complementary role in motor inhibition (Picazio & Koch, 2015). A transcranial magnetic stimulation study showed that a conditioning pulse to the right lateral cerebellum 5-7 ms prior to electrical stimulation of the left motor cortex resulted in a decrease in motor evoked potential amplitude (Ugawa et al., 1995). On the other hand, the higher cerebellar activity in lobules I-VI and VIII during suppression could be due to more variation in the urges being reported during these blocks, in comparison to when blinking was okay, meaning more hand movement was required to rate them (Guell et al., 2018).

As previously mentioned, the anterior insula is involved in multimodal integration and salience (Craig, 2009; Kurth et al., 2010). The activation seen during suppression was in the dorsal-anterior segment, which has been associated with cognitive processing (Kelly et al., 2012; Kurth et al., 2010). Notably, in a meta-analysis by Kurth and colleagues the dorsal-anterior region was the site which was commonly active across task modalities except sensorimotor tasks (Kurth et al., 2010). Therefore, it may be that suppression of an action involves integration of task information so that the automatic response to blink during periods of increased discomfort can be inhibited in blocks of suppression.

The insula and ACC (which includes the MCC in older descriptions) are theorised to be the limbic sensory and motor regions, respectively (Craig, 2009; Craig et al., 2000) and are commonly co-active in studies of urge (Abi-Jaoude et al., 2018; Berman et al., 2012; Bohlhalter et al., 2006; Jackson et al., 2011; Lerner et al., 2009; Mazzone et al., 2010). The MCC has previously been suggested to have a role in selecting an action in response to urge sensations, as intra-cortical stimulation of the MCC induces complex motor responses (Caruana et al., 2018; Jackson et al., 2011). Movement can also be evoked through stimulation of the SMA (Fried et al., 1991), and in some cases, it also induces feelings of urge, which may explain why its activation has frequently been associated with blink suppression (Berman et al., 2012; Lerner et al., 2009). As both the MCC and SMA were active during blinks as well as suppression blocks, these nodes may decide whether to release suppressed behaviours in response to feelings of urge. Similarly, blinks in suppression blocks may involve more influence from these pre- motor regions (Berman et al., 2012). This could be investigated in the future through a comparison of blinks in suppress and free to blink conditions. Alternatively, activation of these regions during ‘Suppress’ blocks could relate to the effort participants exert to keep their eyes open (Lerner et al., 2009).

### 1.6.4 Neural Correlates of Blinking

Insula activation during blinks was restricted to the dorsal anterior insula and the mid-insula. As previously mentioned, the dorsal anterior activation might be linked with task-related integration of information, such as whether blinking was ‘allowed’ during the task block (Kurth et al., 2010). We hypothesise that the mid-insula activation is linked to the movement and sensory aspects of blinking due to its perceived role in somesthesis (Kelly et al., 2012; Kurth et al., 2010).

The DLPFC was active during blinks, which may relate to the task focusing on blinking and deciding when to blink in relation to this. This region is more active during self-initiated blinks and therefore may relate to a conscious decision to blink (Van Eimeren et al., 2001). The DLPFC has not been identified in previous studies looking at the regions associated with blinking during a blink suppression paradigm (Berman et al., 2012; Lerner et al., 2009; Mazzone et al., 2010; Yoon et al., 2005), but most studies did not include event-related analysis of blinks and no studies have required participants to focus on their urges in order to give subjective ratings.

### 1.6.5 Validation of MESPFM

Using MESPFM, neuronal activation was identified within the right insula, cingulate areas, SMA and medial occipital cortex. These regions were found to be commonly active during suppression when data were analysed using the conventional GLM parametric approach.

The three clusters found with MESPFM showed similar activation of the right anterior insula and cingulate regions. The right insula was chosen as our region of interest for the estimation of the activation timeseries due to its consistent activation in fMRI studies of urge (Berman et al., 2012; Jackson et al., 2011; Lerner et al., 2009). Using a conventional analysis approach, we demonstrated that different portions of the right insula were active during suppression, urge and blinks. This is also shown in the activation timeseries obtained with MESPFM (shown in the supplementary material), where activations of the right insula were seen throughout the experiment regardless of task block. As the chosen activation maps relate to the activation seen during the corresponding timepoint we cannot separate suppression from feelings of urge if they happen simultaneously. Interestingly, we demonstrated that urge peaks at blink onset in the ‘Suppression’ blocks but not in the ‘Okay to blink’ blocks, suggesting that in healthy participants the urge-to-blink arises due to the act of suppression. Based on the results seen in our standard GLM analysis, separate subdivisions of the insula could be used in future as refined ROIs to estimate MESPFM activation timeseries to examine if it is possible to categorise cluster activation relating to suppression, urge and blinking separately (Kurth et al., 2010). As this work is a precursor for research looking at the urge-to-tic, it would be useful to see if the same subdivisions of the insula can be identified during a tic suppression paradigm when analysed using the conventional GLM approach.

Furthermore, the regions identified using the MESPFM approach are tighter than those identified using the conventional GLM approach, due to the low number of peaks identified because of the higher spatial and temporal specificity of MESPFM compared with the conventional GLM approach. Enhancing the sensitivity of the MESPFM algorithm to detect BOLD events, while preserving specificity, would lead to the identification of more peaks in the activation timeseries. This would give us more data across subjects and runs, and potentially facilitate the differentiation between urge and suppression networks, eliminating the requirement for continuous subjective urge ratings. Recent advancements in the MESPFM algorithm now incorporate the stability selection technique, eliminating the selection of the regularisation parameter utilized for estimating the activity-inducing signal (here, BIC was used, ensuring high specificity). These improvements demonstrate an increase in sensitivity, while maintaining the specificity of the activation events detected by the algorithm (Uruñuela et al., 2024).

## 1.7 Conclusion

In summary, this study suggests that the urge-to-act network is composed of regions involved in sensory processing and salience, while the action suppression network includes regions involved in executive control and response inhibition. The main findings are that separable regions within the insula contribute to different networks and there is a network overlap in the MCC and SMA that may act to determine when to perform a suppressed motor action. These are novel findings stemming from continuous measurement of urge, which allowed the two networks to be separated. However, the movement involved in this continuous urge rating affected the results due to activation of sensorimotor regions, meaning that we could not reliably ascertain whether these regions have a unique role in urge. Furthermore, the act of rating the urge itself could have affected how the participants experienced urge and therefore the BOLD response associated with it.

This study also validates the use of MESPFM as a timing-free approach to analyse fMRI data collected during action suppression paradigms where the event timings are unknown as might be the case during tic suppression in TS patients. Using the MESPFM approach, we were able to identify regions previously identified as being involved in the urge-to-act. The clusters identified with MESPFM showed an overlap with the regions involved in action suppression as shown by conventional analysis of the same data. Therefore, in future this approach could be used where the regions involved in urge and suppression could be identified without the need for subjective urge ratings.

## Data and Code Availability

The MESPFM algorithm is available in AFNI with the program 3dMEPFM https://afni.nimh.nih.gov/pub/dist/doc/program_help/3dMEPFM.html. The scripts used in the Paradigm Free Mapping analysis are available on GitHub https://github.com/MairiH/PFM_urgetoblink.

## Author Contributions

**MH.** Conceptualisation, Methodology, Software, Formal Analysis, Investigation, Writing – Original Draft

**EU.** Methodology, Software, Formal Analysis, Writing – Review & Editing

**CCG.** Conceptualisation, Methodology, Software, Formal Analysis, Writing – Review & Editing

**PG.** Conceptualisation, Methodology, Writing – Review & Editing, Funding Acquisition

**KD.** Investigation, Writing – Review & Editing

**VB.** Methodology, Software, Writing – Review & Editing

**IM.** Investigation

**RSP.** Methodology

**SJ.** Conceptualisation, Methodology, Writing – Review & Editing, Supervision, Funding Acquisition

## Acknowledgements

Data collection for this paper was supported by research grants from NIHR Nottingham Biomedical Research Centre, Tourettes Action (TA), Medical Research Council (MRC) (T032588) and Tourette Association of America (TAA). The views expressed are those of the authors and not necessarily those of the NHS, the NIHR, TA, MRC, TAA or the Department of Health.

## ABBREVIATIONS

ACC: Anterior cingulate cortex
AIC: Akaike information criterion
ATS: Activation time series
BIC: Bayesian information criterion
BOLD: Blood oxygenation level dependent
DLPFC: Dorsolateral prefrontal cortex
fMRI: functional magnetic resonance imaging
FOV: Field of view
GLM: General linear model
HRF: Haemodynamic response function
IFG: Inferior frontal gyrus
MCC: Mid-cingulate cortex
ME-ICA: Multi-echo independent component analysis
MESPFM: Multi-echo sparse paradigm free mapping
MFG: Medial frontal gyrus
OCD: Obsessive compulsive disorder
PCC: Posterior cingulate cortex
PFC: Prefrontal cortex
PU: Premonitory urge
R2*: transverse relaxation rate
ROI: Region of interest
SD: Standard deviation
SEM: Standard error of the mean
SFG: Superior frontal gyrus
SMA: Supplementary motor area
SMG: Supramarginal gyrus
SNR: Signal-to-noise ratio
TE: Echo time
TI: Inversion time
TR: Repetition time
TS: Tourette Syndrome
tSNR: total signal-to-noise ratio

## Appendix A: Temporal Signal-to-Noise Ratio (tSNR)

**Figure A.1.**
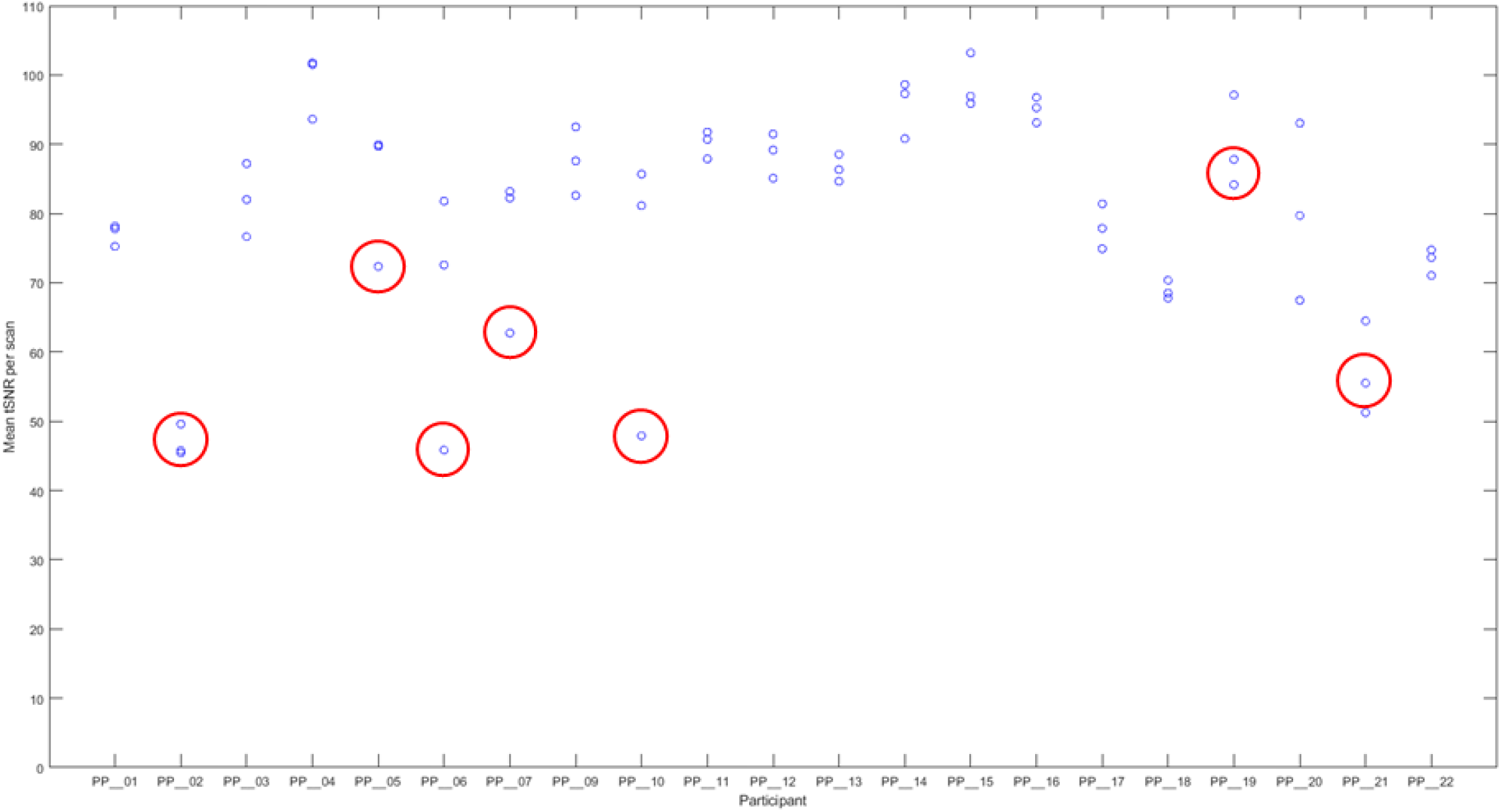
A graph showing the mean tSNR for each fMRI run of the blink suppression paradigm, where scans encircled in red were excluded due to a maintained absolute mean displacement over 1.5mm. If found, scans with a tSNR below 30 would have een excluded.

**Figure A.2.**
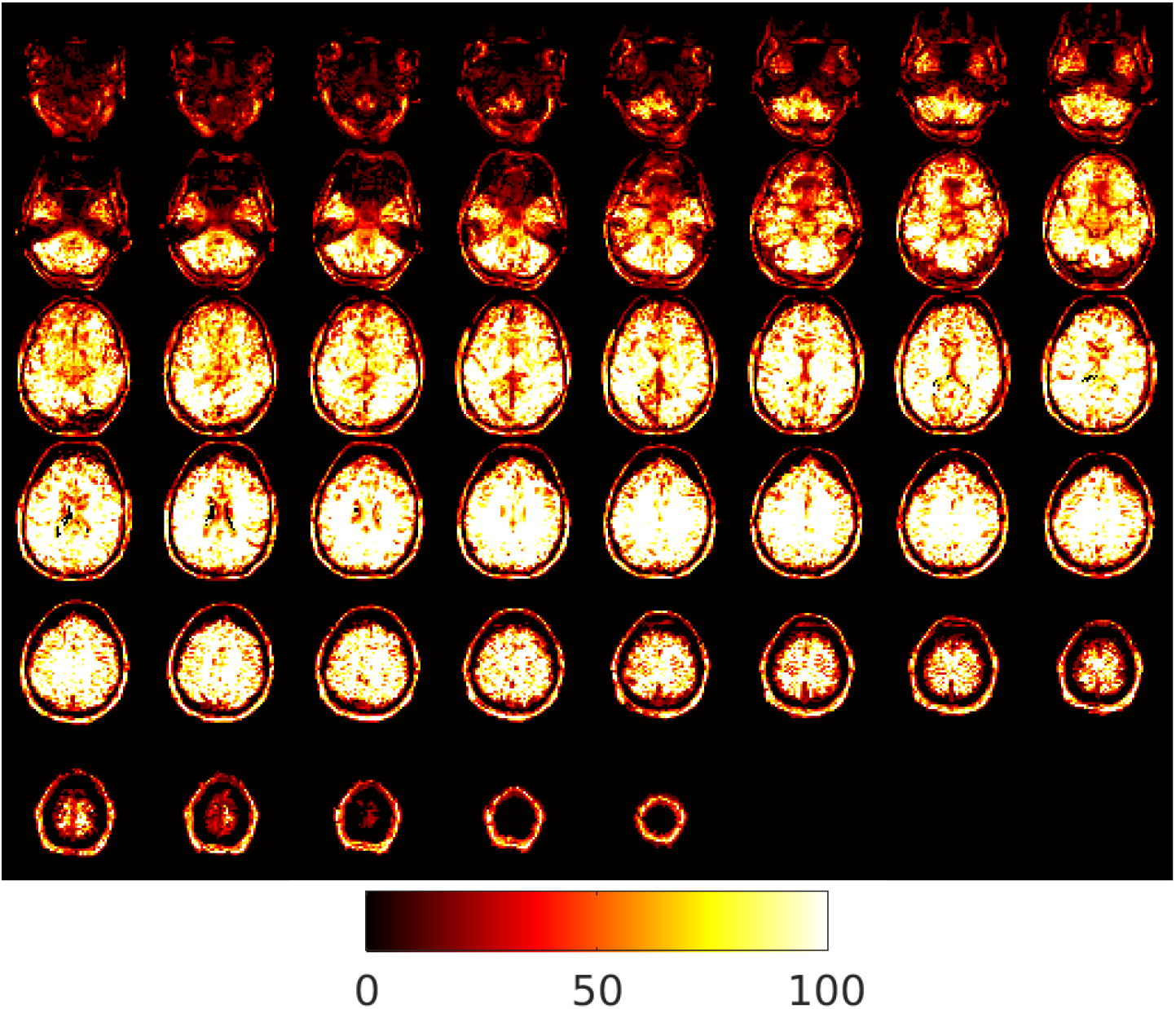
An example (Sub01 run01) fMRI image with high tSNR.

**Figure A.3.**
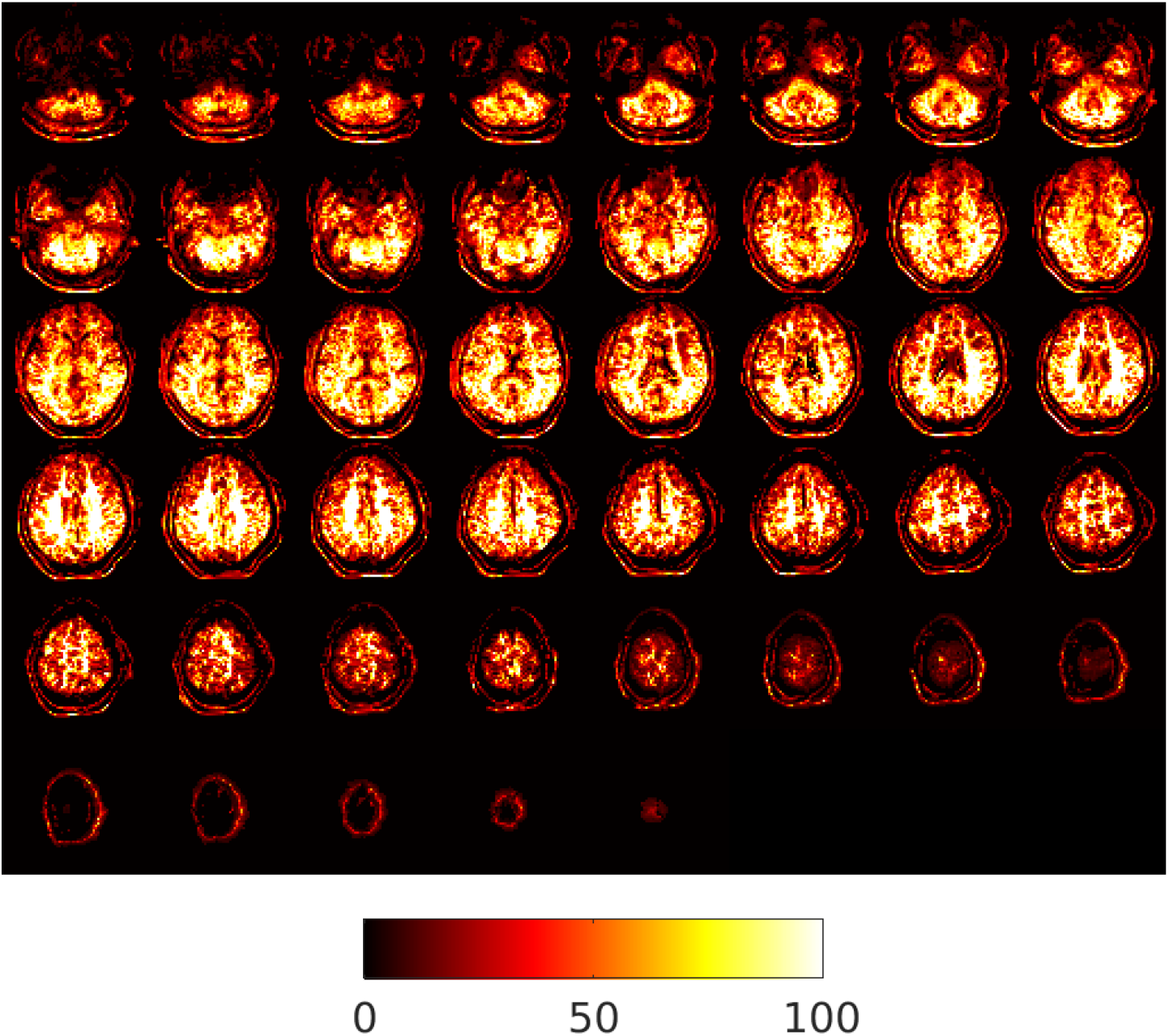
An example (Sub02 run01) fMRI scan which was excluded due to a maintained absolute mean displacement over 1.5mm which caused a drop in tSNR.

## Appendix B: Local Maxima Cluster Index

**Table B.1.**
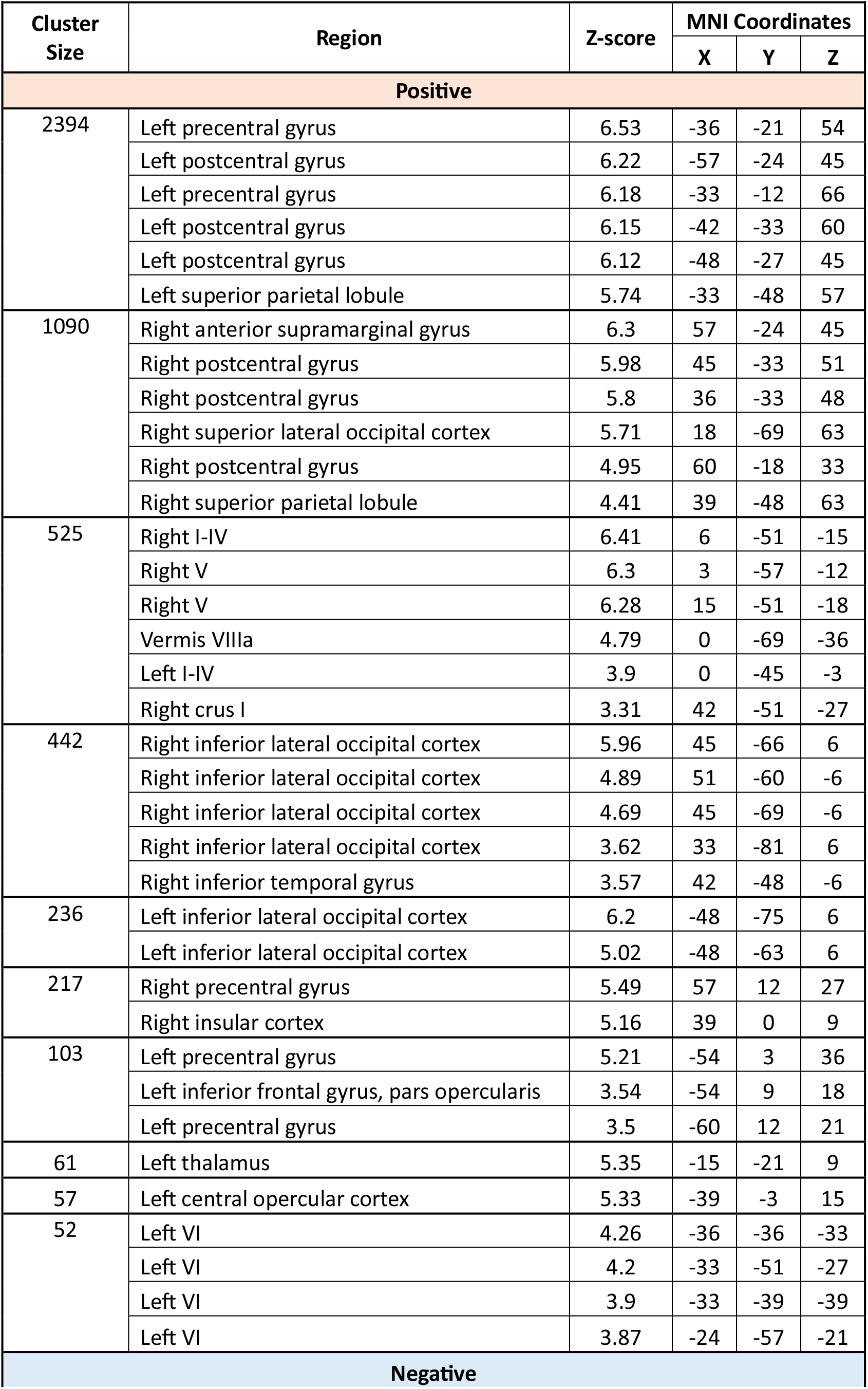
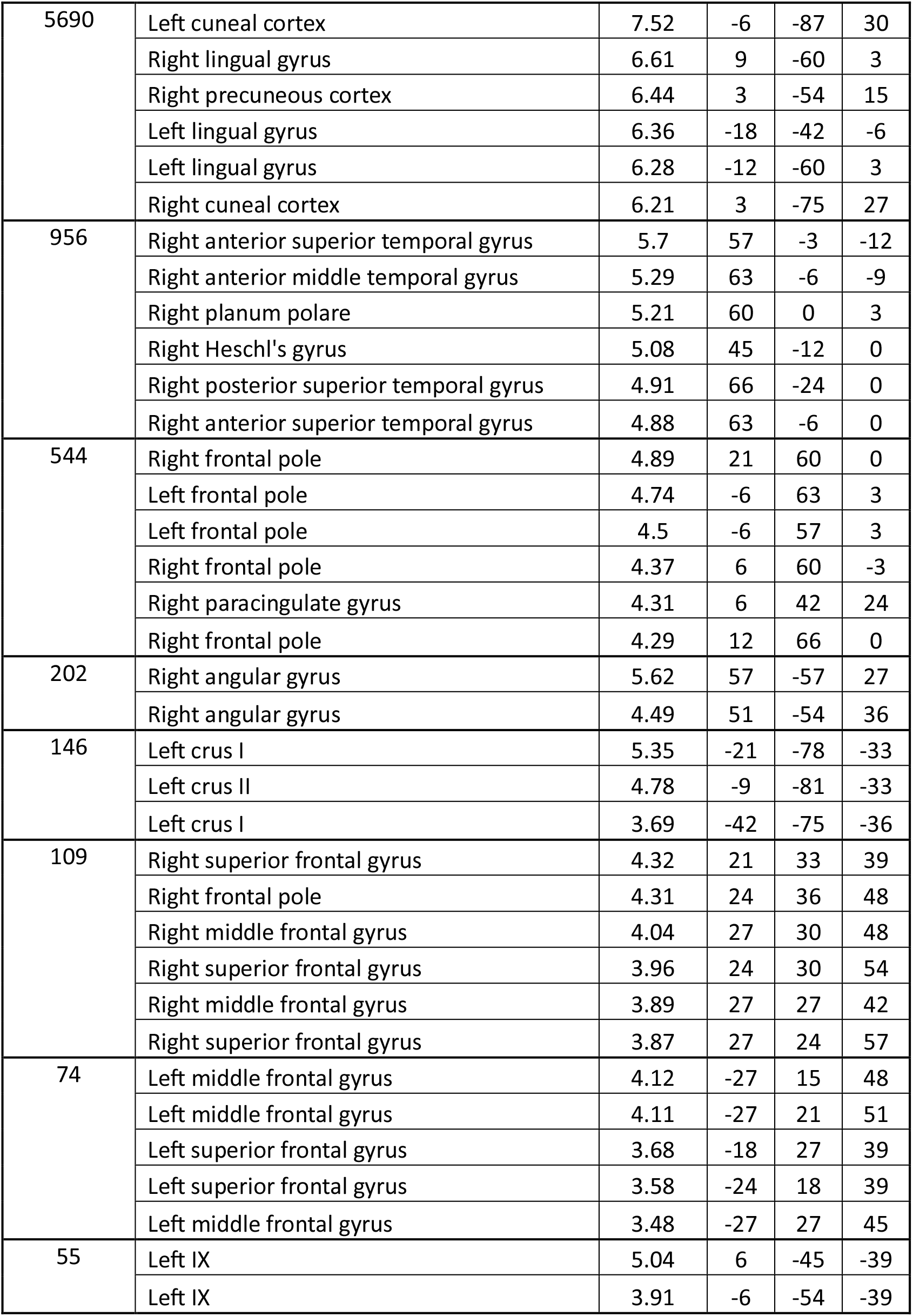
Local maxima cluster index for ‘Suppress’ blocks.

**Table B.2.** . Local maxima cluster index for ‘Okay to blink’ blocks.

| Cluster Size | Region | Z-score | MNI Coordinates |  |  |
| --- | --- | --- | --- | --- | --- |
|  |  |  | X | Y | Z |
| Positive |  |  |  |  |  |
| 1259 | Left postcentral gyrus | 5.73 | -57 | -27 | 48 |
|  | Left postcentral gyrus | 5.46 | -39 | -36 | 54 |
|  | Left postcentral gyrus | 5.39 | -45 | -27 | 45 |
|  | Left precentral gyrus | 5.3 | -42 | -21 | 60 |
|  | Left postcentral gyrus | 5.22 | -36 | -36 | 45 |
|  | Left superior parietal lobule | 5.14 | -33 | -45 | 54 |
| 797 | Right precentral gyrus | 5.92 | 54 | -21 | 42 |
|  | Right postcentral gyrus | 5.56 | 42 | -30 | 48 |
|  | Right superior lateral occipital cortex | 4.8 | 18 | -63 | 63 |
|  | Right superior parietal lobule | 4.71 | 39 | -48 | 63 |
| 460 | Right inferior lateral occipital cortex | 5.48 | 45 | -69 | 6 |
|  | Right inferior lateral occipital cortex | 4.94 | 54 | -60 | -6 |
|  | Right inferior lateral occipital cortex | 4.45 | 45 | -69 | -6 |
|  | Right superior lateral occipital cortex | 4.12 | 30 | -84 | 9 |
|  | Right inferior temporal gyrus | 4 | 45 | -48 | -6 |
|  | Right cerebral white matter (superior longitudinal fasciculus) | 3.85 | 42 | -51 | 3 |
| 212 | Right precentral gyrus | 5.07 | 33 | -6 | 57 |
|  | Right precentral gyrus | 3.63 | 15 | -12 | 72 |
|  | Right supplementary motor area | 3.24 | 12 | -9 | 54 |
| 165 | Left inferior lateral occipital cortex | 6.23 | -45 | -72 | 6 |
|  | Left cerebral white matter (inferior longitudinal fasciculus) | 3.84 | -36 | -63 | 9 |
| 135 | Right V | 4.7 | 12 | -51 | -18 |
|  | Right V | 4.46 | 24 | -42 | -30 |
| 96 | Right precentral gyrus | 5.09 | 57 | 12 | 27 |
| Negative |  |  |  |  |  |
| 4237 | Left cuneal cortex | 6.57 | -6 | -87 | 27 |
|  | Left lingual gyrus | 6.42 | -12 | -60 | 3 |
|  | Left cuneal cortex | 6.42 | -9 | -87 | 18 |
|  | Left intracalcarine cortex | 6.22 | -12 | -75 | 15 |
|  | Right lingual gyrus | 6.21 | 12 | -60 | 3 |
|  | Right precuneous cortex | 6.14 | 18 | -57 | 9 |
| 903 | Right central opercular cortex | 5.46 | 60 | -3 | 6 |
|  | Right anterior superior temporal gyrus | 5 | 57 | -3 | -9 |
|  | Right temporal pole | 4.65 | 54 | 6 | -15 |
|  | Right planum polare | 4.57 | 42 | 0 | -15 |
|  | Right middle temporal gyrus | 4.51 | 66 | -39 | 3 |
|  | Right planum polare | 4.49 | 63 | -18 | 3 |
| 311 | Right angular gyrus | 5.49 | 57 | -54 | 24 |
|  | Right angular gyrus | 4.91 | 57 | -54 | 36 |
|  | Right posterior supramarginal gyrus | 3.82 | 48 | -39 | 21 |
|  | Right angular gyrus | 3.74 | 39 | -51 | 21 |
|  | Right angular gyrus | 3.59 | 54 | -54 | 51 |
| 283 | Right frontal pole | 5.12 | 27 | 54 | 36 |
|  | Right frontal pole | 4.54 | 18 | 72 | 6 |
|  | Right frontal pole | 4.25 | 21 | 63 | 0 |
|  | Right frontal pole | 4.2 | 30 | 48 | 30 |
|  | Right frontal pole | 4.04 | 30 | 57 | 18 |
|  | Right frontal pole | 4.02 | 27 | 45 | 39 |
| 175 | Left insular cortex | 4.54 | -42 | -6 | -9 |
|  | Left planum temporale | 4.34 | -60 | -9 | 3 |
|  | Left central opercular cortex | 4.04 | -51 | -6 | 12 |
|  | Left insular cortex | 4.01 | -33 | 12 | -18 |
|  | Left planum polare | 4.01 | -54 | 0 | -6 |
|  | Left temporal pole | 3.49 | -30 | 6 | -21 |
| 172 | Anterior cingulate cortex | 4.44 | 6 | 12 | 39 |
|  | Paracingulate gyrus | 3.99 | 6 | 33 | 30 |
|  | Anterior cingulate cortex | 3.96 | -6 | 39 | 18 |
|  | Anterior cingulate cortex | 3.95 | 0 | 33 | 24 |
|  | Anterior cingulate cortex | 3.85 | 3 | 39 | 21 |
|  | Anterior cingulate cortex | 3.42 | 6 | 45 | 6 |
| 162 | Left crus I | 5.65 | -18 | -78 | -33 |
|  | Left crus I | 3.9 | -15 | -90 | -24 |
|  | Left VI | 3.78 | -9 | -72 | -24 |
|  | Vermis VI | 3.58 | 3 | -72 | -21 |
| 102 | Left posterior supramarginal gyrus | 4.9 | -54 | -48 | 18 |
|  | Left middle temporal gyrus | 3.97 | -57 | -48 | 6 |
|  | Left middle temporal gyrus | 3.7 | -66 | -54 | 6 |
|  | Left posterior supramarginal gyrus | 3.52 | -57 | -45 | 33 |
|  | Left planum temporale | 3.36 | -57 | -33 | 15 |
| 30 | Right middle frontal gyrus | 3.9 | 39 | 27 | 33 |

**Table B.3.** Local maxima cluster index for ‘Random’ active baseline blocks.

| Cluster Size | Region | Z-score | MNI Coordinates |  |  |
| --- | --- | --- | --- | --- | --- |
|  |  |  | X | Y | Z |
| Positive |  |  |  |  |  |
| 1986 | Left postcentral gyrus | 6.26 | -57 | -24 | 45 |
|  | Left precentral gyrus | 6.18 | -36 | -21 | 54 |
|  | Left postcentral gyrus | 6.13 | -42 | -33 | 60 |
|  | Left precentral gyrus | 6.03 | -33 | -12 | 63 |
|  | Left postcentral gyrus | 6.01 | -48 | -27 | 45 |
|  | Left superior parietal lobule | 5.73 | -33 | -48 | 57 |
| 939 | Right anterior supramarginal gyrus | 5.96 | 57 | -24 | 45 |
|  | Right postcentral gyrus | 5.75 | 42 | -33 | 48 |
|  | Right postcentral gyrus | 5.68 | 33 | -36 | 48 |
|  | Right superior lateral occipital cortex | 5.16 | 18 | -66 | 63 |
|  | Right postcentral gyrus | 5.12 | 60 | -15 | 33 |
|  | Right superior parietal lobule | 4.38 | 39 | -48 | 60 |
| 500 | Right inferior lateral occipital cortex | 5.89 | 45 | -69 | 6 |
|  | Right inferior lateral occipital cortex | 4.91 | 51 | -60 | -6 |
|  | Right inferior lateral occipital cortex | 4.59 | 45 | -69 | -6 |
|  | Right cerebral white matter (inferior longitudinal fasciculus) | 4.4 | 39 | -60 | -3 |
|  | Right superior lateral occipital cortex | 4.36 | 30 | -84 | 9 |
|  | Right inferior temporal cortex | 3.85 | 42 | -48 | -6 |
| 364 | Right V | 6.09 | 15 | -51 | -18 |
|  | Right I-IV | 5.87 | 6 | -51 | -15 |
|  | Right V | 5.59 | 3 | -57 | -12 |
|  | Left I-IV | 3.4 | 0 | -45 | -3 |
| 205 | Left inferior lateral occipital cortex | 6.45 | -48 | -75 | 6 |
|  | Left inferior lateral occipital cortex | 4.63 | -48 | -63 | 6 |
| 123 | Right precentral gyrus | 5.49 | 57 | 12 | 27 |
| 102 | Left precentral gyrus | 5.23 | -54 | 3 | 33 |
| 66 | Vermis VIIIa | 4.7 | 3 | -66 | -36 |
| 50 | Left thalamus | 4.98 | -15 | -24 | 9 |
| 39 | Left inferior lateral occipital cortex | 4.14 | -27 | -90 | 6 |
| 36 | Left central opercular cortex | 5.19 | -39 | -3 | 15 |
| Negative |  |  |  |  |  |
| 4473 | Left cuneal cortex | 7.33 | -6 | -87 | 30 |
|  | Left lingual gyrus | 6.87 | -12 | -60 | 3 |
|  | Left intracalcarine cortex | 6.73 | -6 | -75 | 15 |
|  | Right cuneal cortex | 6.71 | 3 | -75 | 27 |
|  | Left lingual gyrus | 6.63 | -18 | -42 | -6 |
|  | Right lingual gyrus | 6.58 | 12 | -60 | 3 |
| 1007 | Right central opercular cortex | 5.95 | 60 | 0 | 6 |
|  | Right anterior superior temporal gyrus | 5.54 | 57 | 0 | -15 |
|  | Right temporal pole | 4.99 | 45 | 9 | -27 |
|  | Right Heschl's gyrus | 4.95 | 45 | -12 | 0 |
|  | Right planum polare | 4.86 | 45 | -3 | -12 |
|  | Right temporal pole | 4.85 | 45 | 15 | -15 |
| 571 | Right paracingulate gyrus | 4.54 | 6 | 42 | 21 |
|  | Right frontal pole | 4.44 | 21 | 60 | 0 |
|  | Right paracingulate gyrus | 4.29 | 12 | 42 | 18 |
|  | Right frontal pole | 4.27 | 24 | 54 | 12 |
|  | Right frontal pole | 4.19 | 24 | 57 | 27 |
|  | Right frontal pole | 4.13 | 24 | 54 | -3 |
| 313 | Right angular gyrus | 5.72 | 57 | -54 | 24 |
|  | Right angular gyrus | 4.87 | 51 | -54 | 36 |
|  | Right angular gyrus | 3.88 | 54 | -54 | 51 |
|  | Right angular gyrus | 3.78 | 39 | -51 | 21 |
|  | Right posterior supramarginal gyrus | 3.49 | 48 | -39 | 21 |
| 279 | Left planum polare | 4.69 | -51 | -3 | -6 |
|  | Left insular cortex | 4.59 | -42 | -3 | -12 |
|  | Left planum polare | 4.38 | -57 | -9 | 3 |
|  | Left frontal orbital cortex | 4.25 | -33 | 12 | -21 |
|  | Left anterior superior temporal gyrus | 4.24 | -63 | -12 | 0 |
|  | Left anterior superior temporal gyrus | 4.1 | -54 | -12 | -6 |
| 240 | Left angular gyrus | 5.2 | -54 | -51 | 18 |
|  | Left posterior superior temporal gyrus | 4.13 | -48 | -33 | 3 |
|  | Left planum polare | 3.98 | -60 | -33 | 15 |
|  | Left posterior supramarginal gyrus | 3.97 | -63 | -48 | 21 |
|  | Left superior lateral occipital cortex | 3.85 | -57 | -63 | 21 |
|  | Left posterior middle temporal gyrus | 3.7 | -45 | -42 | 3 |
| 229 | Right middle frontal gyrus | 4.59 | 39 | 9 | 45 |
|  | Right superior frontal gyrus | 4.43 | 21 | 33 | 39 |
|  | Right middle frontal gyrus | 4.17 | 27 | 33 | 48 |
|  | Right superior frontal gyrus | 3.96 | 24 | 24 | 51 |
|  | Right middle frontal gyrus | 3.9 | 36 | 12 | 33 |
|  | Right superior frontal gyrus | 3.88 | 27 | 24 | 57 |
| 152 | Left crus I | 5.38 | -21 | -81 | -30 |
|  | Left crus II | 4.96 | -9 | -78 | -33 |
| 64 | Left insular cortex | 4.06 | -36 | 18 | -3 |
|  | Left insular cortex | 3.97 | -27 | 27 | 0 |
|  | Left frontal operculum cortex | 3.64 | -30 | 27 | 9 |
|  | Left putamen | 3.55 | -21 | 18 | -3 |
|  | Left cerebral white matter (inferior fronto-occipital fasciculus) | 3.49 | -24 | 27 | 12 |
| 50 | Left crus I | 4.13 | -42 | -66 | -42 |
|  | Left crus II | 3.87 | -36 | -72 | -42 |
|  | Left crus I | 3.8 | -42 | -75 | -36 |
|  | Left crus I | 3.73 | -54 | -63 | -30 |
| 49 | Right IX | 4.59 | 3 | -45 | -42 |
|  | Left IX | 3.84 | -9 | -51 | -33 |
| 31 | Right thalamus | 4.07 | 6 | -9 | 12 |
|  | Right thalamus | 3.52 | 3 | -21 | 3 |
|  | Left thalamus | 3.41 | -3 | -6 | 6 |

**Table B.4.**
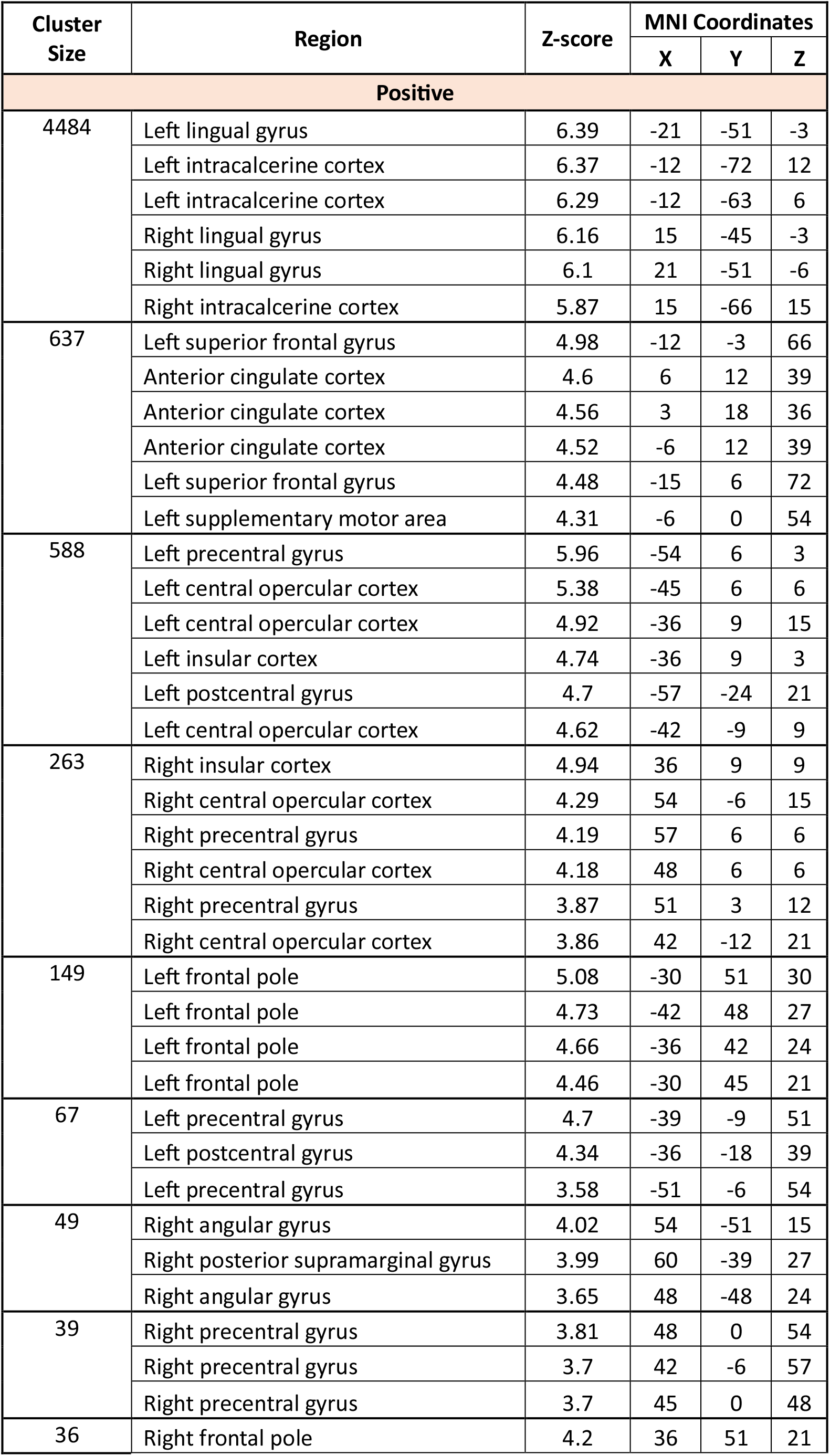
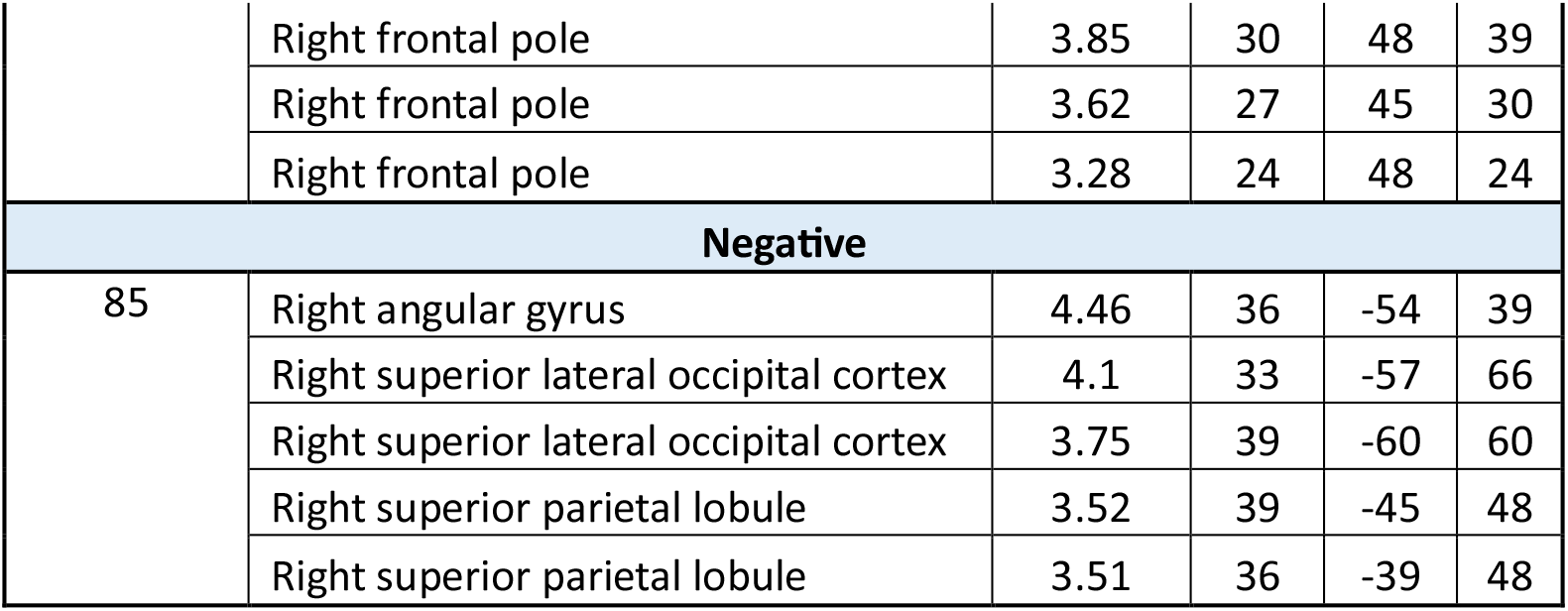
Local maxima cluster index for blinks.

| Cluster Size | Region | Z-score | MNI Coordinates |  |  |
| --- | --- | --- | --- | --- | --- |
|  |  |  | X | Y | Z |
| Positive |  |  |  |  |  |
| 4484 | Left lingual gyrus | 6.39 | -21 | -51 | -3 |
|  | Left intracalcerine cortex | 6.37 | -12 | -72 | 12 |
|  | Left intracalcerine cortex | 6.29 | -12 | -63 | 6 |
|  | Right lingual gyrus | 6.16 | 15 | -45 | -3 |
|  | Right lingual gyrus | 6.1 | 21 | -51 | -6 |
|  | Right intracalcerine cortex | 5.87 | 15 | -66 | 15 |
| 637 | Left superior frontal gyrus | 4.98 | -12 | -3 | 66 |
|  | Anterior cingulate cortex | 4.6 | 6 | 12 | 39 |
|  | Anterior cingulate cortex | 4.56 | 3 | 18 | 36 |
|  | Anterior cingulate cortex | 4.52 | -6 | 12 | 39 |
|  | Left superior frontal gyrus | 4.48 | -15 | 6 | 72 |
|  | Left supplementary motor area | 4.31 | -6 | 0 | 54 |
| 588 | Left precentral gyrus | 5.96 | -54 | 6 | 3 |
|  | Left central opercular cortex | 5.38 | -45 | 6 | 6 |
|  | Left central opercular cortex | 4.92 | -36 | 9 | 15 |
|  | Left insular cortex | 4.74 | -36 | 9 | 3 |
|  | Left postcentral gyrus | 4.7 | -57 | -24 | 21 |
|  | Left central opercular cortex | 4.62 | -42 | -9 | 9 |
| 263 | Right insular cortex | 4.94 | 36 | 9 | 9 |
|  | Right central opercular cortex | 4.29 | 54 | -6 | 15 |
|  | Right precentral gyrus | 4.19 | 57 | 6 | 6 |
|  | Right central opercular cortex | 4.18 | 48 | 6 | 6 |
|  | Right precentral gyrus | 3.87 | 51 | 3 | 12 |
|  | Right central opercular cortex | 3.86 | 42 | -12 | 21 |
| 149 | Left frontal pole | 5.08 | -30 | 51 | 30 |
|  | Left frontal pole | 4.73 | -42 | 48 | 27 |
|  | Left frontal pole | 4.66 | -36 | 42 | 24 |
|  | Left frontal pole | 4.46 | -30 | 45 | 21 |
| 67 | Left precentral gyrus | 4.7 | -39 | -9 | 51 |
|  | Left postcentral gyrus | 4.34 | -36 | -18 | 39 |
|  | Left precentral gyrus | 3.58 | -51 | -6 | 54 |
| 49 | Right angular gyrus | 4.02 | 54 | -51 | 15 |
|  | Right posterior supramarginal gyrus | 3.99 | 60 | -39 | 27 |
|  | Right angular gyrus | 3.65 | 48 | -48 | 24 |
| 39 | Right precentral gyrus | 3.81 | 48 | 0 | 54 |
|  | Right precentral gyrus | 3.7 | 42 | -6 | 57 |
|  | Right precentral gyrus | 3.7 | 45 | 0 | 48 |
| 36 | Right frontal pole | 4.2 | 36 | 51 | 21 |
|  | Right frontal pole | 3.85 | 30 | 48 | 39 |
|  | Right frontal pole | 3.62 | 27 | 45 | 30 |
|  | Right frontal pole | 3.28 | 24 | 48 | 24 |
| **Negative** |  |  |  |  |  |
| 85 | Right angular gyrus | 4.46 | 36 | -54 | 39 |
|  | Right superior lateral occipital cortex | 4.1 | 33 | -57 | 66 |
|  | Right superior lateral occipital cortex | 3.75 | 39 | -60 | 60 |
|  | Right superior parietal lobule | 3.52 | 39 | -45 | 48 |
|  | Right superior parietal lobule | 3.51 | 36 | -39 | 48 |

**Table B.5.** Local maxima cluster index related to the subjective urge ratings.

| Cluster Size | Region | Z-score | MNI Coordinates |  |  |
| --- | --- | --- | --- | --- | --- |
|  |  |  | X | Y | Z |
| Positive |  |  |  |  |  |
| 1317 | Right anterior thalamic radiation | 4.91 | 3 | -30 | 3 |
|  | Left thalamus | 4.61 | -6 | -33 | 3 |
|  | Left intracalcarine cortex | 4.49 | -12 | -72 | 15 |
|  | Left lingual gyrus | 4.42 | -12 | -78 | -3 |
|  | Right occipital pole | 4.35 | 9 | -99 | 9 |
|  | Right thalamus | 4.32 | 18 | -27 | 0 |
| 110 | Left insular cortex | 4.58 | -42 | -9 | 0 |
|  | Left insular cortex | 4.52 | -39 | -15 | 6 |
| 69 | Right planum polare | 4.02 | 48 | -9 | -3 |
|  | Right insular cortex | 3.99 | 39 | -9 | 0 |
|  | Right planum polare | 3.78 | 48 | 3 | -3 |
|  | Right Heschl's gyrus | 3.73 | 42 | -18 | 3 |
|  | Right planum polare | 3.49 | 45 | -3 | -9 |
| 50 | Left insular cortex | 3.9 | -30 | 12 | -12 |
|  | Left putamen | 3.88 | -21 | 6 | -9 |
|  | Left putamen | 3.36 | -15 | 12 | -9 |
| 41 | Paracingulate gyrus | 3.98 | -3 | 33 | 33 |
|  | Anterior cingulate gyrus | 3.87 | 0 | 24 | 33 |
| 37 | Right central opercular cortex | 3.79 | 48 | -6 | 9 |
|  | Right insular cortex | 3.66 | 33 | -12 | 18 |
| Negative |  |  |  |  |  |
| 49 | Right superior lateral occipital cortex | 4.29 | 18 | -75 | 57 |
|  | Right superior lateral occipital cortex | 3.89 | 27 | -72 | 57 |
|  | Right precuneous cortex | 3.47 | 9 | -69 | 54 |
| 42 | Right posterior supramarginal gyrus | 4.18 | 39 | -45 | 39 |
|  | Right posterior supramarginal gyrus | 3.99 | 36 | -39 | 36 |
| 33 | Left superior lateral occipital cortex | 3.96 | -21 | -63 | 51 |
|  | Left precuneous cortex | 3.68 | -6 | -69 | 51 |

**Table B.6.**
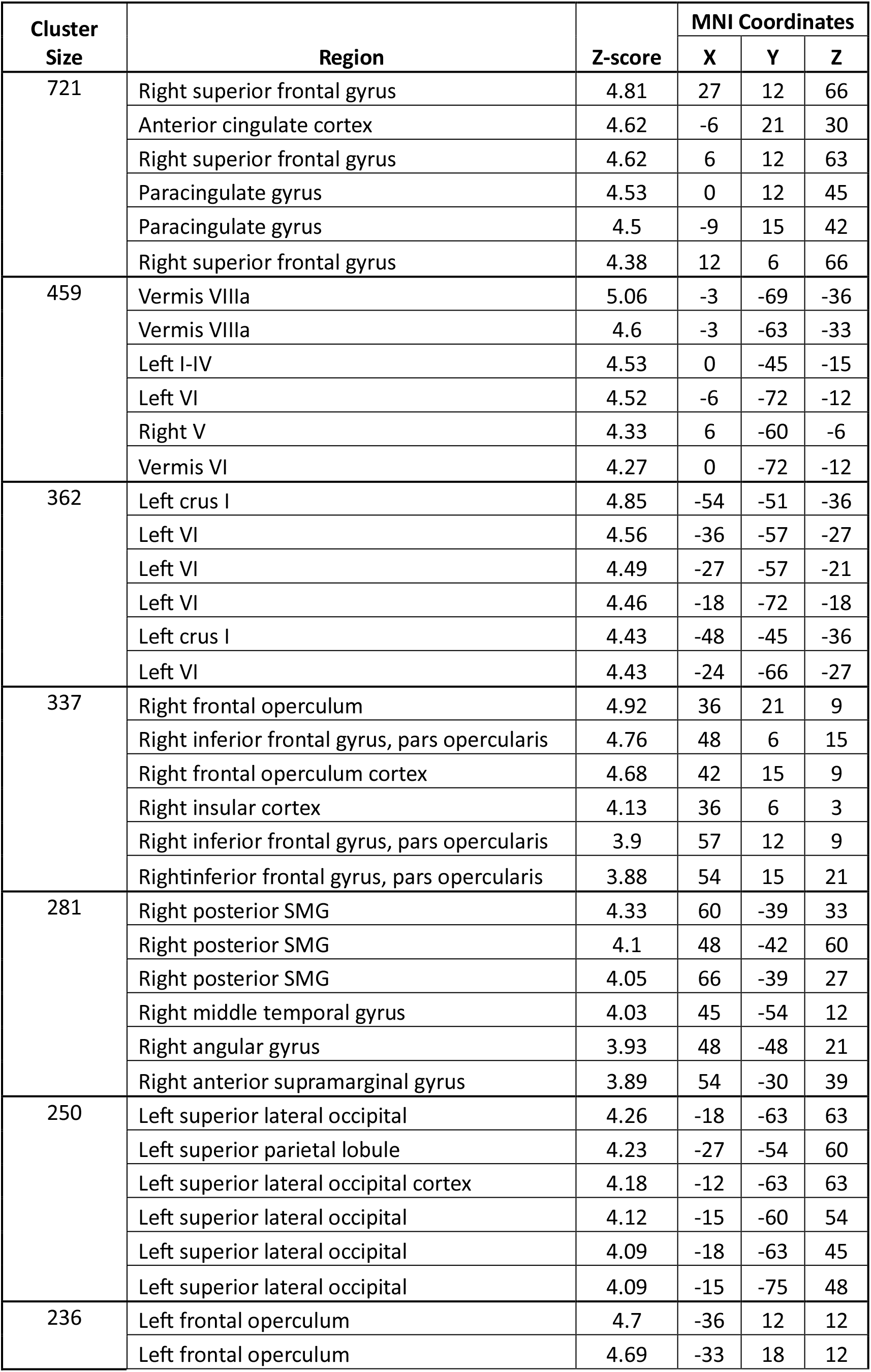
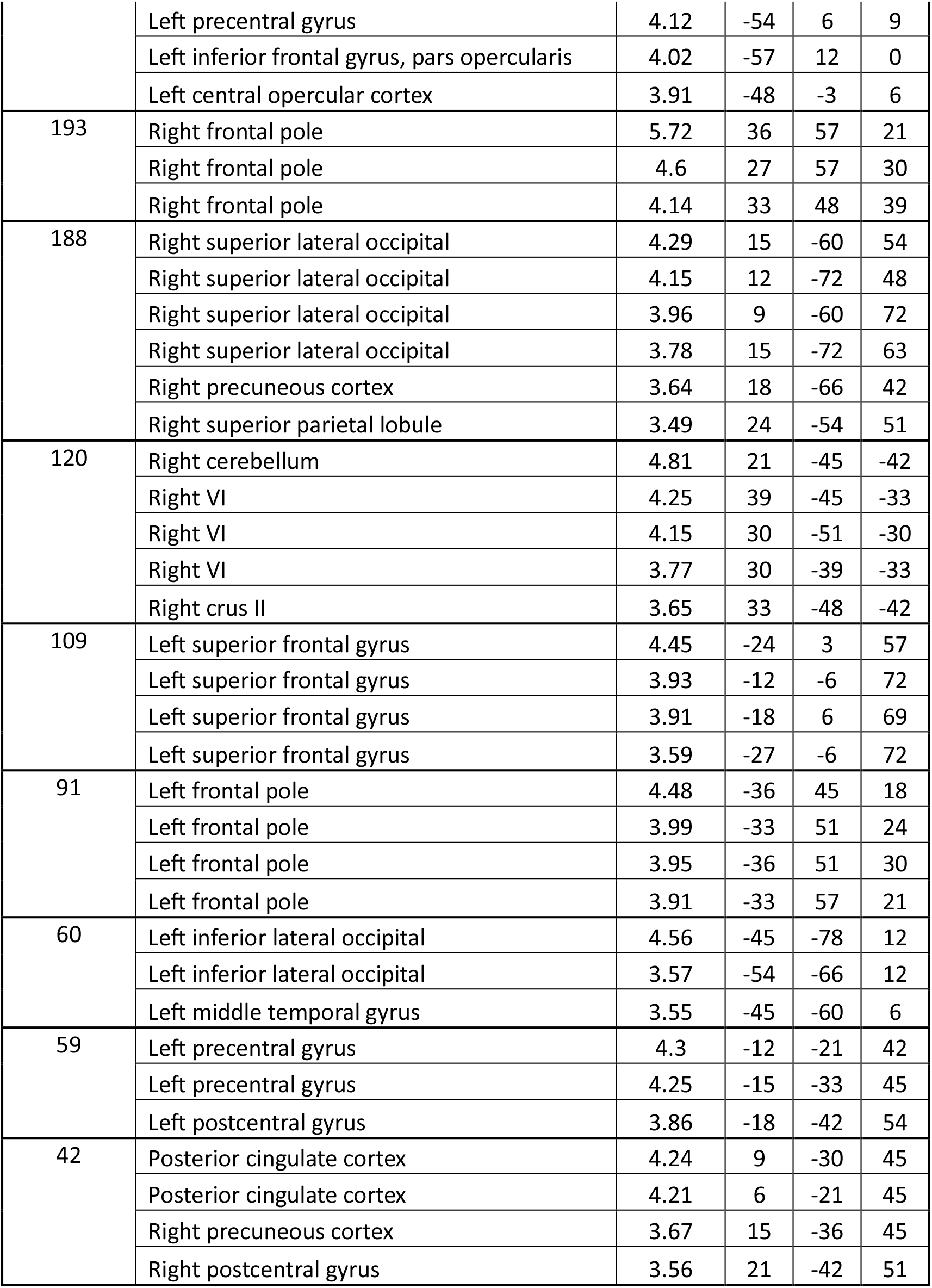
Local maxima cluster index when contrasting ‘Suppress’ > ’Okay to blink’ blocks.

**Table B.7.** Local maxima cluster index when contrasting ’Okay to blink’ > ‘Suppress’ blocks.

| Cluster Size | Region | Z-score | MNI Coordinates |  |  |
| --- | --- | --- | --- | --- | --- |
|  |  |  | X | Y | Z |
| 261 | Posterior cingulate cortex | 5 | 3 | -48 | 21 |
|  | Posterior cingulate cortex | 4.25 | -9 | -51 | 21 |
|  | Posterior cingulate cortex | 4.04 | 0 | -42 | 36 |
|  | Posterior cingulate cortex | 4.02 | -6 | -45 | 12 |
|  | Left precuneous cortex | 3.84 | -3 | -60 | 36 |
| 207 | Left middle frontal gyrus | 4.6 | -27 | 18 | 48 |
|  | Left middle frontal gyrus | 4.58 | -33 | 18 | 54 |
|  | Left superior frontal gyrus | 4.43 | -15 | 24 | 48 |
|  | Left superior frontal gyrus | 4.39 | -21 | 27 | 48 |
|  | Left frontal lobe | 3.86 | -15 | 45 | 54 |
|  | Paracingulate gyrus | 3.84 | -3 | 39 | 36 |
| 203 | Left superior lateral occipital | 4.63 | -42 | -72 | 42 |
|  | Left superior lateral occipital | 4.52 | -39 | -63 | 39 |
|  | Left superior lateral occipital | 4.39 | -45 | -72 | 33 |
|  | Left superior lateral occipital | 4.33 | -36 | -72 | 48 |
|  | Left angular gyrus | 4.12 | -42 | -60 | 33 |
| 52 | Frontal pole | 4.55 | 0 | 63 | -6 |
|  | Right frontal pole | 4.48 | 6 | 63 | -6 |
| 52 | Left hippocampus | 3.77 | -36 | -18 | -12 |
|  | Left parahippocampal gyrus | 3.58 | -27 | -33 | -12 |
|  | Left amygdala | 3.55 | -27 | -9 | -18 |
| 51 | Left frontal orbital cortex | 4.61 | -48 | 36 | -9 |
|  | Left frontal pole | 4.03 | -42 | 45 | -6 |
| 44 | Right frontal pole | 4.04 | 18 | 36 | 48 |
|  | Right superior frontal gyrus | 3.79 | 18 | 36 | 57 |
| 42 | Right crus I | 4.04 | 27 | -81 | -24 |
|  | Right crus II | 3.93 | 24 | -84 | -36 |
|  | Right crus I | 3.84 | 30 | -84 | -30 |
|  | Right crus I | 3.67 | 39 | -81 | -27 |
|  | Right crus II | 3.54 | 15 | -84 | -36 |

**Table B.8.** Local maxima cluster index when contrasting ‘Random’ > Urge blocks.

| Cluster Size | Region | Z-score | MNI Coordinates |  |  |
| --- | --- | --- | --- | --- | --- |
|  |  |  | X | Y | Z |
| 1660 | Left precentral gyrus | 6.53 | -36 | -21 | 54 |
|  | Left postcentral gyrus | 6.37 | -57 | -24 | 45 |
|  | Left postcentral gyrus | 6.19 | -42 | -33 | 60 |
|  | Left postcentral gyrus | 6.19 | -48 | -27 | 45 |
|  | Left precentral gyrus | 6.07 | -33 | -12 | 63 |
|  | Left superior parietal lobule | 5.76 | -33 | -48 | 57 |
| 953 | Right anterior SMG | 6.26 | 57 | -24 | 45 |
|  | Right postcentral gyrus | 5.78 | 45 | -33 | 51 |
|  | Right postcentral gyrus | 5.72 | 36 | -33 | 48 |
|  | Right superior lateral occipital | 5.47 | 18 | -69 | 63 |
|  | Right postcentral gyrus | 4.84 | 60 | -15 | 33 |
|  | Right superior parietal lobule | 4.32 | 39 | -48 | 60 |
| 437 | Right inferior lateral occipital | 6.02 | 45 | -69 | 6 |
|  | Right inferior temporal gyrus | 4.96 | 51 | -57 | -6 |
|  | Right inferior temporal gyrus | 4.13 | 42 | -48 | -6 |
|  | Right superior lateral occipital | 3.78 | 30 | -84 | 9 |
| 386 | Right V | 6.48 | 15 | -51 | -18 |
|  | Right I-IV | 6.3 | 6 | -51 | -15 |
|  | Right VI | 3.83 | 15 | -63 | -21 |
|  | Left I-IV | 3.59 | 0 | -45 | -3 |
| 386 | Right precentral gyrus | 5.05 | 27 | -9 | 51 |
|  | Right precentral gyrus | 4.82 | 27 | -9 | 57 |
|  | Right precentral gyrus | 4.75 | 27 | -12 | 63 |
|  | Right precentral gyrus | 4.67 | 33 | -12 | 60 |
|  | Right superior frontal gyrus | 4.43 | 24 | 3 | 60 |
|  | Right precentral gyrus | 3.6 | 15 | -9 | 63 |
| 205 | Left inferior lateral occipital | 6.47 | -45 | -72 | 6 |
|  | Left inferior lateral occipital | 4.87 | -48 | -63 | 6 |
| 120 | Right precentral gyrus | 5.56 | 57 | 12 | 27 |
| 85 | Left precentral gyrus | 5.19 | -54 | 3 | 36 |
| 72 | Vermis VIIIa | 4.72 | 3 | -66 | -36 |
| 47 | Left thalamus | 4.93 | -18 | -21 | 9 |
| 31 | Left central opercular cortex | 5.07 | -39 | -3 | 15 |

**Table B.9.** Local maxima cluster index when contrasting Urge > ‘Random’ blocks.

| Cluster Size | Region | Z-score | MNI Coordinates |  |  |
| --- | --- | --- | --- | --- | --- |
|  |  |  | X | Y | Z |
| 5595 | Left cuneal cortex | 8.05 | -6 | -87 | 30 |
|  | Left lingual gyrus | 6.87 | -18 | -42 | -6 |
|  | Left lingual gyrus | 6.8 | -12 | -60 | 3 |
|  | Left intracalcarine cortex | 6.79 | -6 | -75 | 15 |
|  | Right cuneal cortex | 6.73 | 3 | -75 | 27 |
|  | Right lingual gyrus | 6.68 | 9 | -60 | 3 |
| 1176 | Right central opercular cortex | 5.99 | 60 | 0 | 6 |
|  | Right anterior superior temporal gyrus | 5.69 | 57 | 0 | -15 |
|  | Right Heschl's gyrus | 5.21 | 45 | -12 | 0 |
|  | Right temporal pole | 5.05 | 45 | 9 | -27 |
|  | Right temporal pole | 4.93 | 45 | 15 | -15 |
|  | Right temporal pole | 4.91 | 36 | 6 | -18 |
| 665 | Paracingulate cortex | 4.76 | 6 | 42 | 21 |
|  | Paracingulate cortex | 4.57 | 0 | 42 | 27 |
|  | Right frontal pole | 4.49 | 21 | 60 | 0 |
|  | Anterior cingulate cortex | 4.42 | -3 | 33 | 24 |
|  | Paracingulate cortex | 4.38 | 12 | 42 | 18 |
|  | Left frontal pole | 4.33 | -6 | 57 | 3 |
| 291 | Right angular gyrus | 5.52 | 57 | -54 | 24 |
|  | Right angular gyrus | 4.74 | 51 | -54 | 36 |
|  | Right angular gyrus | 3.76 | 39 | -51 | 21 |
|  | Right angular gyrus | 3.67 | 45 | -45 | 21 |
|  | Right angular gyrus | 3.66 | 54 | -54 | 51 |
|  | Right posterior supramarginal gyrus | 3.66 | 48 | -39 | 21 |
| 132 | Left Crus I | 5.25 | -21 | -78 | -33 |
|  | Left Crus II | 4.65 | -9 | -78 | -33 |
| 94 | Right superior frontal gyrus | 4.13 | 21 | 33 | 39 |
|  | Right middle frontal gyrus | 4.06 | 27 | 33 | 48 |
|  | Right superior frontal gyrus | 3.7 | 27 | 24 | 57 |
|  | Right superior frontal gyrus | 3.62 | 24 | 24 | 51 |
| 61 | Right IX | 5.14 | 3 | -45 | -42 |
|  | Left IX | 4.22 | -6 | -51 | -33 |
| 47 | Right middle frontal gyrus | 4.87 | 39 | 9 | 45 |
|  | Right middle frontal gyrus | 3.95 | 36 | 12 | 33 |
|  | Right middle frontal gyrus | 3.36 | 42 | 24 | 33 |
|  | Right middle frontal gyrus | 3.25 | 45 | 24 | 42 |

**Table B.10.** Local maxima cluster index when contrasting Blinks > Urge blocks.

| Cluster Size | Region | Z-score | MNI Coordinates |  |  |
| --- | --- | --- | --- | --- | --- |
|  |  |  | X | Y | Z |
| 4149 | Left lingual gyrus | 6.18 | -21 | -51 | -3 |
|  | Left intracalcerine cortex | 6.05 | -12 | -63 | 6 |
|  | Left intracalcerine cortex | 6.05 | -12 | -72 | 12 |
|  | Right lingual gyrus | 5.98 | 15 | -45 | -3 |
|  | Left lingual gyrus | 5.77 | -9 | -57 | 0 |
|  | Right precuneous cortex | 5.74 | 18 | -60 | 12 |
| 538 | Left superior frontal gyrus | 4.97 | -12 | -3 | 66 |
|  | Anterior cingulate cortex | 4.52 | -6 | 12 | 39 |
|  | Left superior frontal gyrus | 4.45 | -18 | 6 | 72 |
|  | Anterior cingulate cortex | 4.42 | 3 | 12 | 39 |
|  | Anterior cingulate cortex | 4.36 | 3 | 18 | 36 |
|  | Right superior frontal gyrus | 4.29 | 12 | 0 | 63 |
| 502 | Left central opercular cortex | 5.5 | -45 | 6 | 6 |
|  | Left precentral gyrus | 5.45 | -57 | 6 | 3 |
|  | Left frontal opercular cortex | 5.15 | -45 | 12 | 0 |
|  | Left postcentral gyrus | 5.12 | -57 | -24 | 21 |
|  | Left frontal opercular cortex | 4.92 | -36 | 12 | 15 |
|  | Left insular cortex | 4.6 | -36 | 9 | 3 |
| 201 | Right insular cortex | 4.73 | 36 | 9 | 9 |
|  | Right central opercular cortex | 4.24 | 54 | -6 | 15 |
|  | Right precentral gyrus | 4.03 | 57 | 6 | 6 |
|  | Right central opercular cortex | 3.94 | 48 | 6 | 6 |
|  | Right precentral gyrus | 3.84 | 51 | 3 | 12 |
|  | Right central opercular cortex | 3.66 | 42 | -12 | 21 |
| 138 | Left frontal pole | 5.07 | -30 | 51 | 30 |
|  | Left frontal pole | 4.73 | -30 | 45 | 21 |
|  | Left frontal pole | 4.59 | -42 | 48 | 27 |
|  | Left frontal pole | 4.22 | -24 | 60 | 33 |
| 54 | Left postcentral gyrus | 4.71 | -36 | -18 | 39 |
|  | Left precentral gyrus | 4.68 | -42 | -9 | 51 |
| 37 | Right frontal pole | 4.43 | 36 | 51 | 21 |
|  | Right frontal pole | 3.89 | 30 | 48 | 36 |
|  | Right frontal pole | 3.68 | 27 | 45 | 30 |

## Appendix C: Activation Time Series

**Figure C.1.**
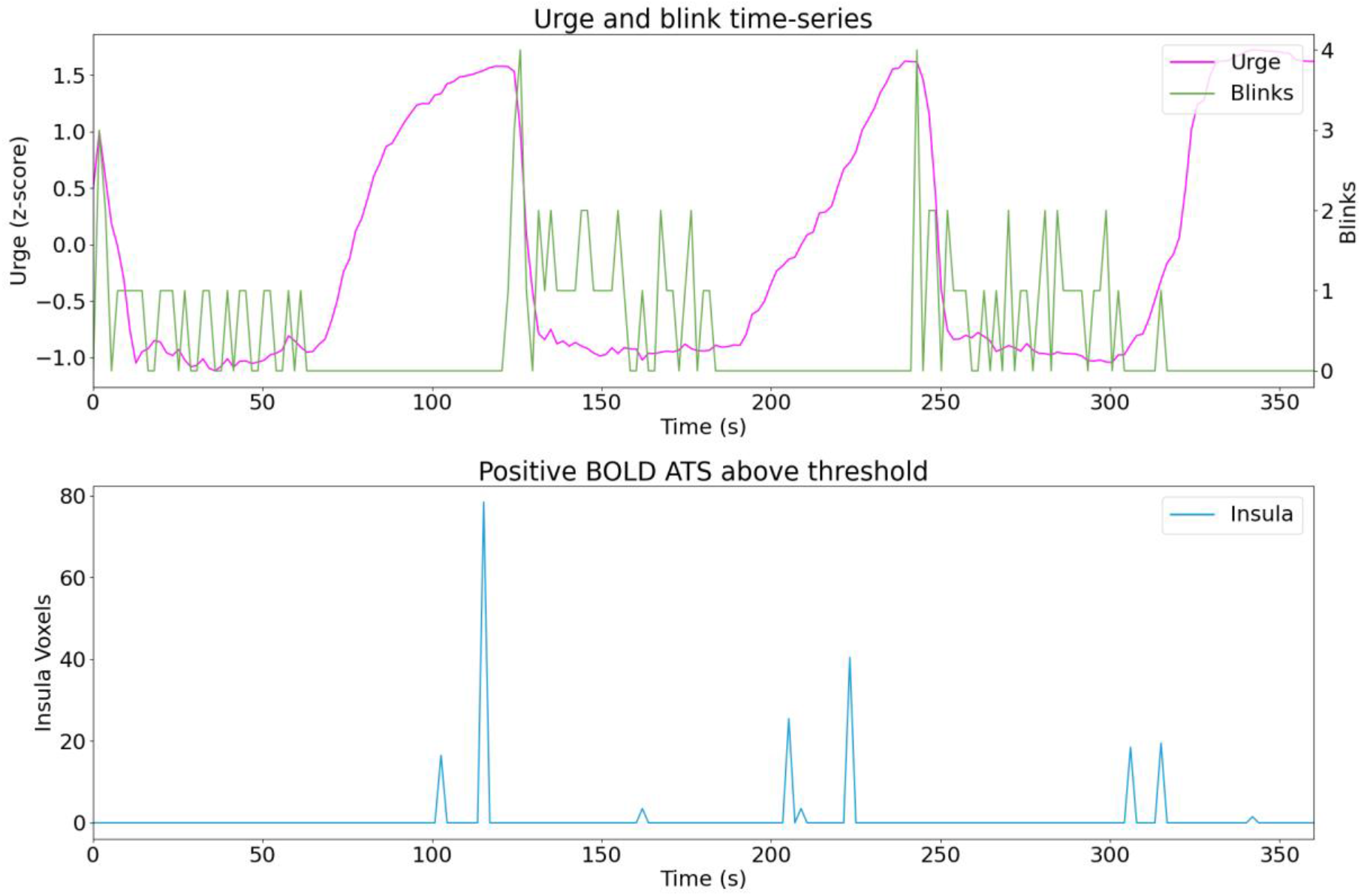
Sub01 run01.

**Figure C.2.**
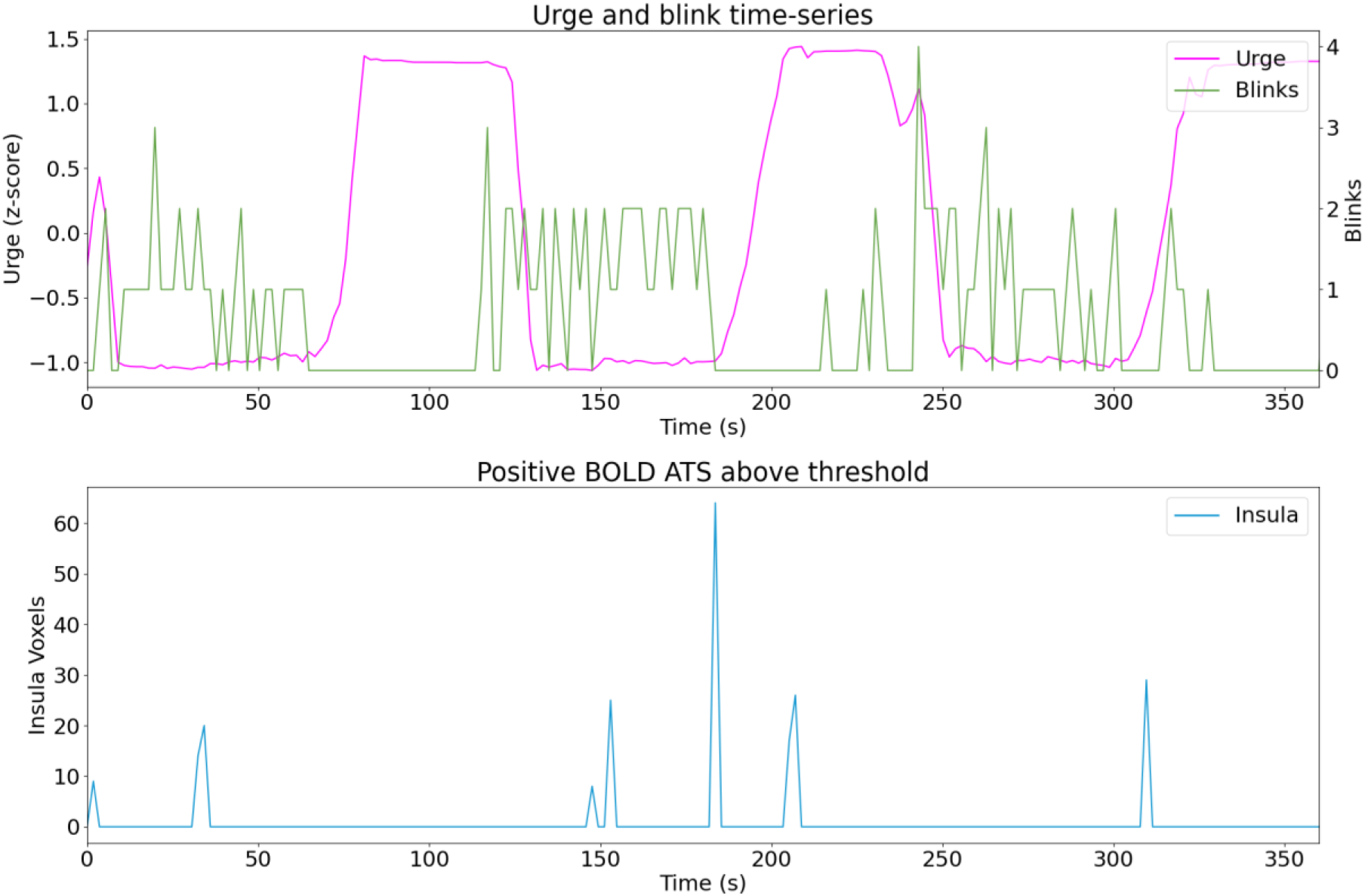
Sub01 run02.

**Figure C.3.**
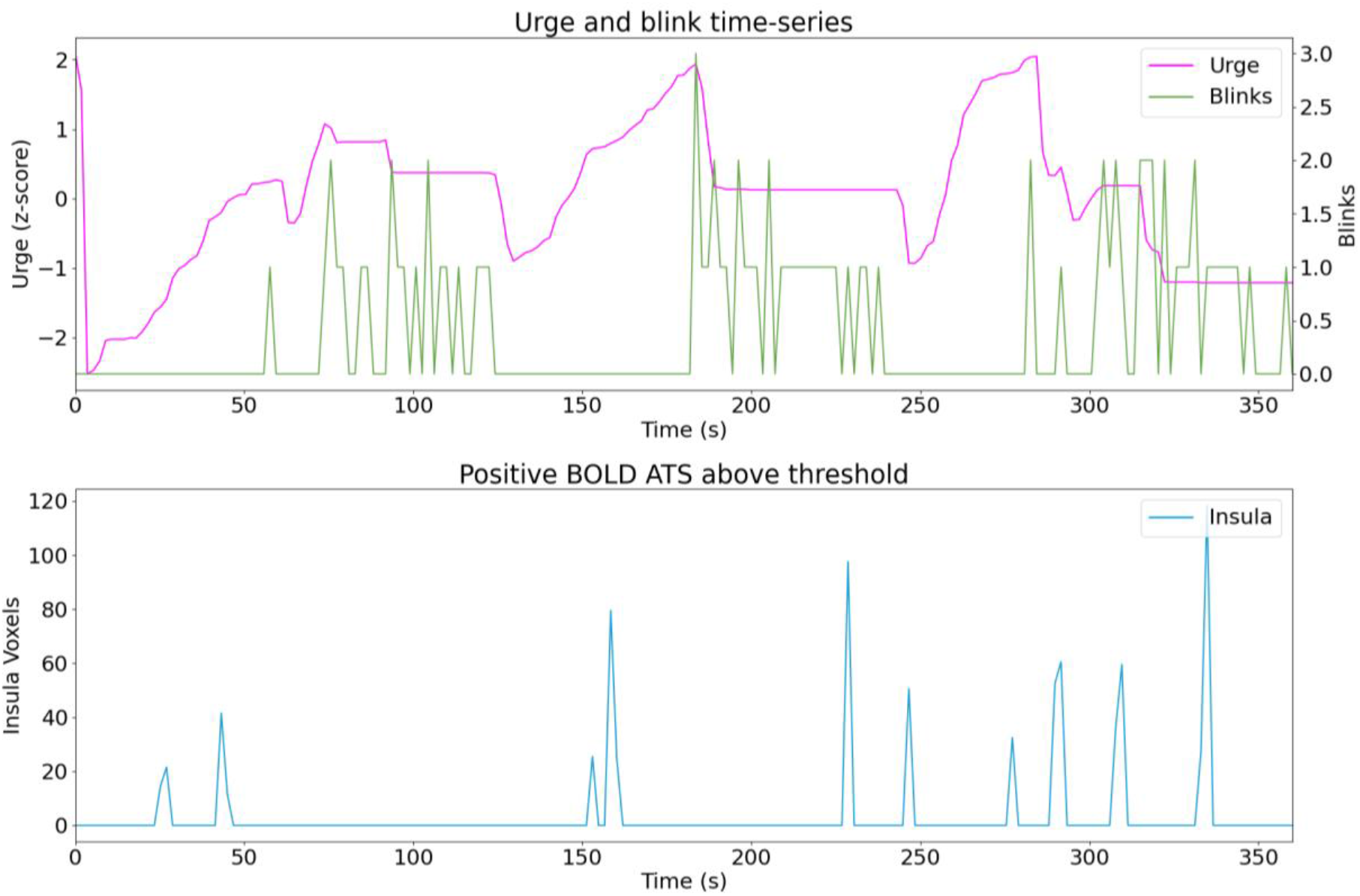
Sub03 run01.

**Figure C.4.**
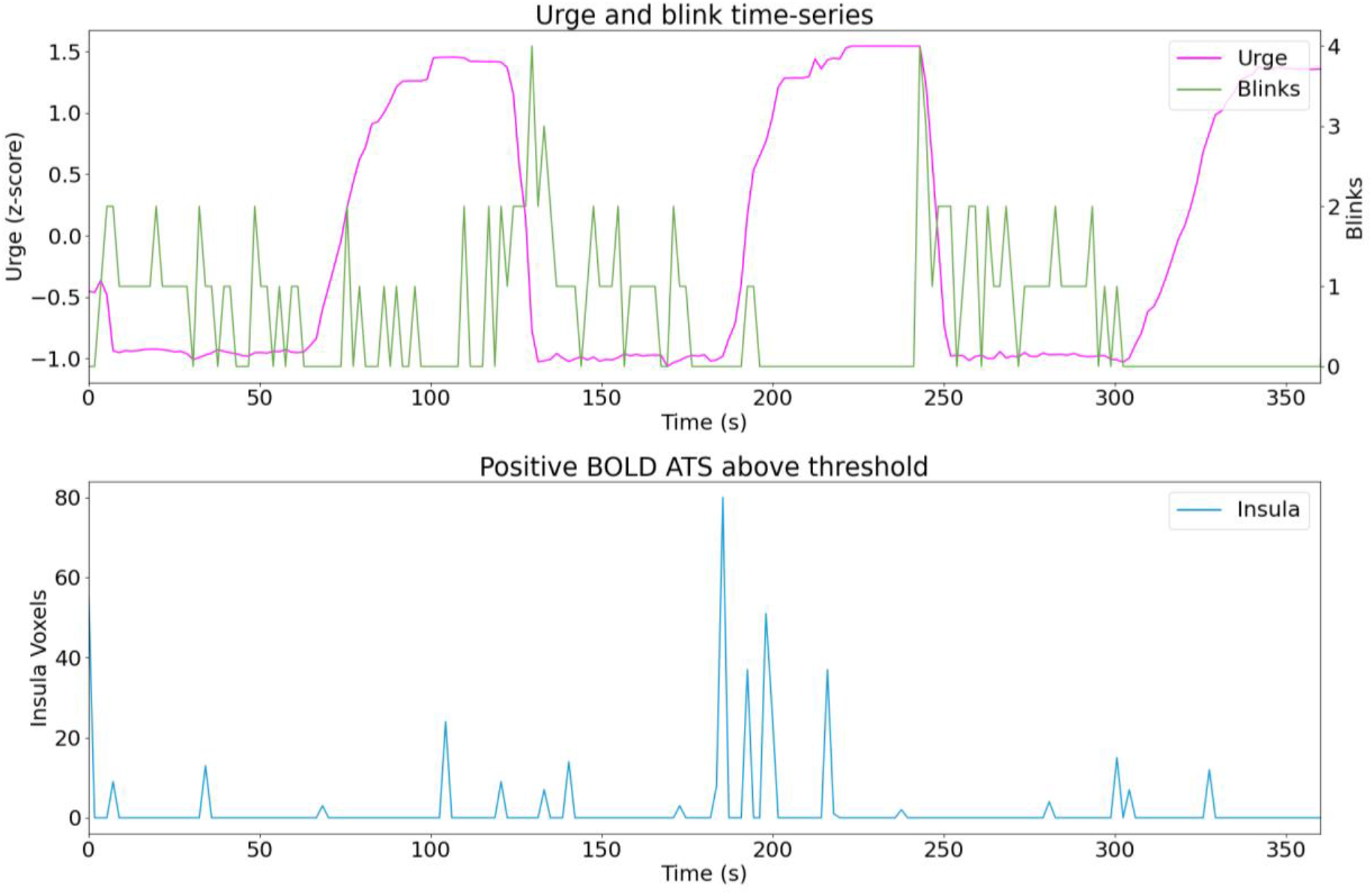
Sub01 run03.

**Figure C.5.**
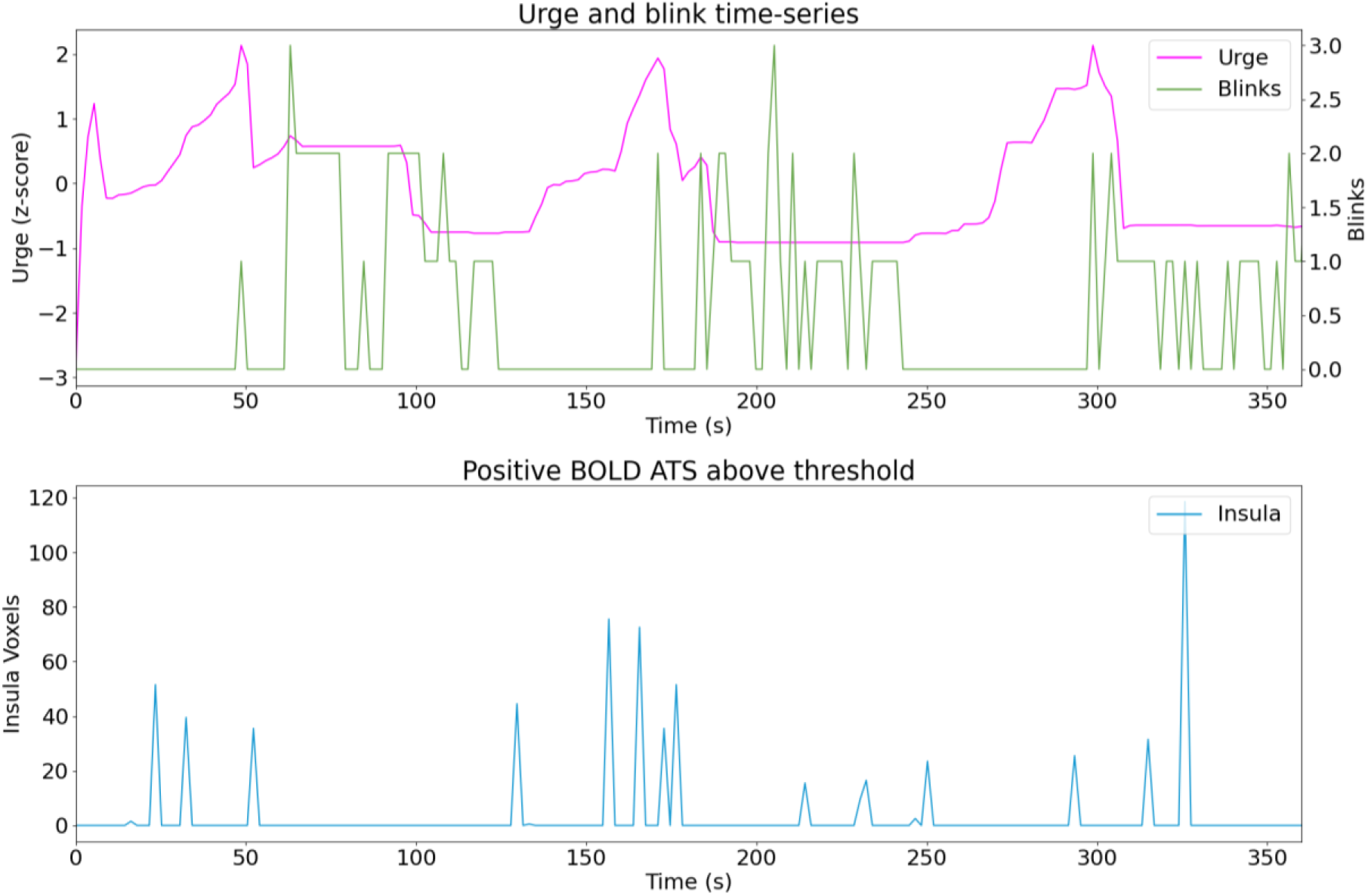
Sub03 run03.

**Figure C.6.**
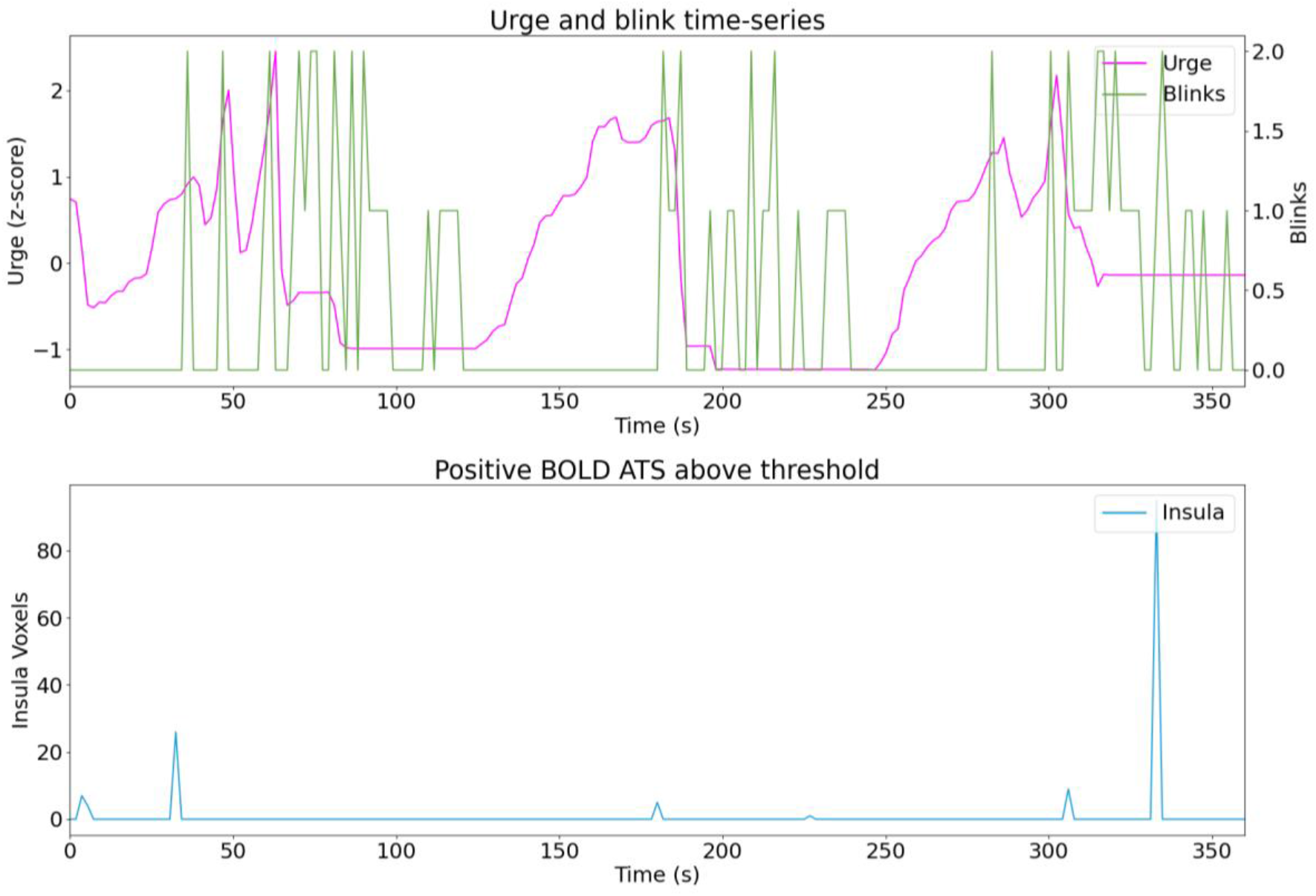
Sub03 run02.

**Figure C.7.**
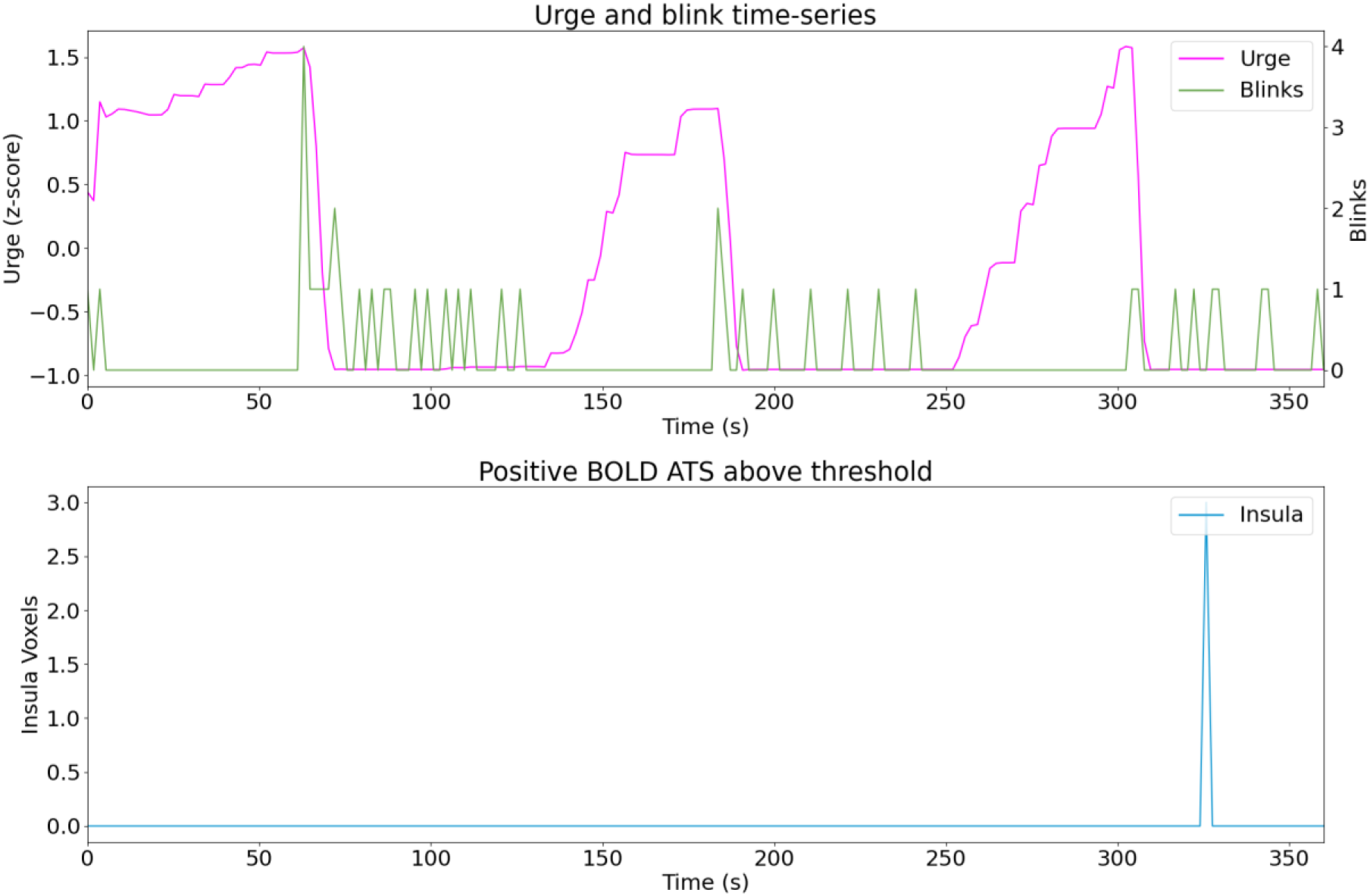
Sub04 run01.

**Figure C.8.**
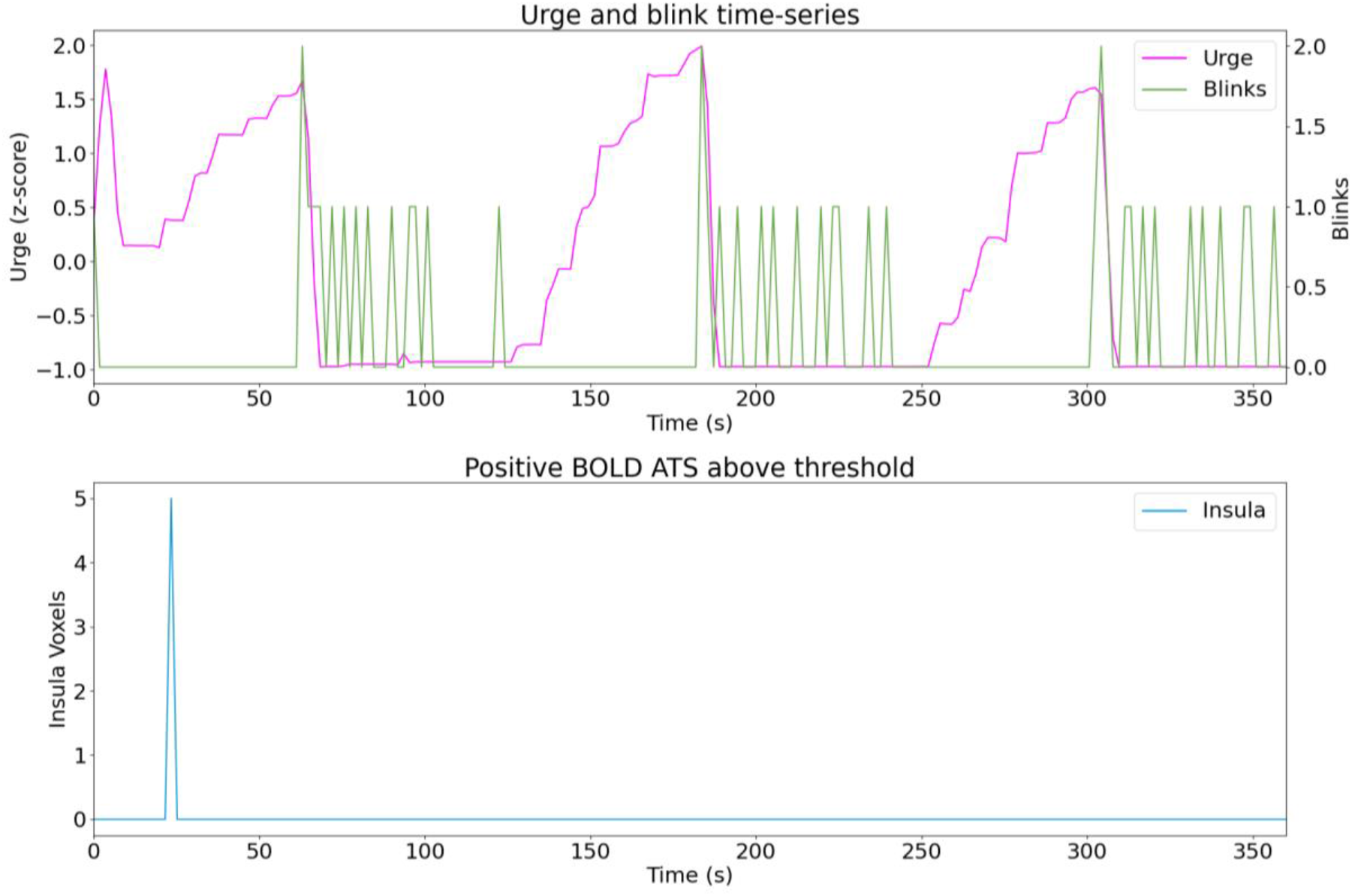
Sub04 run02.

**Figure C.9.**
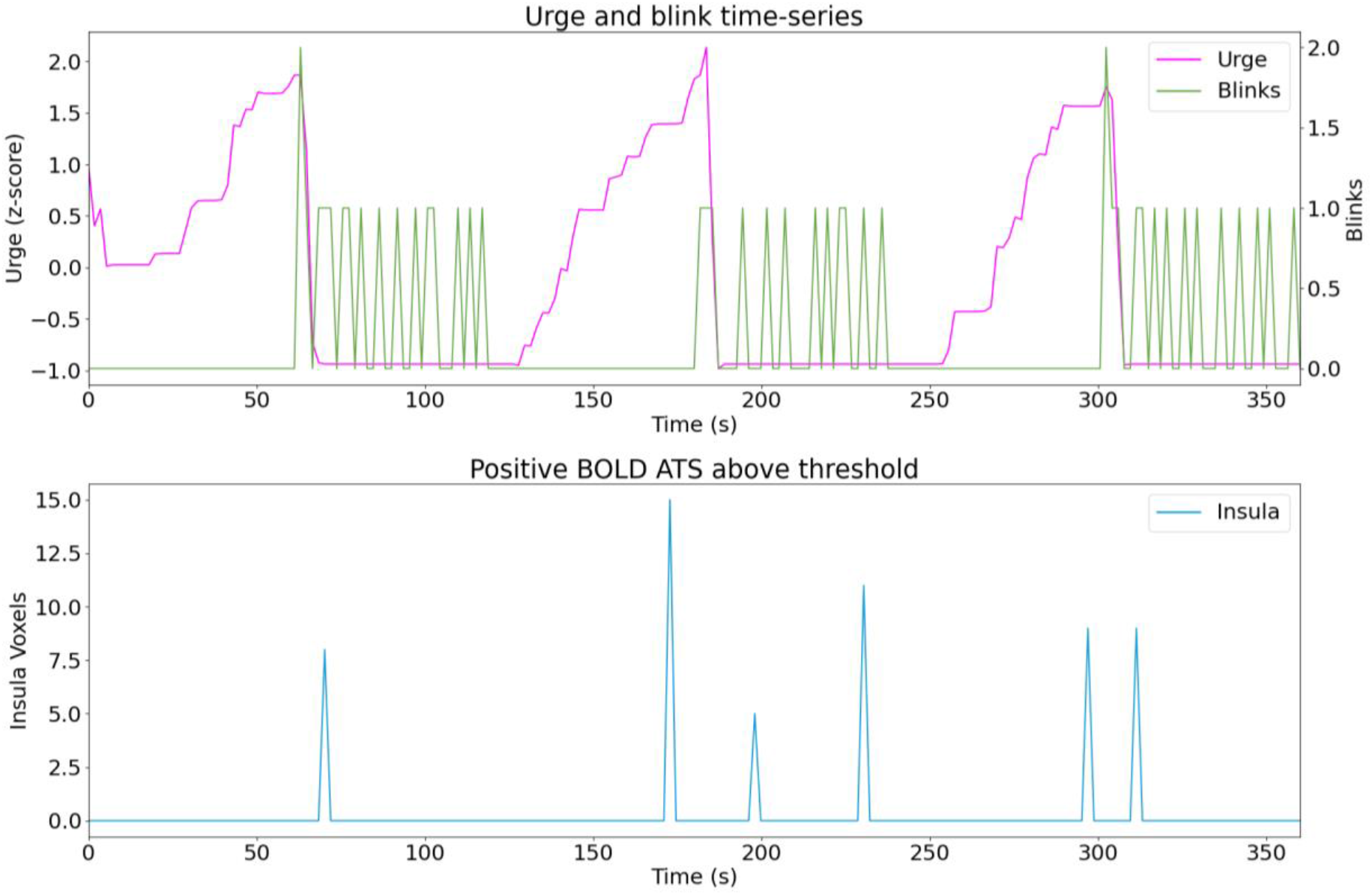
Sub04 run03.

**Figure C.10.**
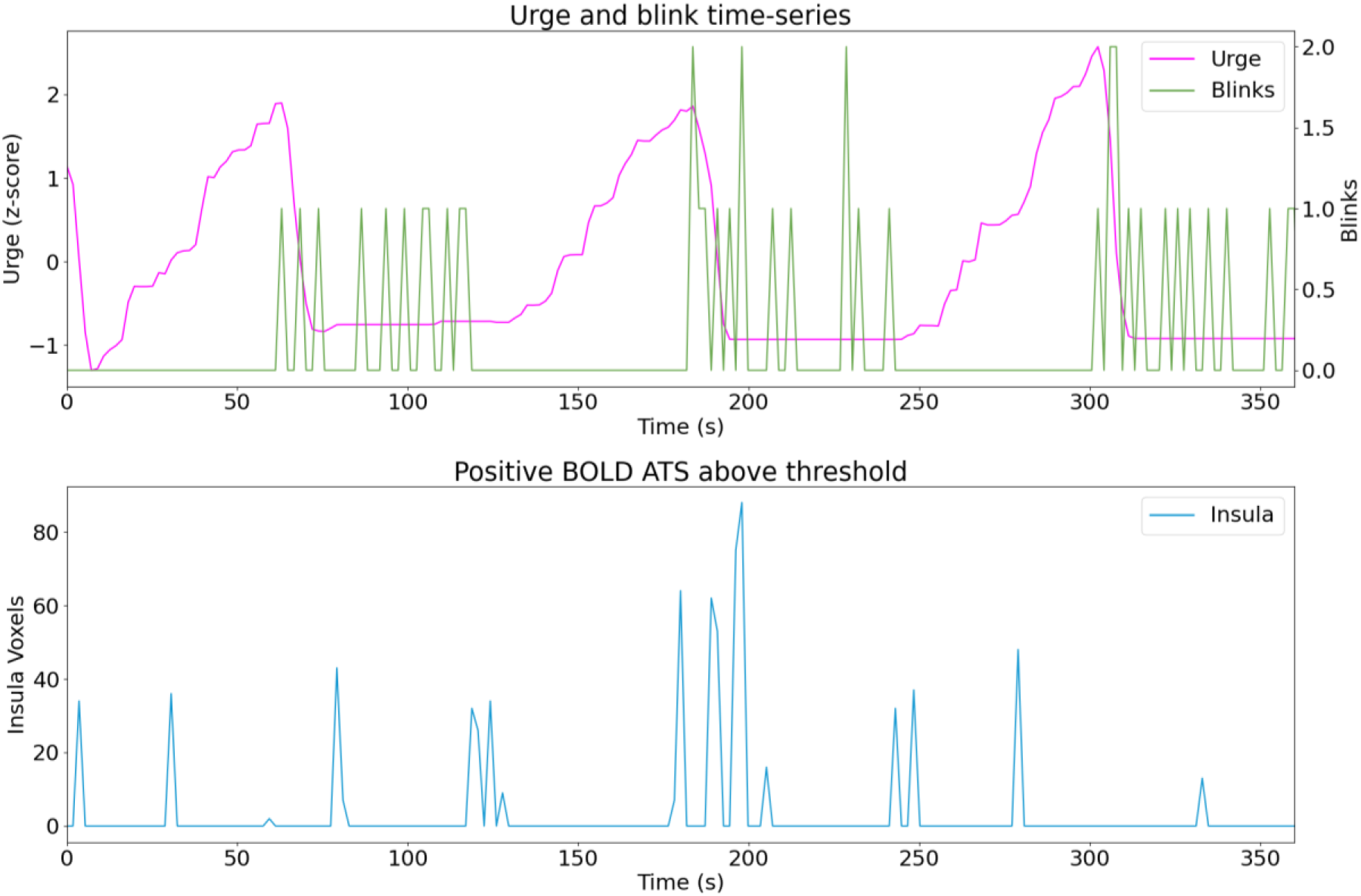
Sub05 run02.

**Figure C.11.**
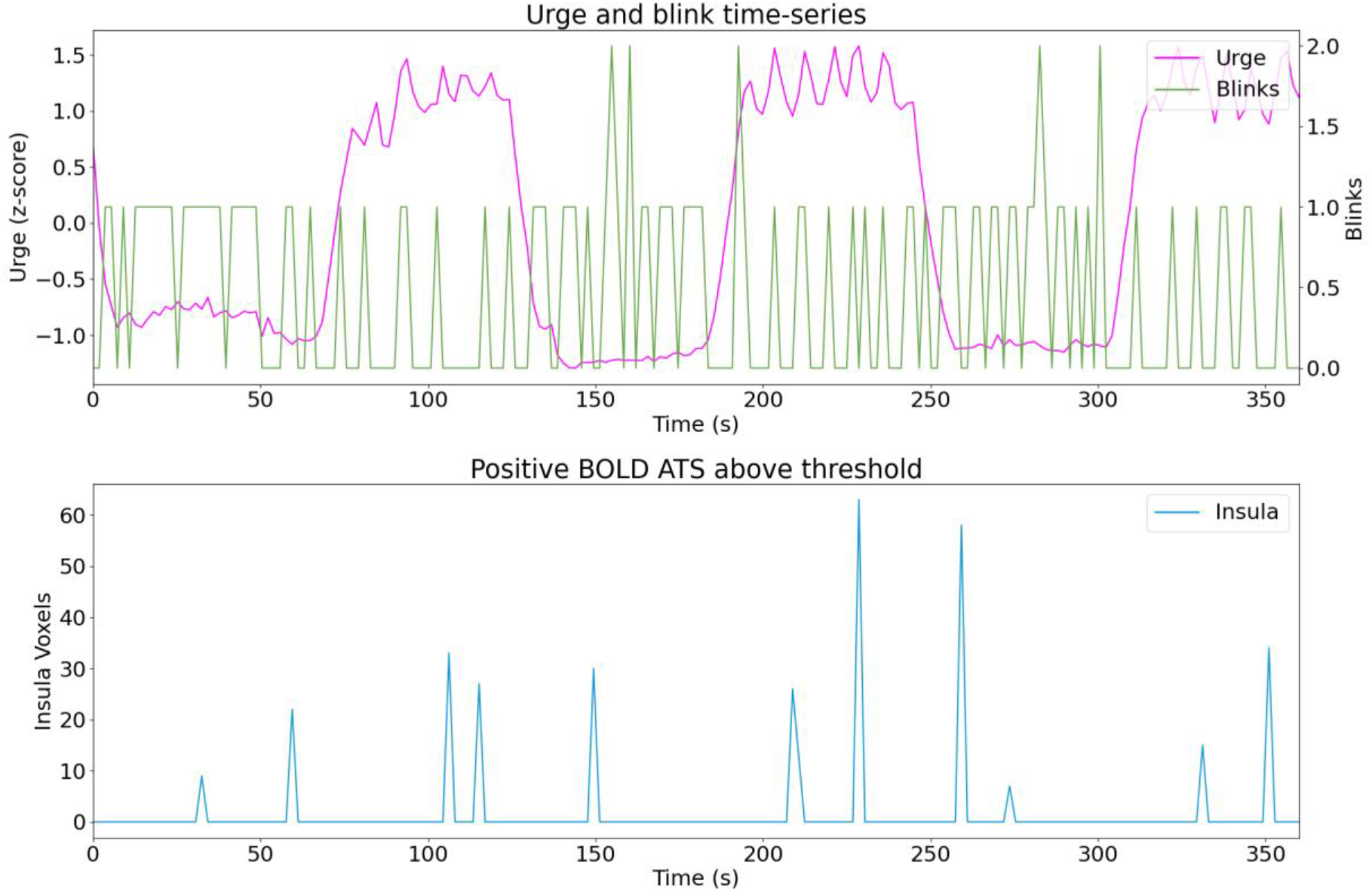
Sub06 run02.

**Figure C.12.**
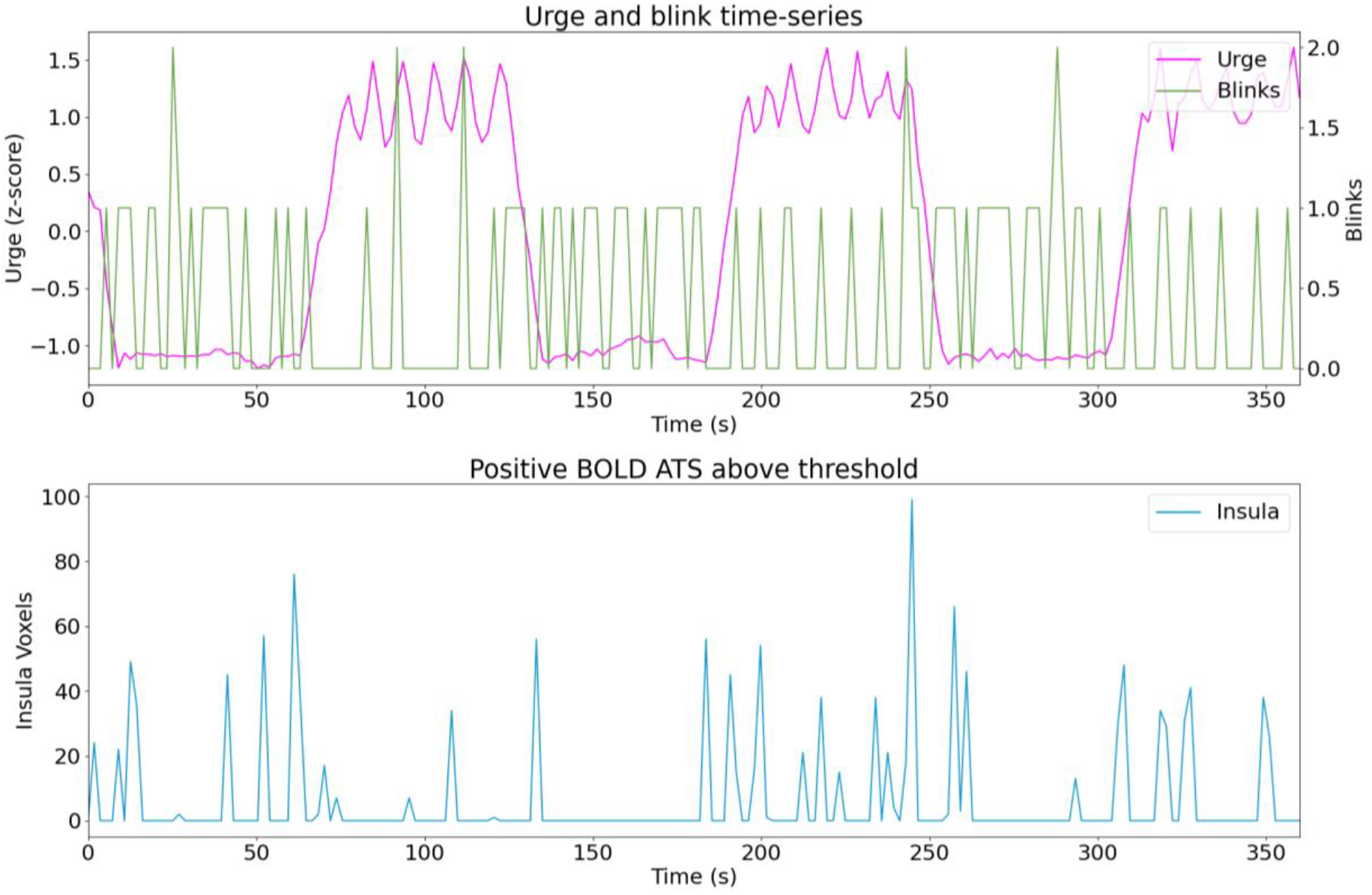
Sub06 run03.

**Figure C.13.**
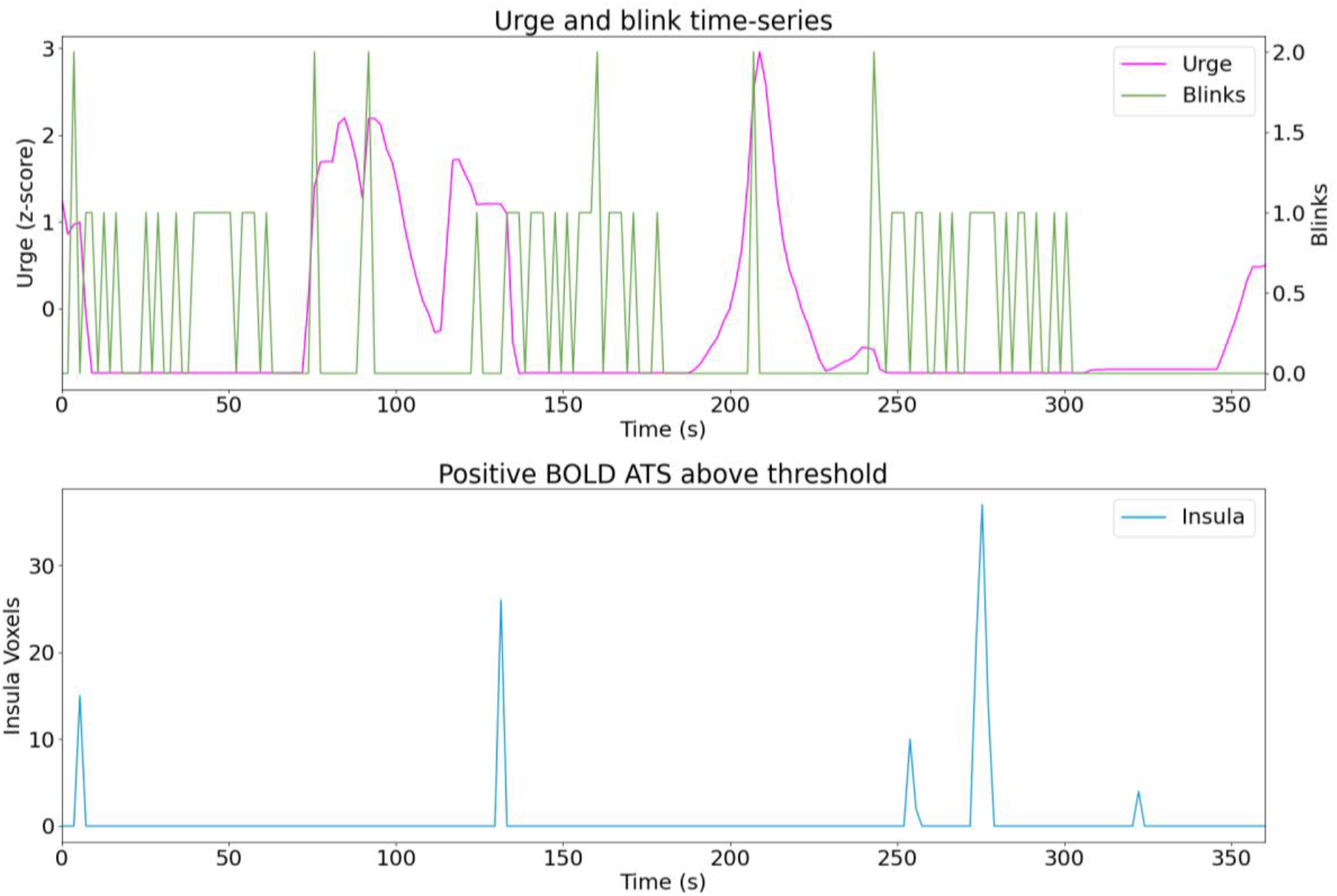
Sub07 run01.

**Figure C.14.**
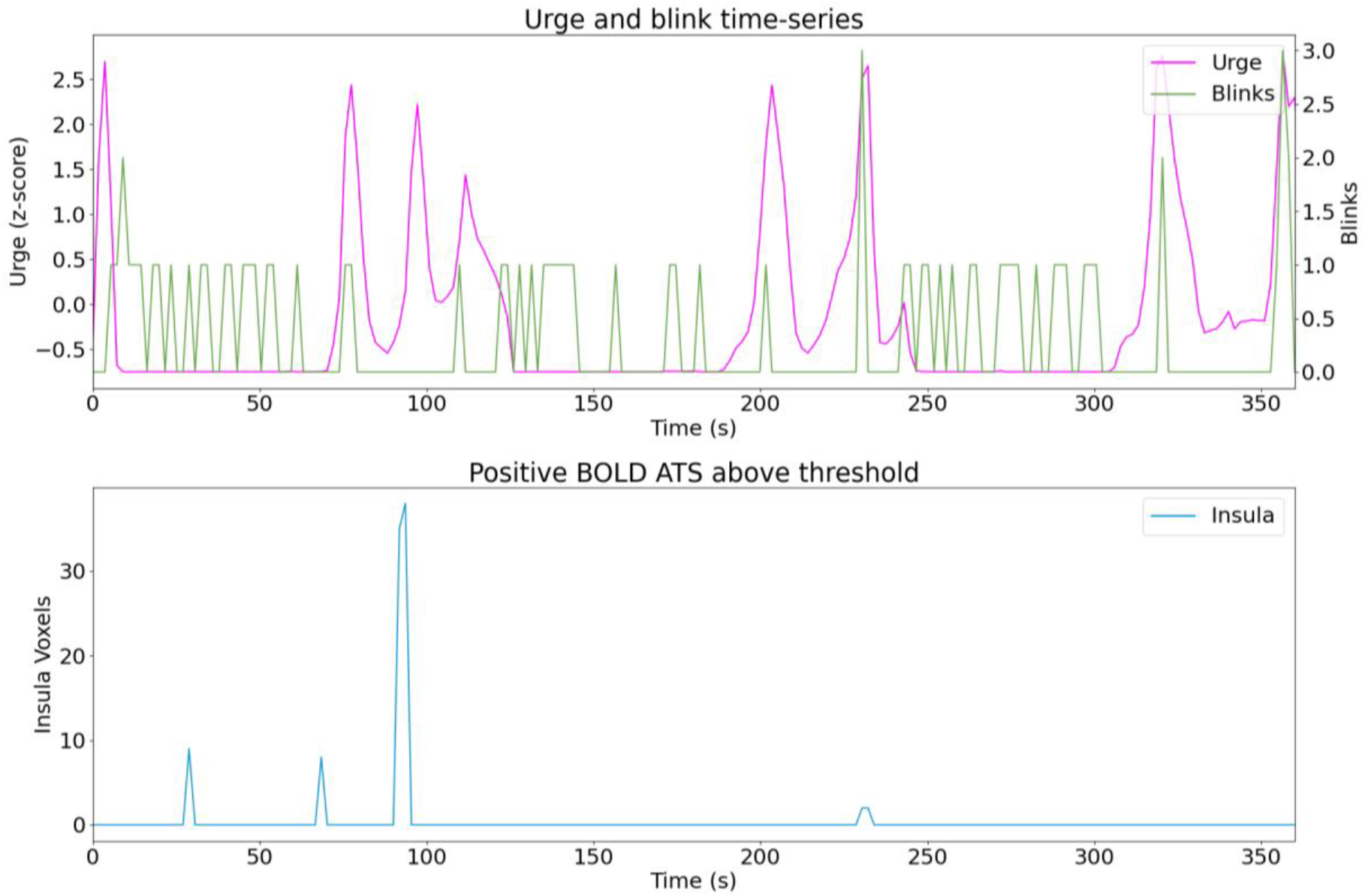
Sub07 run02.

**Figure C.15.**
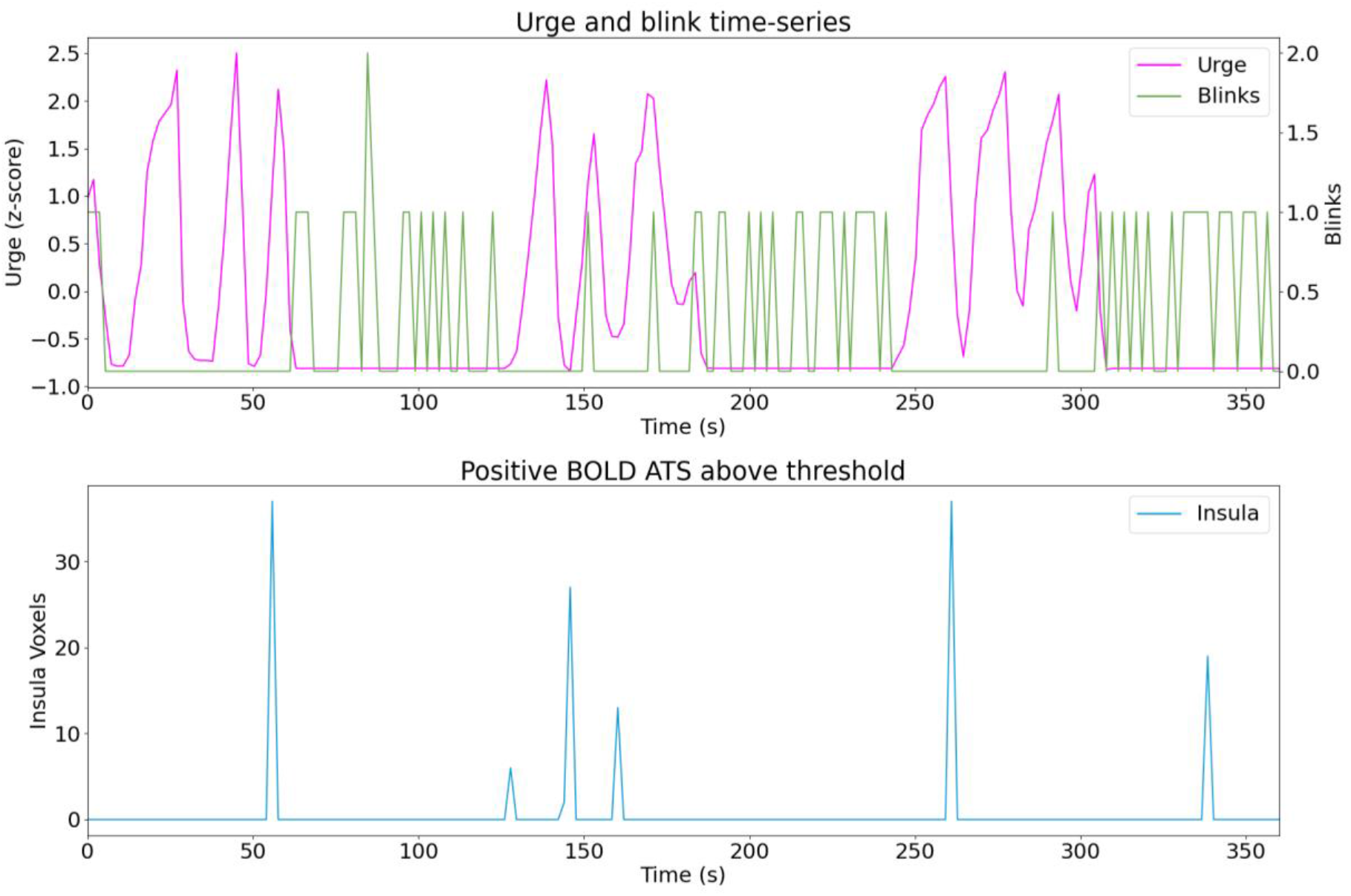
Sub09 run01.

**Figure C.16.**
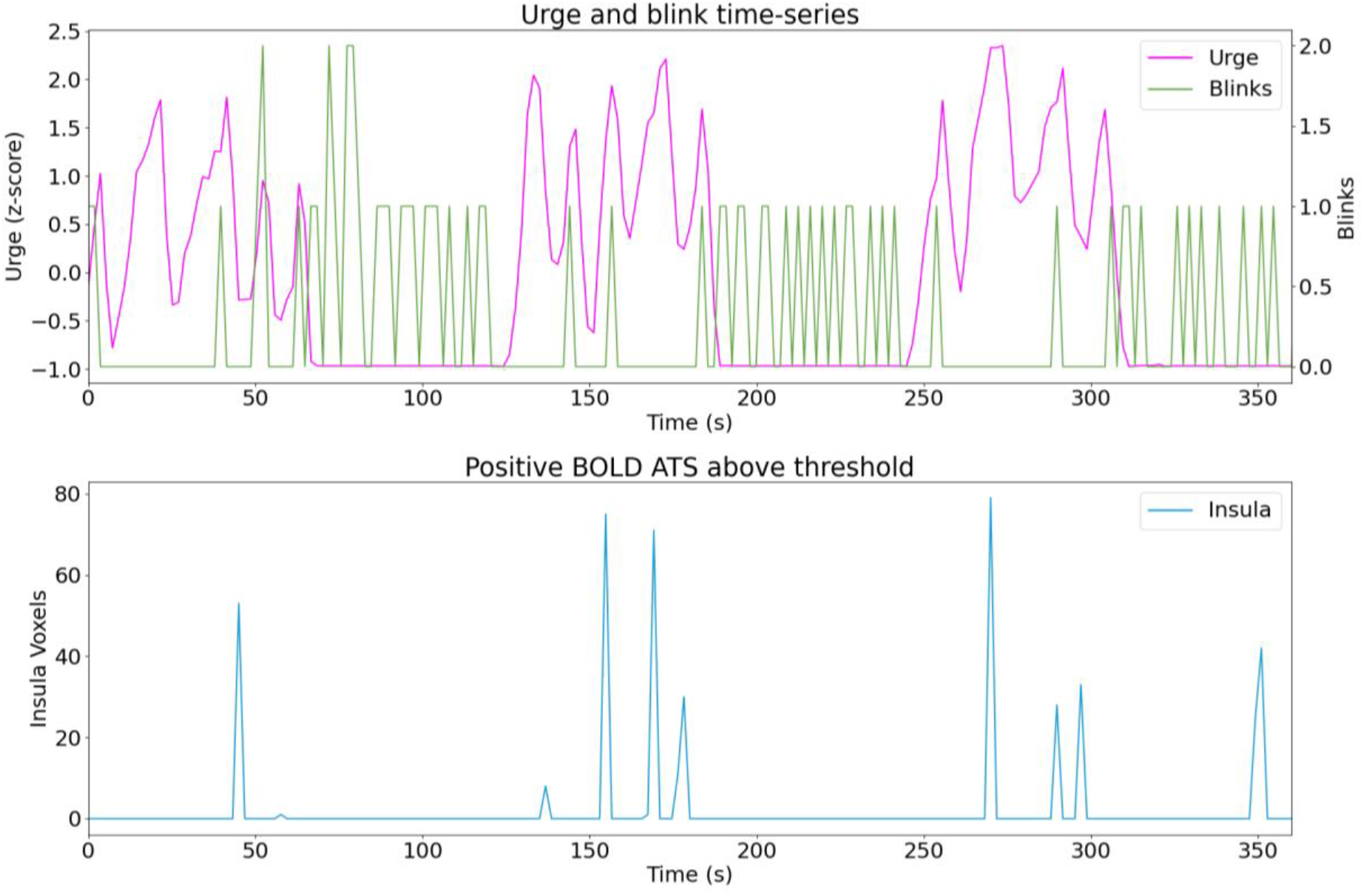
Sub09 run02.

**Figure C.17.**
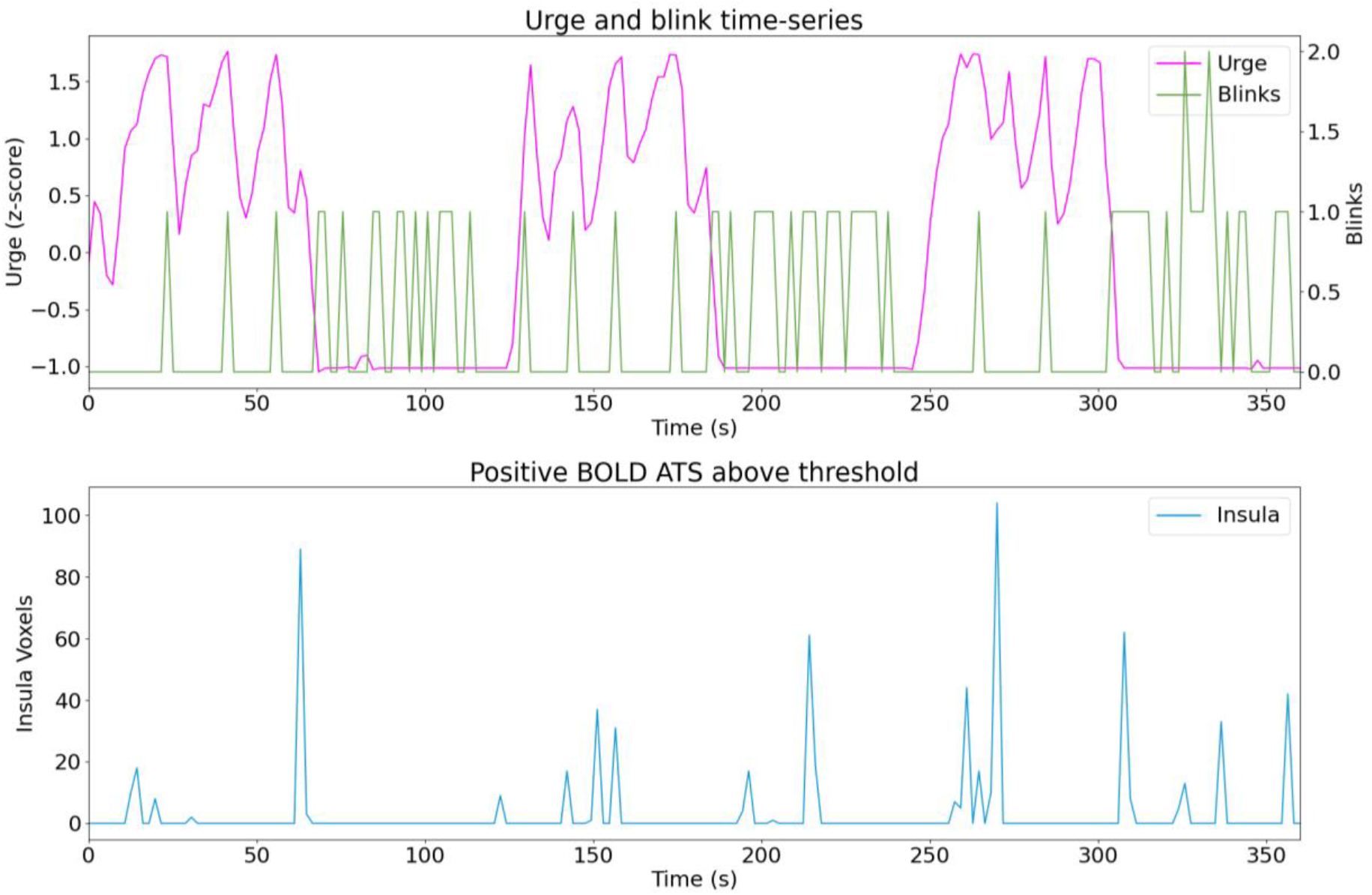
Sub09 run03.

**Figure C.18.**
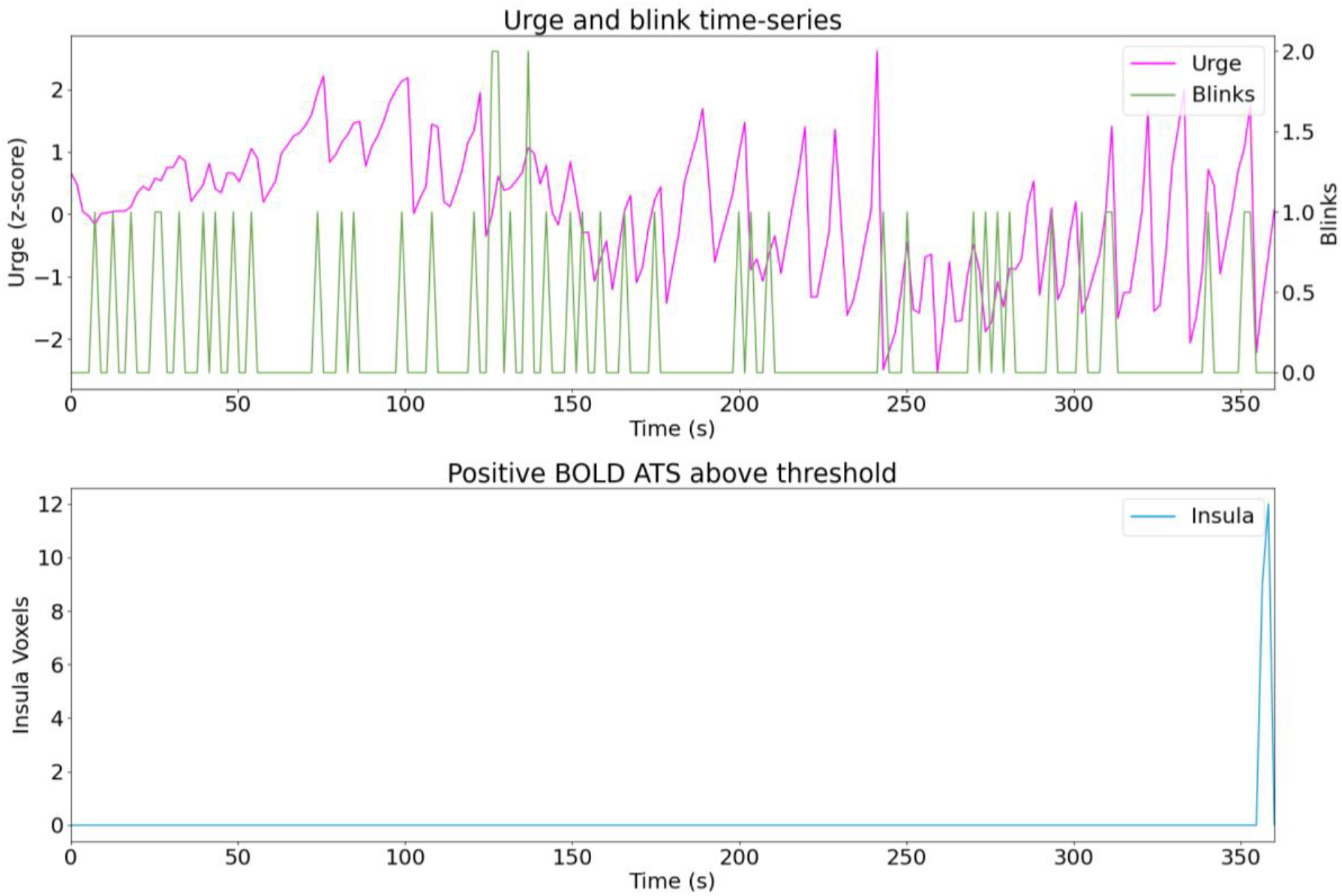
Sub10 run01.

**Figure C.19.**
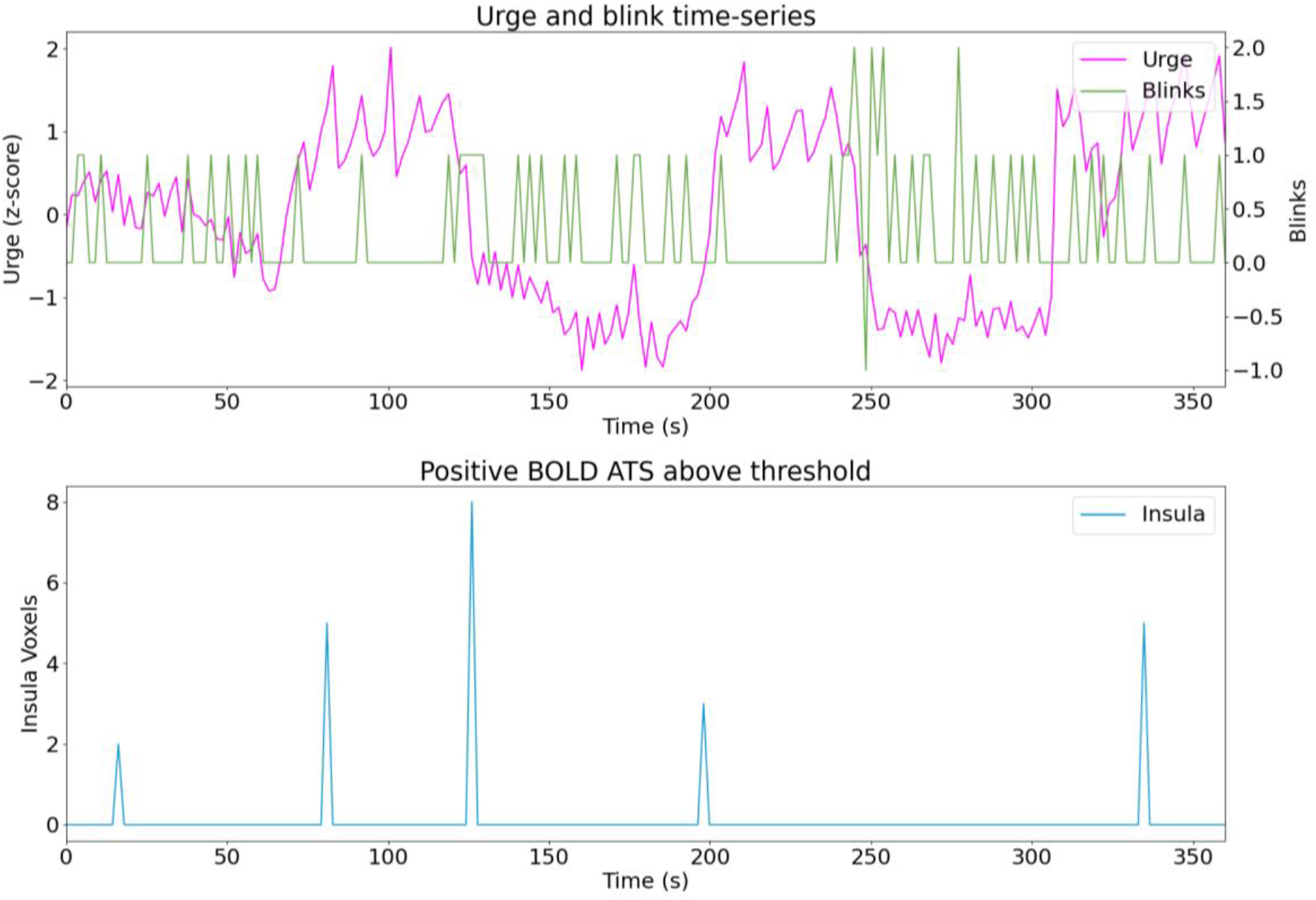
Sub10 run02.

**Figure C.20.**
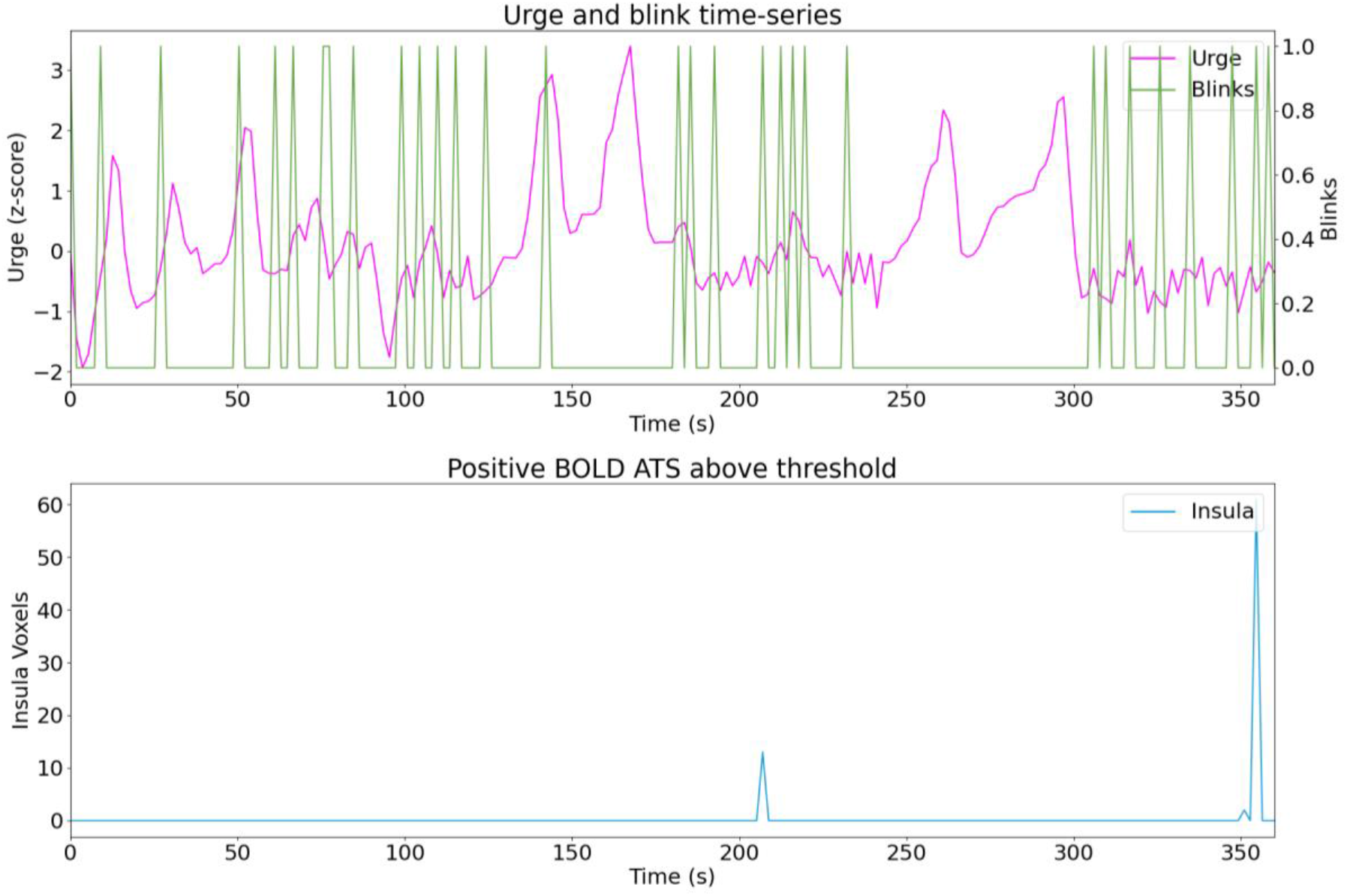
Sub11 run01.

**Figure C.21.**
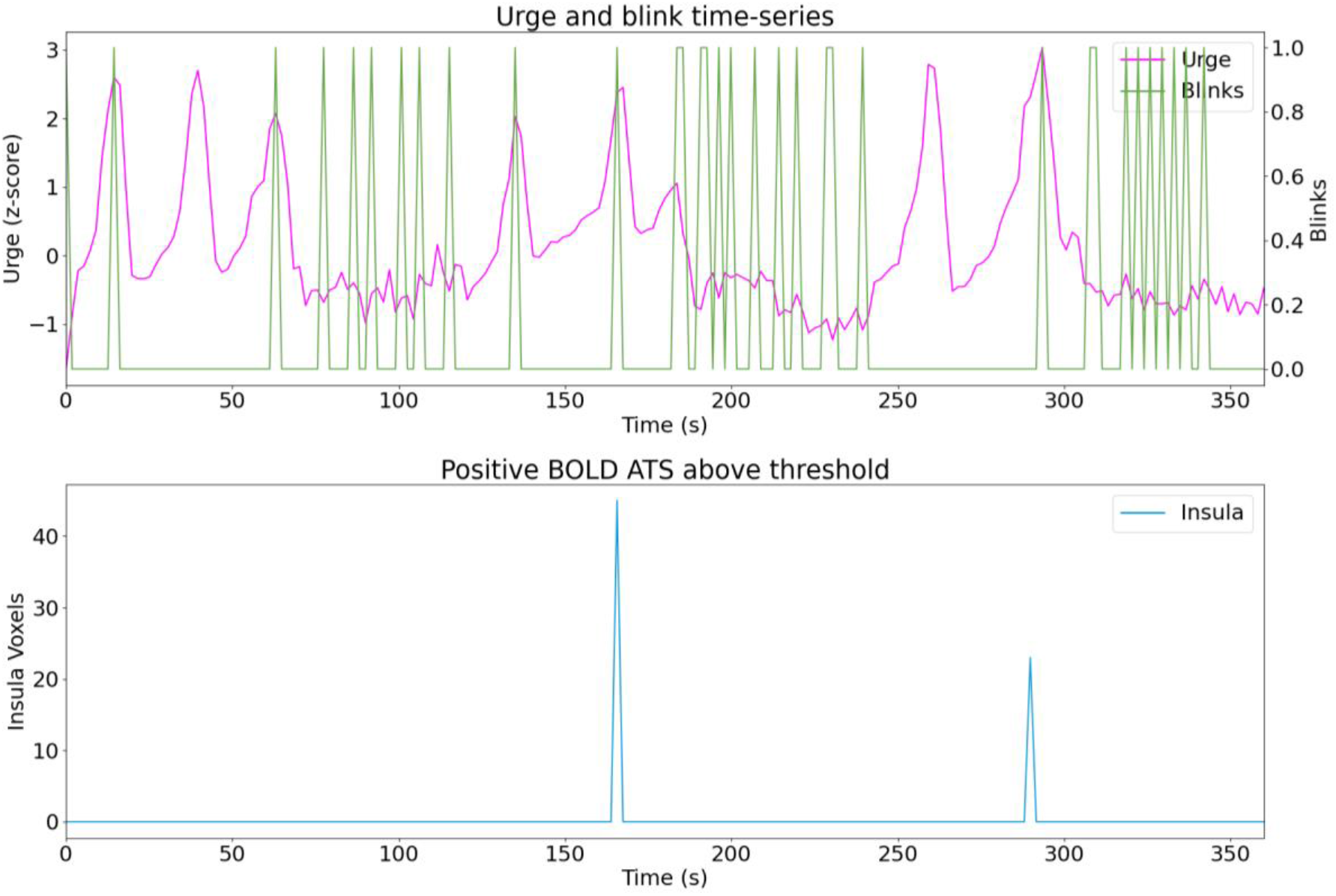
Sub11 run02.

**Figure C.22.**
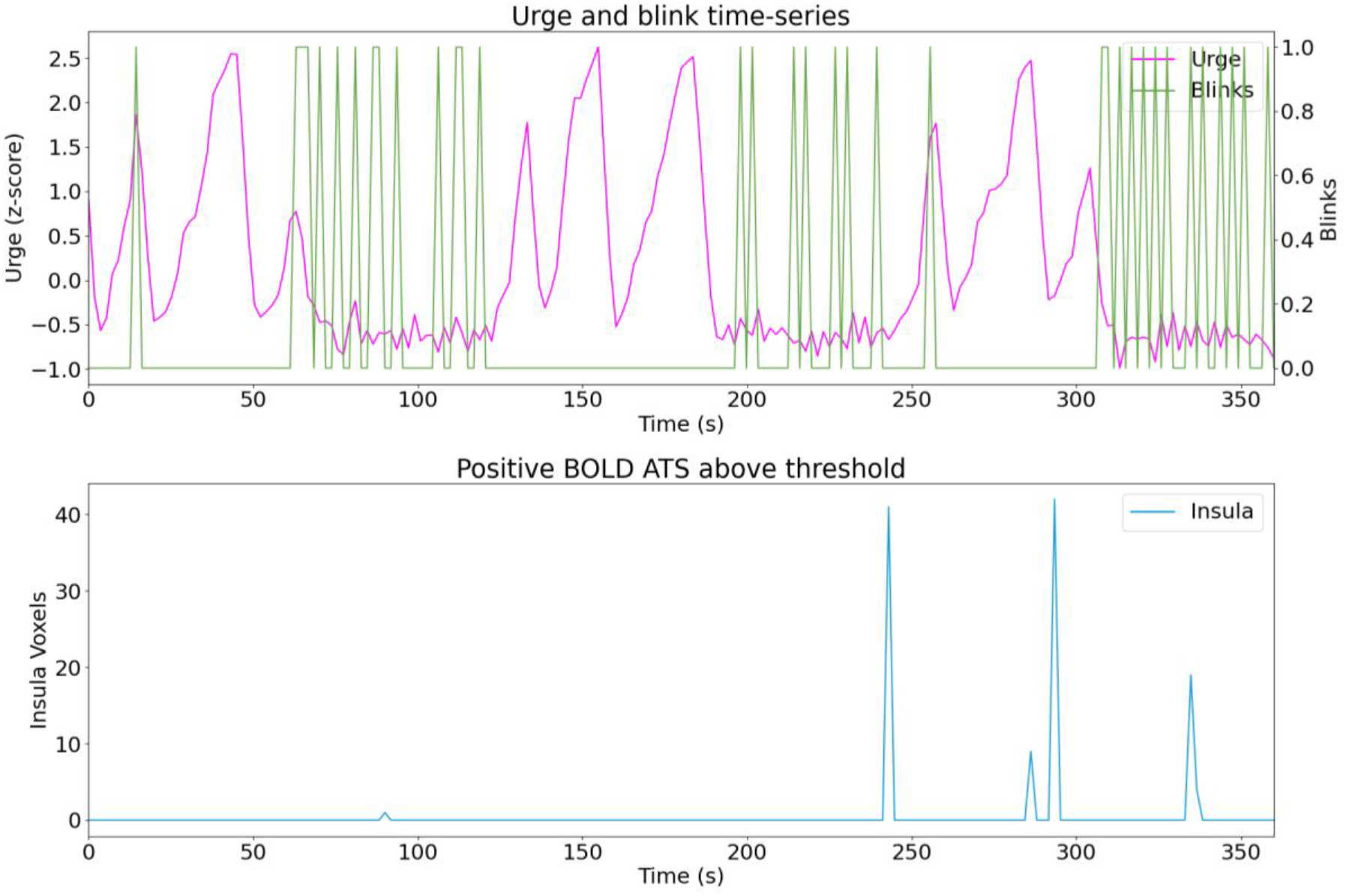
Sub11 run03.

**Figure C.23.**
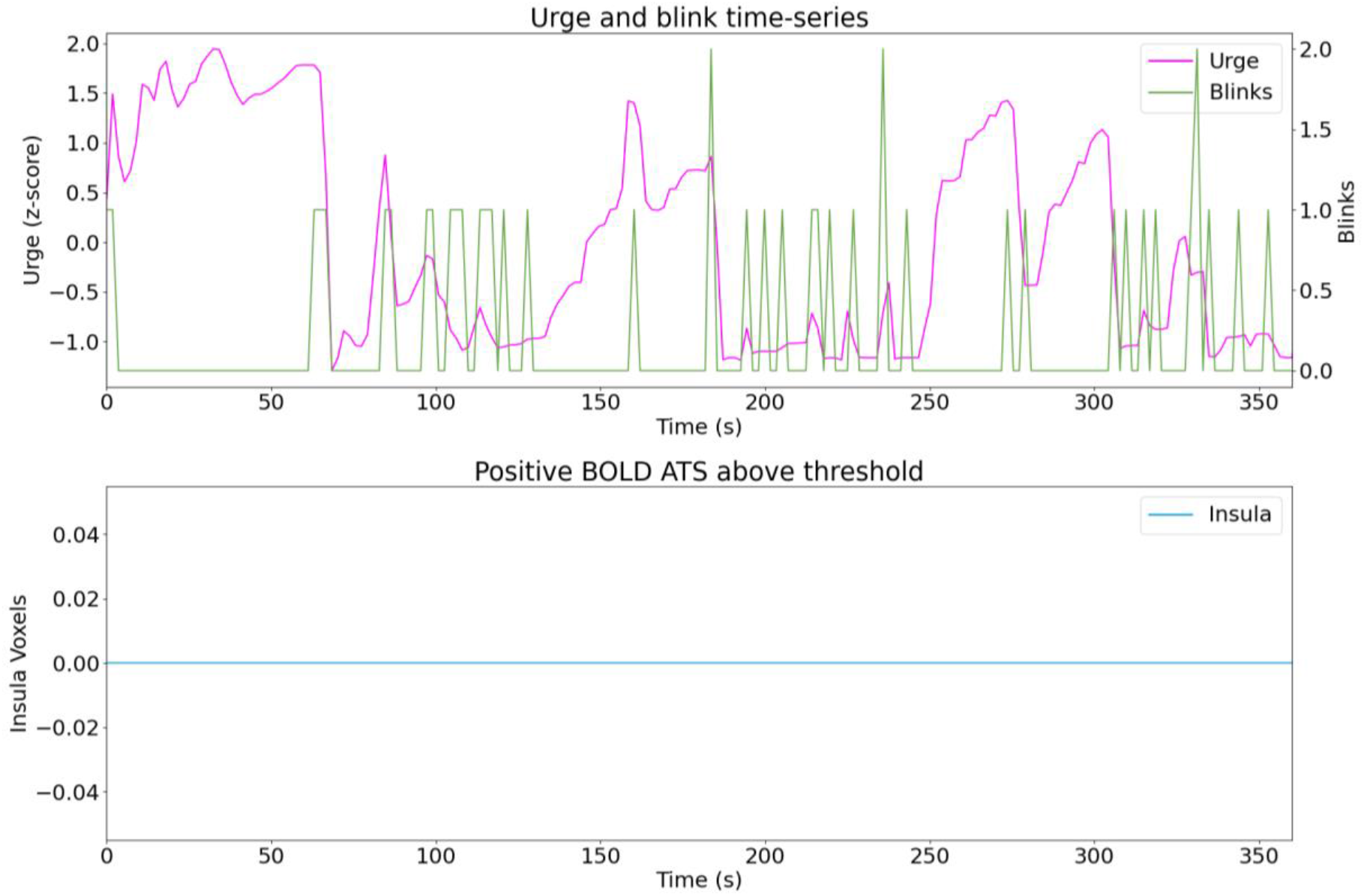
Sub12 run01.

**Figure C.24.**
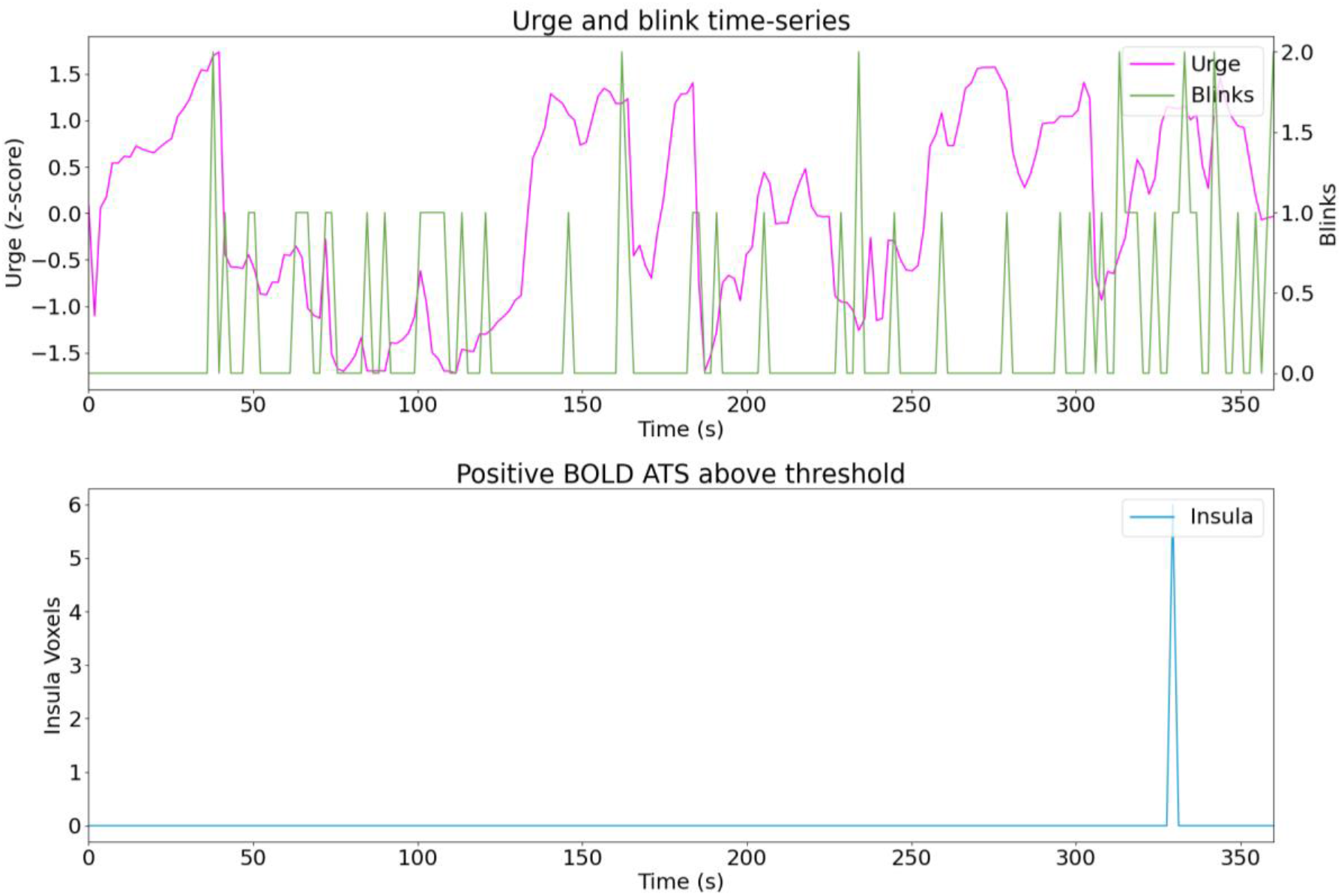
Sub12 run02.

**Figure C.25.**
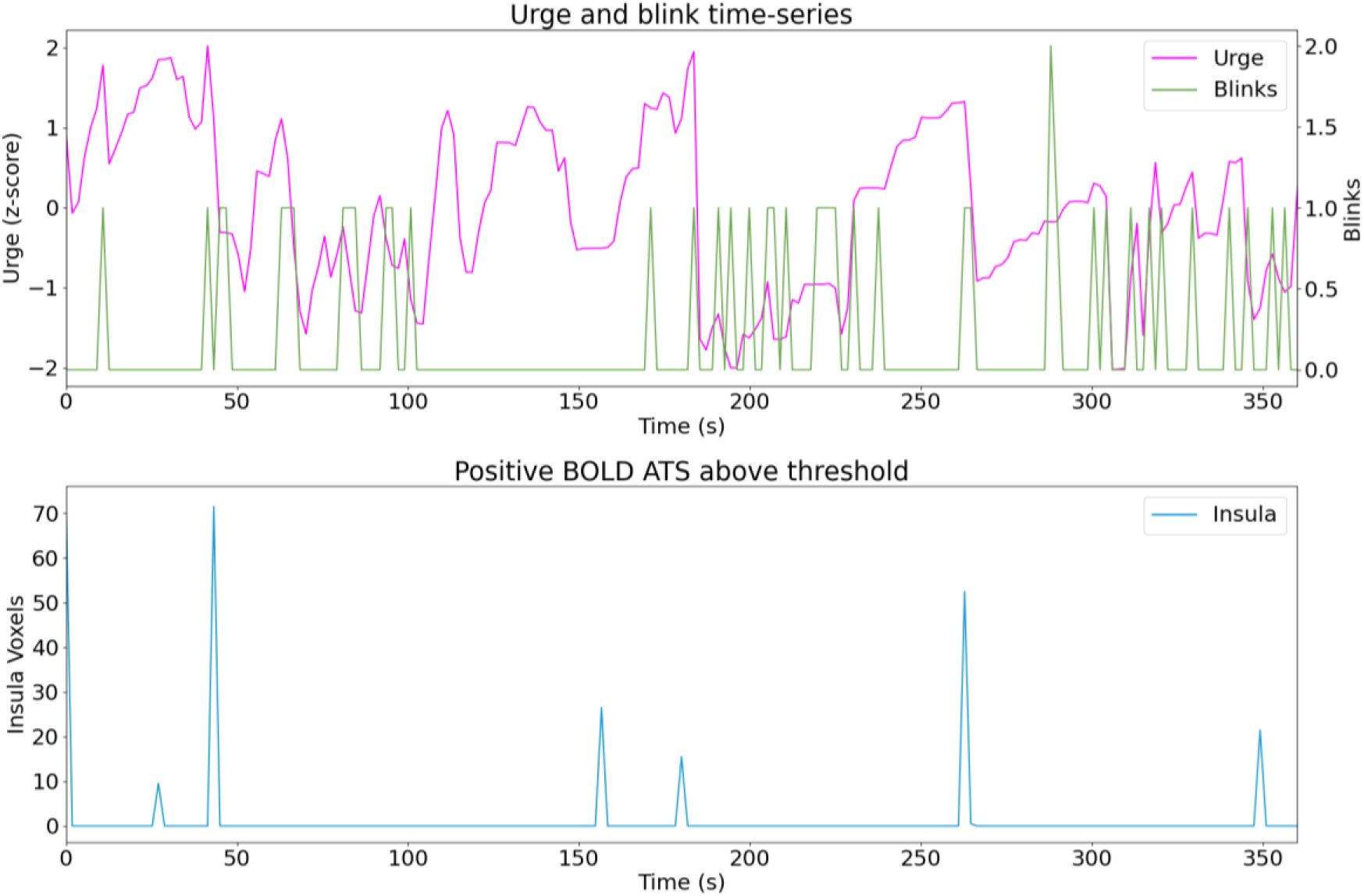
Sub12 run03.

**Figure C.26.**
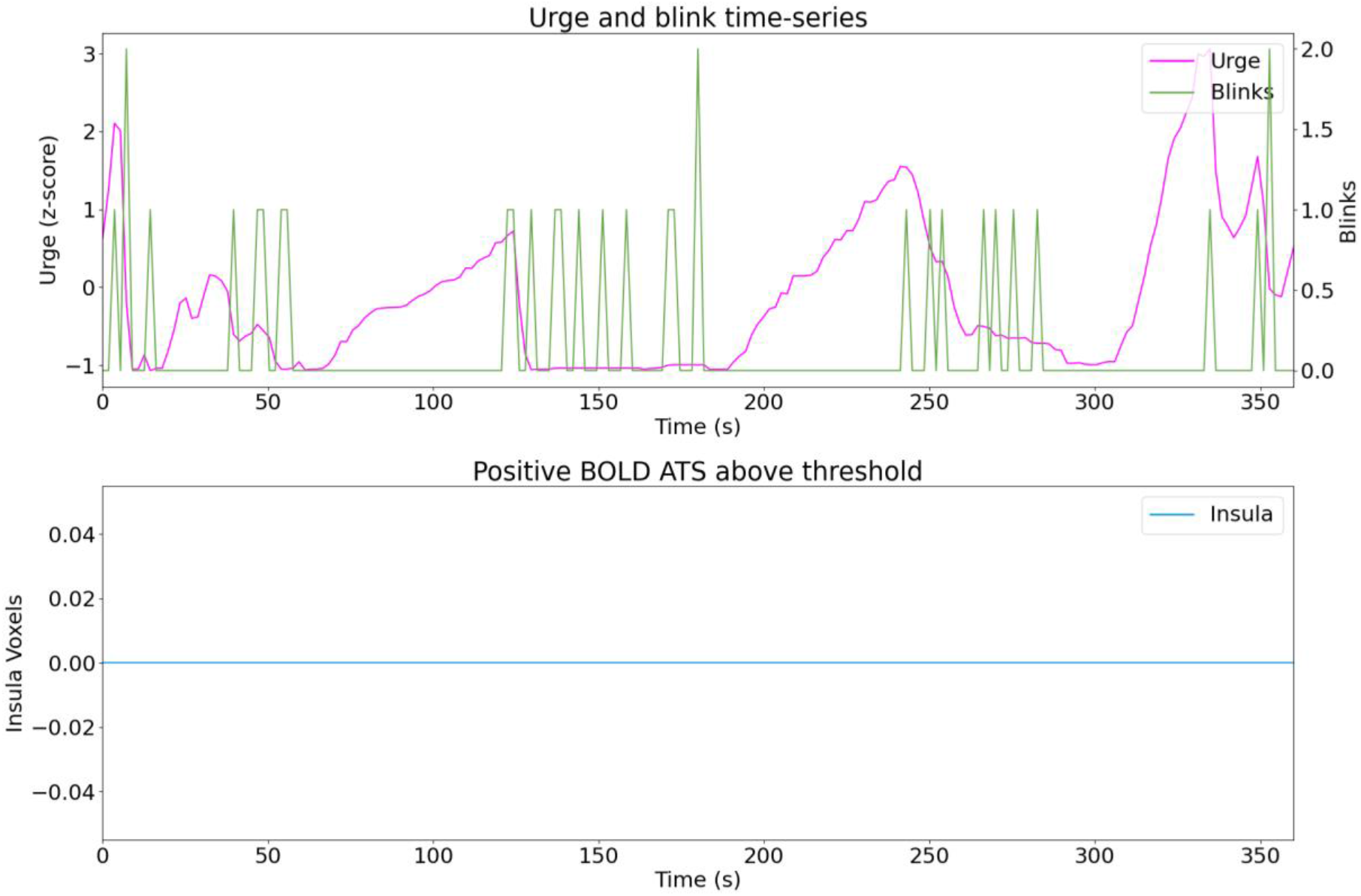
Sub13 run01.

**Figure C.27.**
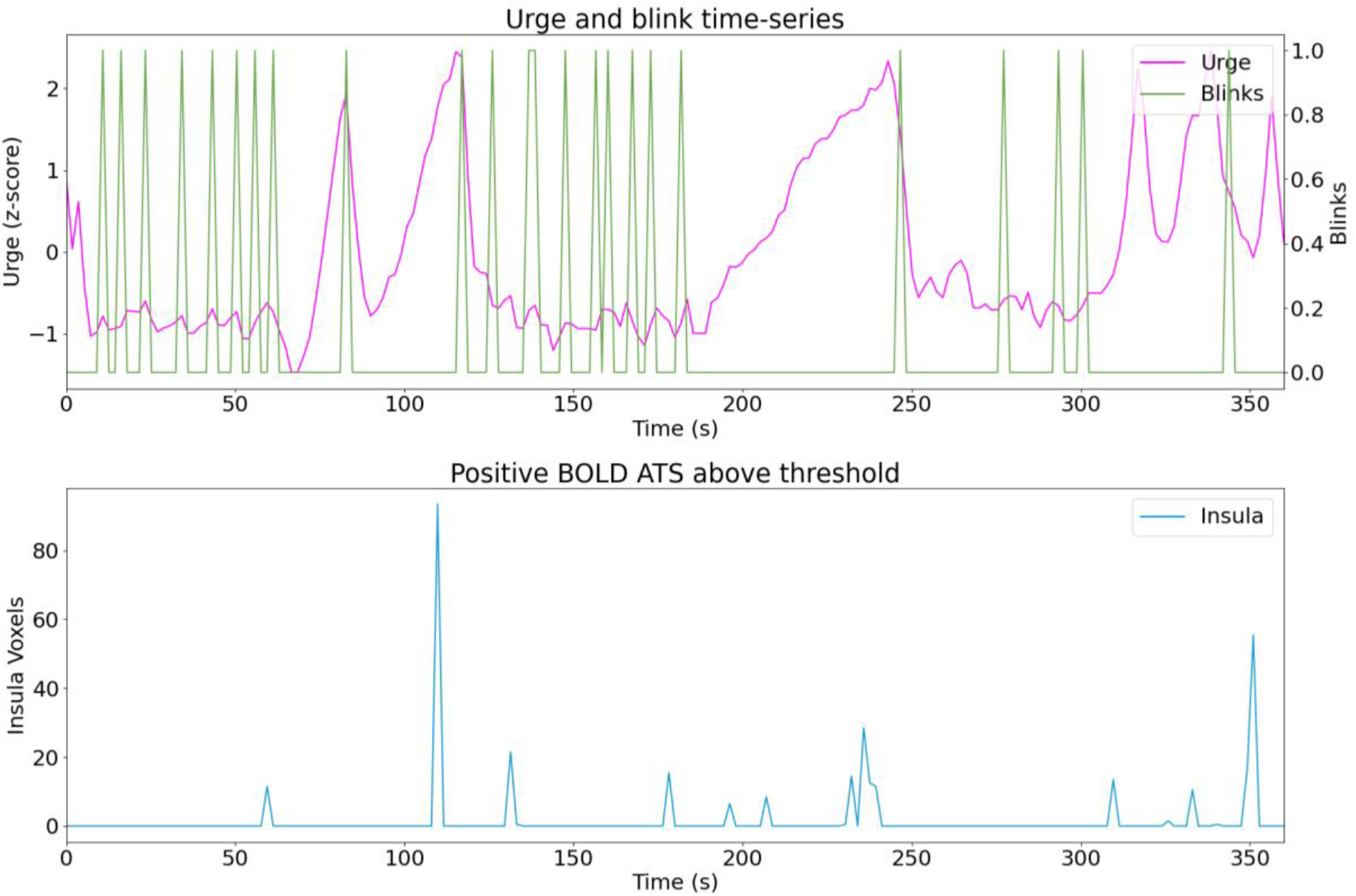
Sub13 run02.

**Figure C.28.**
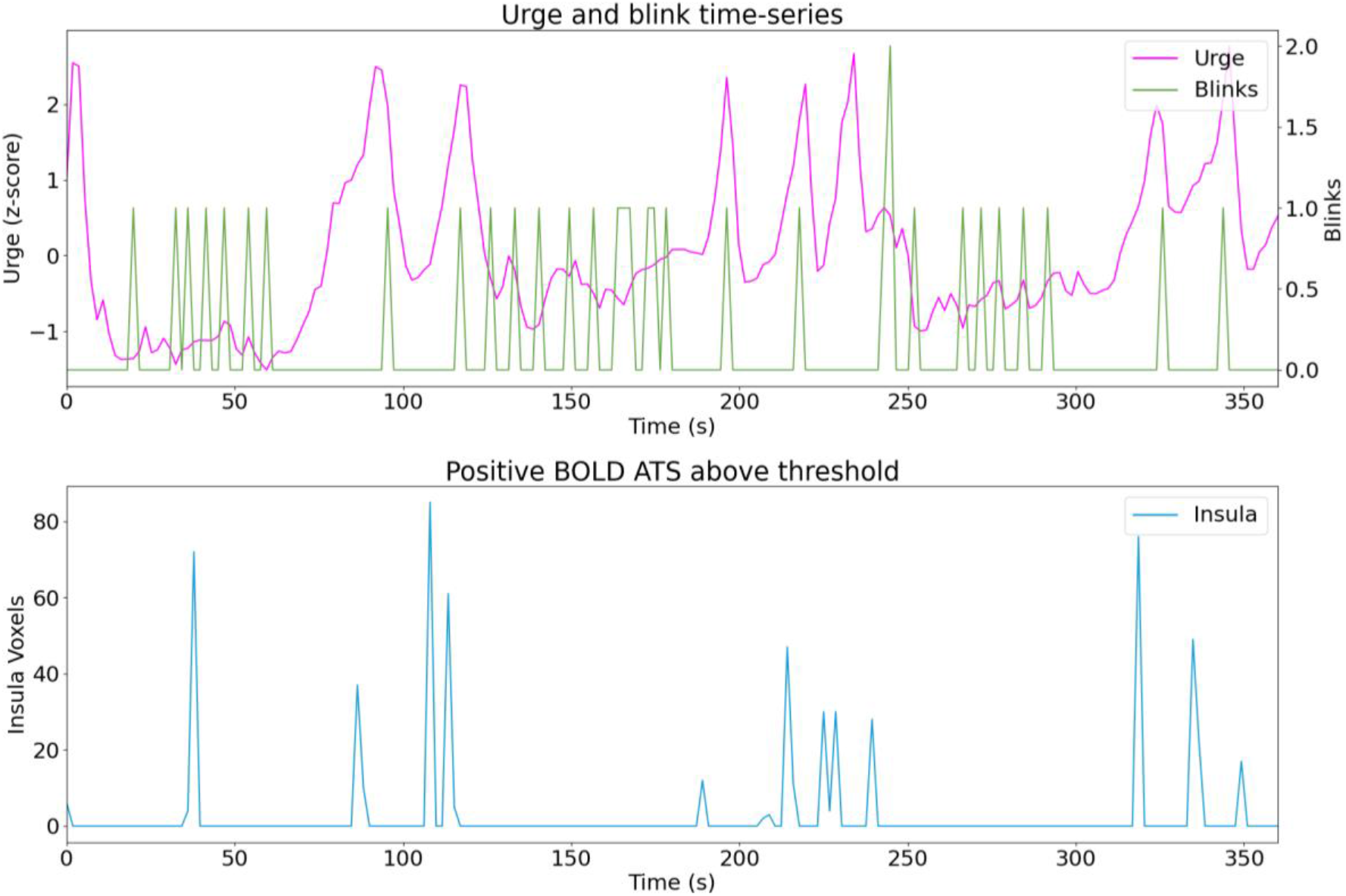
Sub13 run03.

**Figure C.29.**
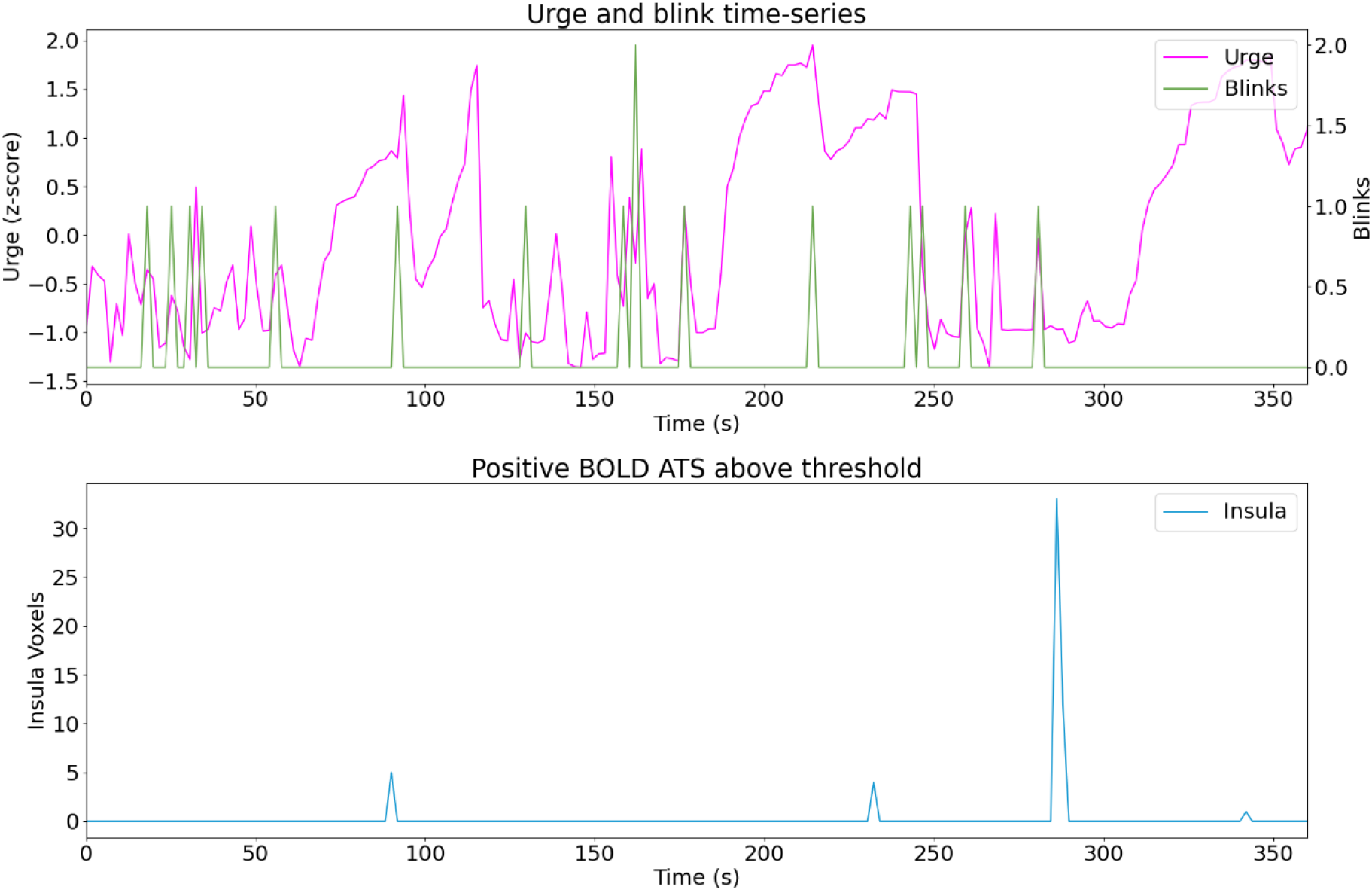
Sub14 run01.

**Figure C.30.**
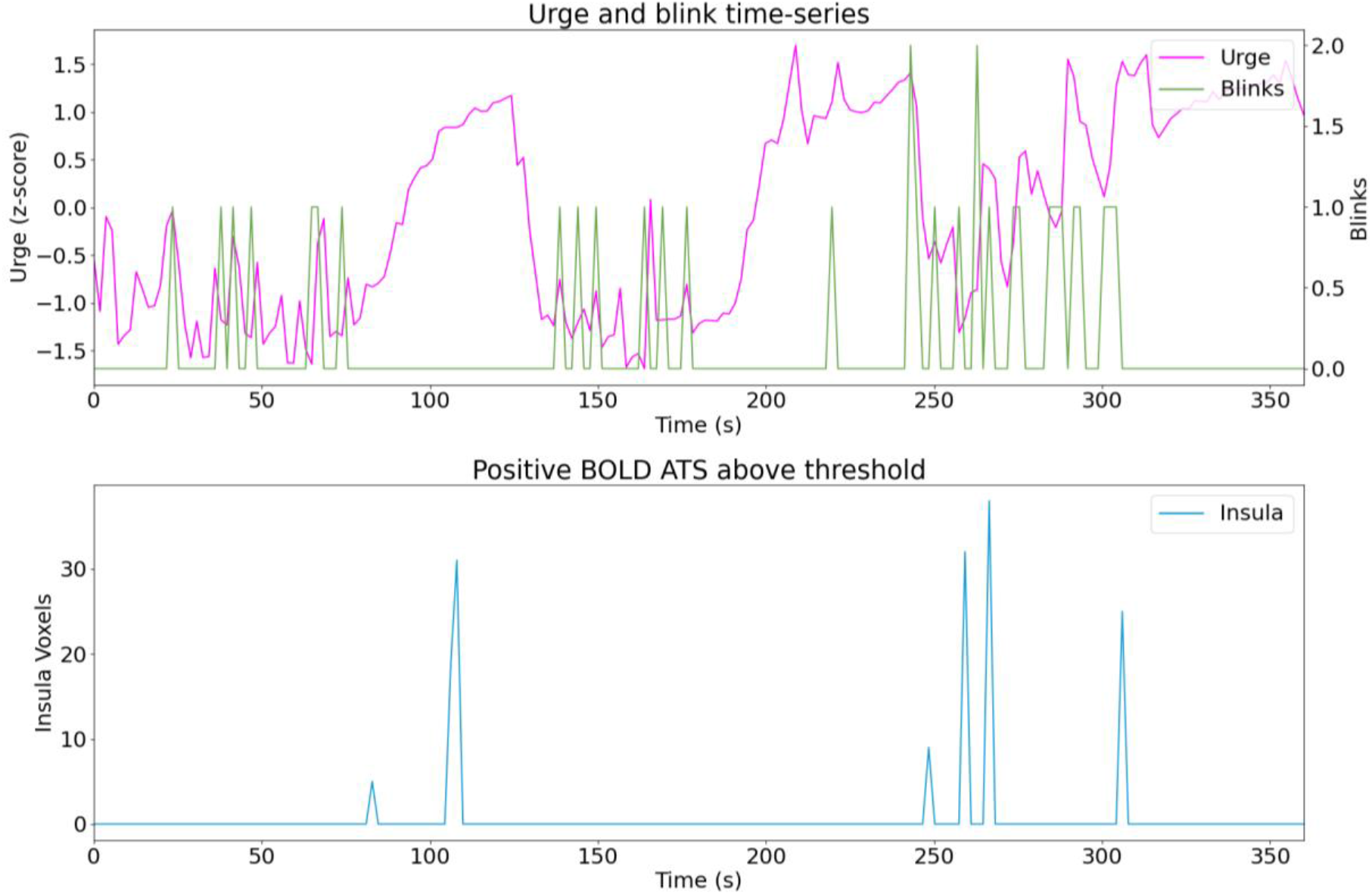
Sub14 run02.

**Figure C.31.**
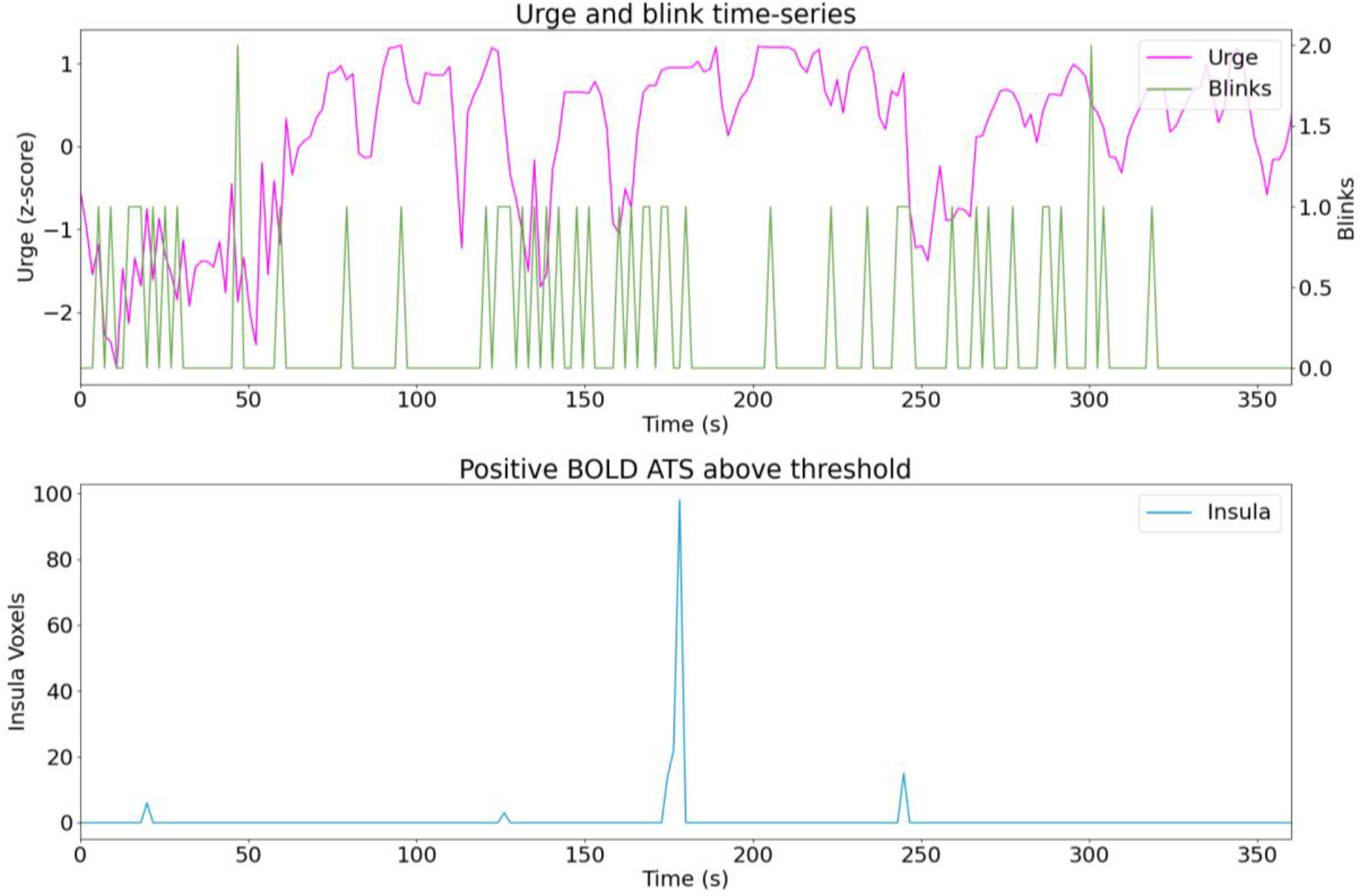
Sub14 run03.

**Figure C.32.**
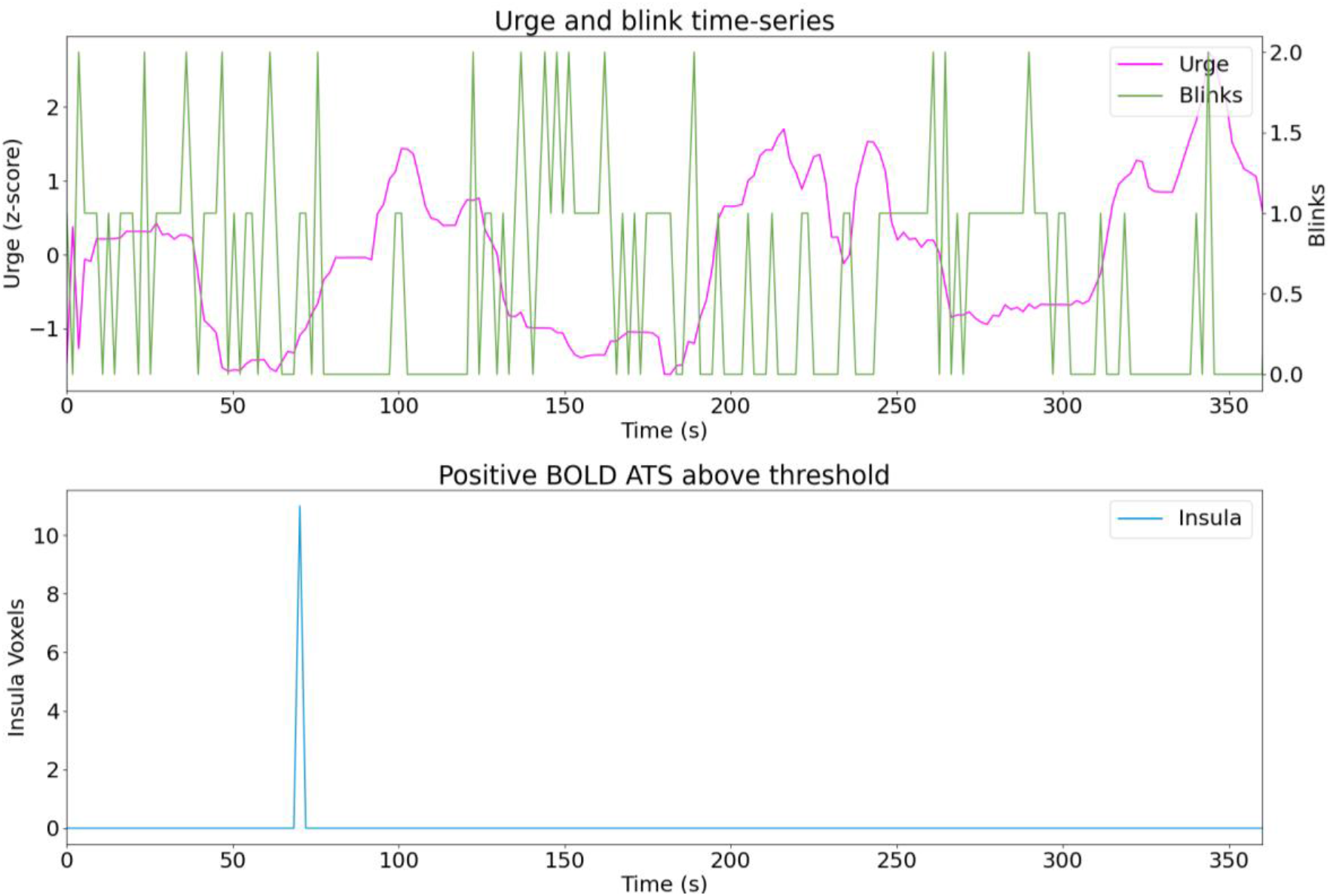
Sub15 run01.

**Figure C.33.**
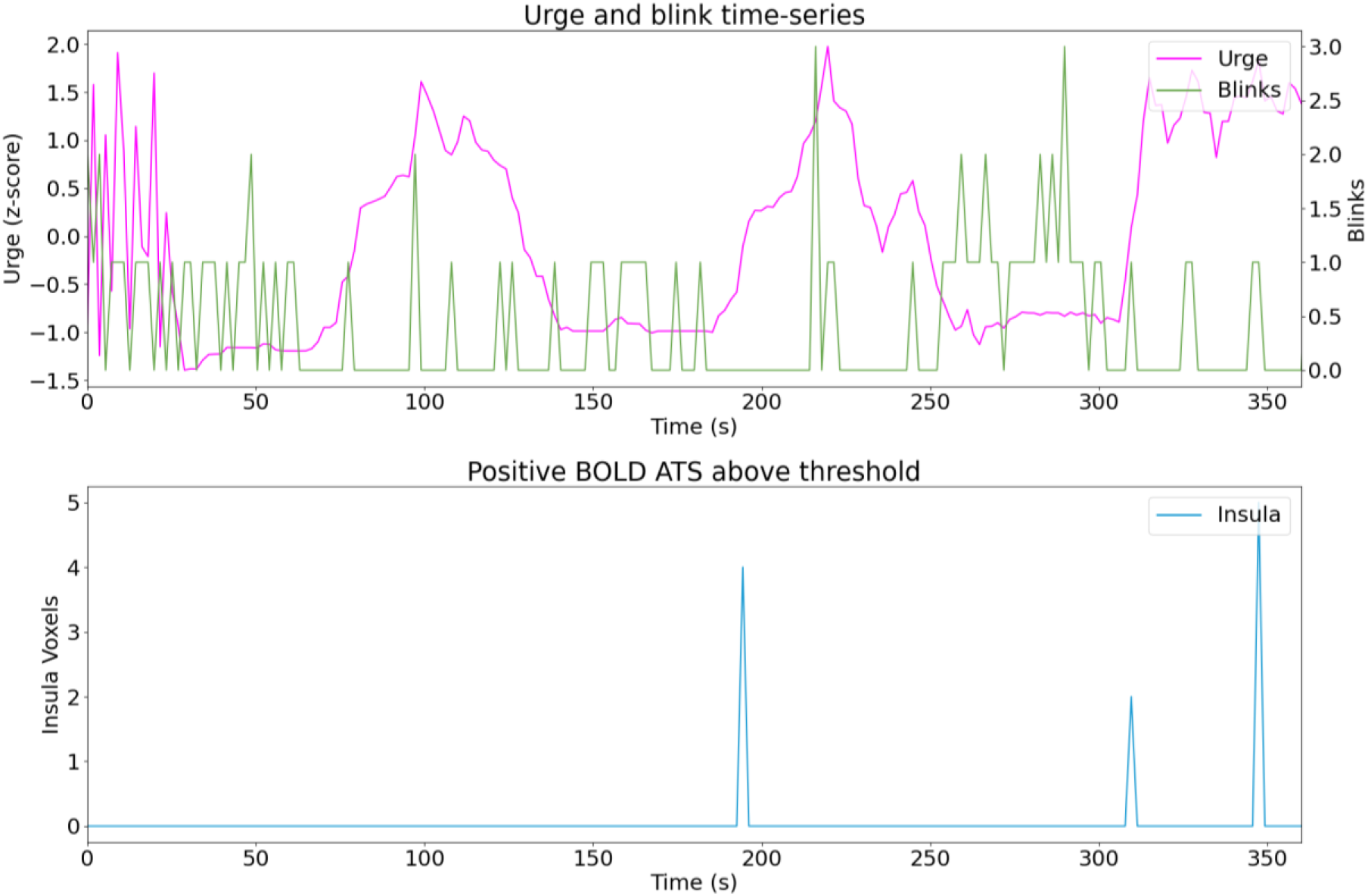
Sub15 run02.

**Figure C.34.**
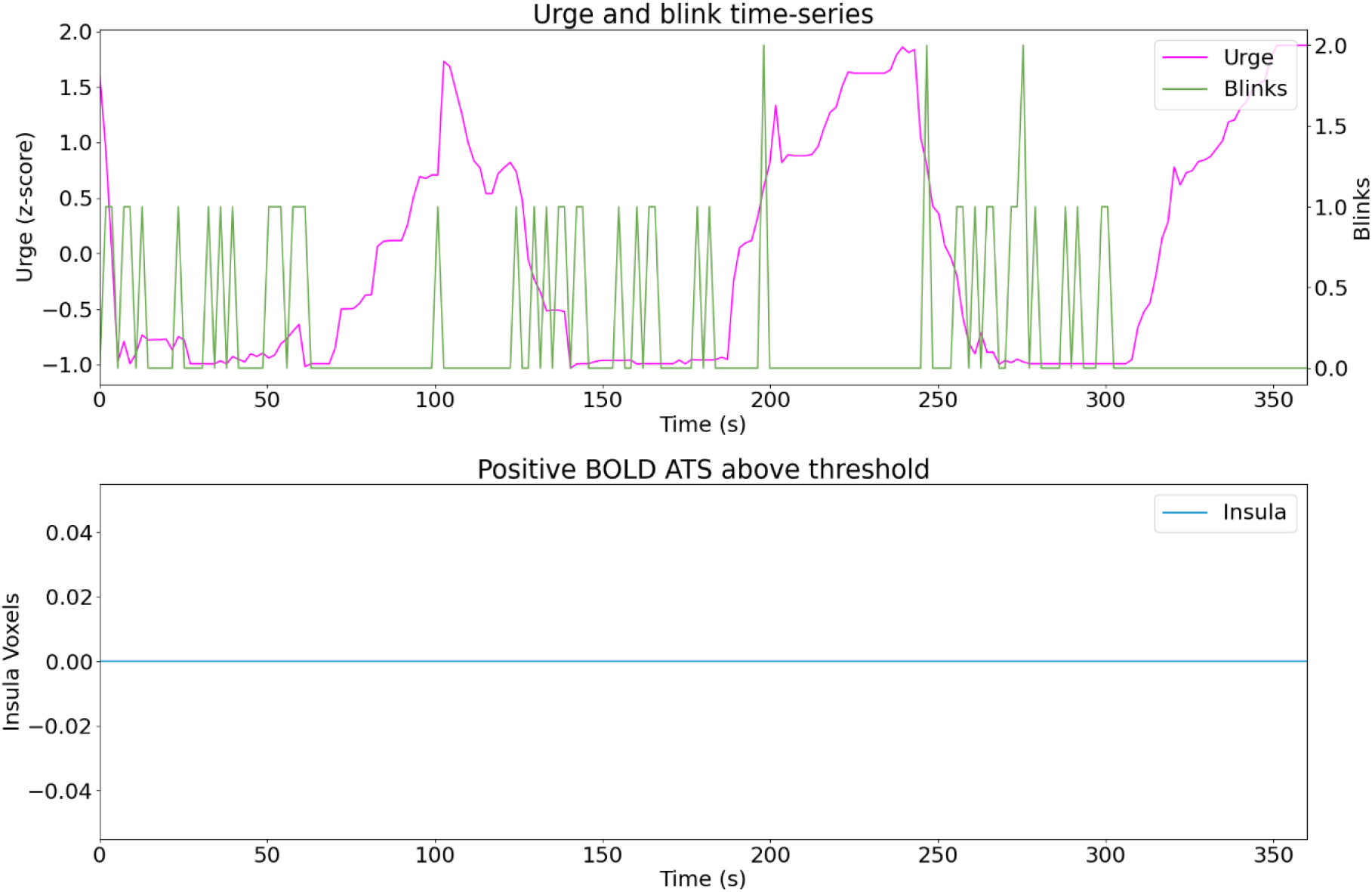
Sub15 run03.

**Figure C.35.**
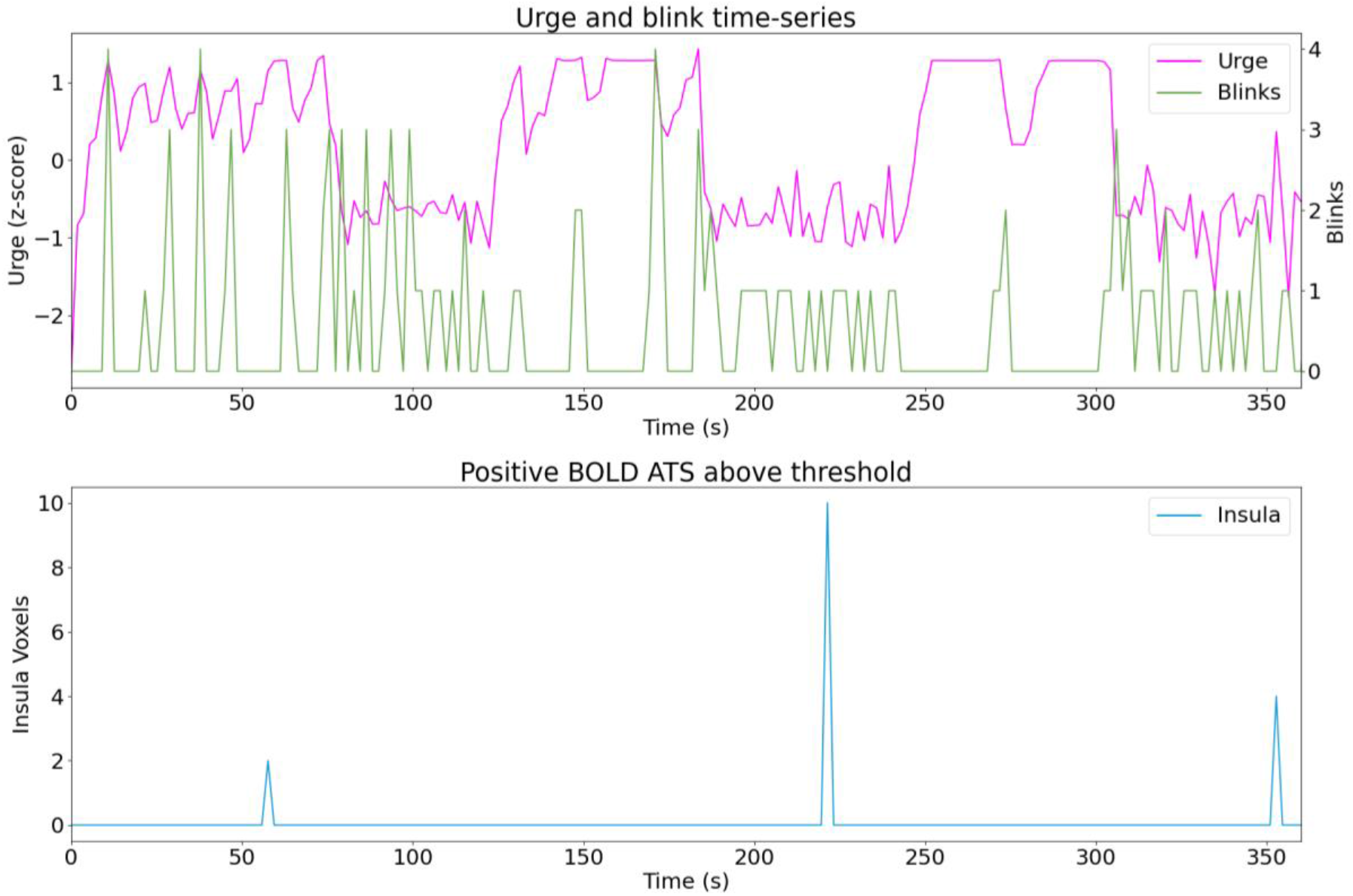
Sub16 run01.

**Figure C.36.**
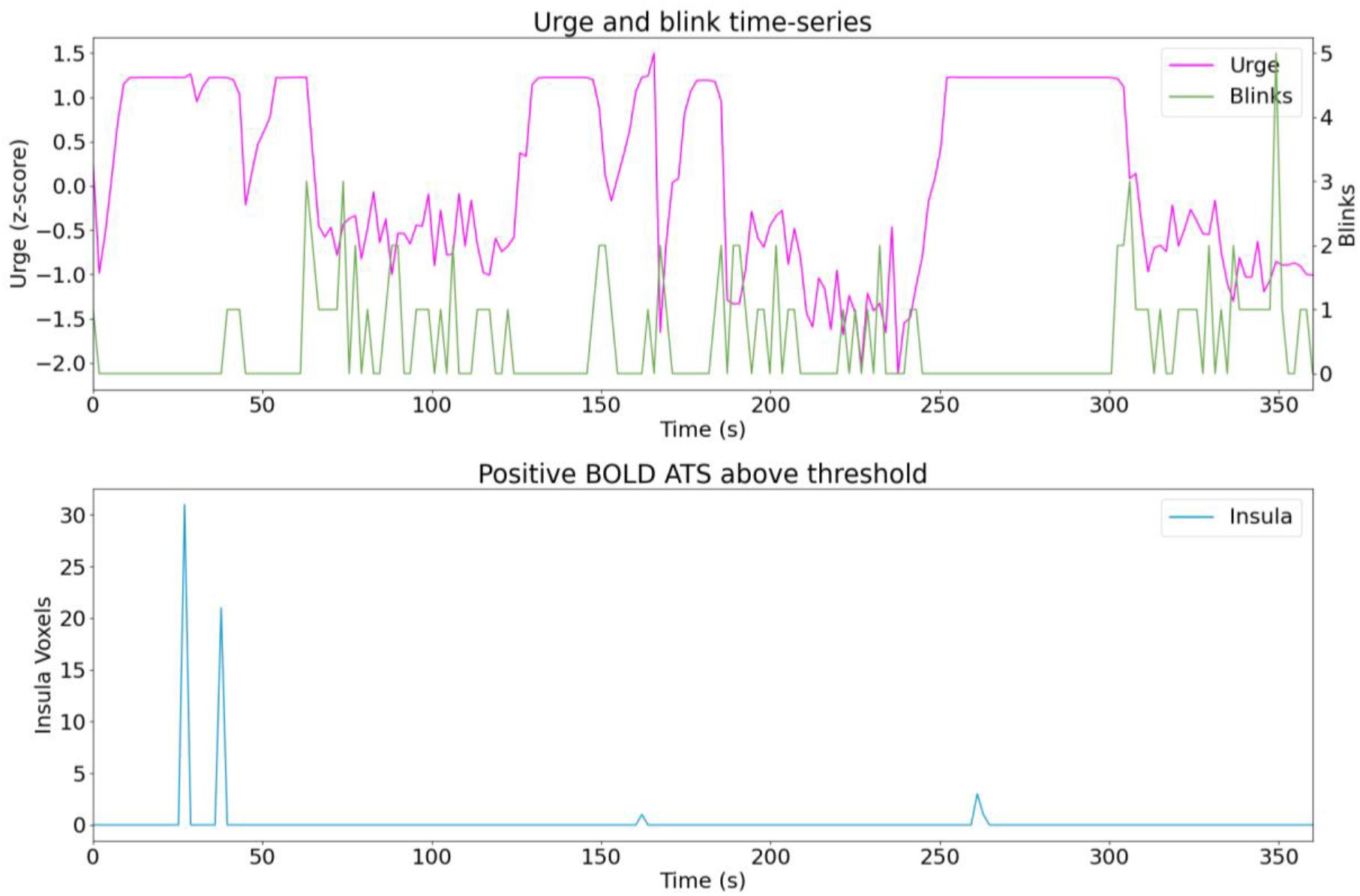
Sub16 run02.

**Figure C.37.**
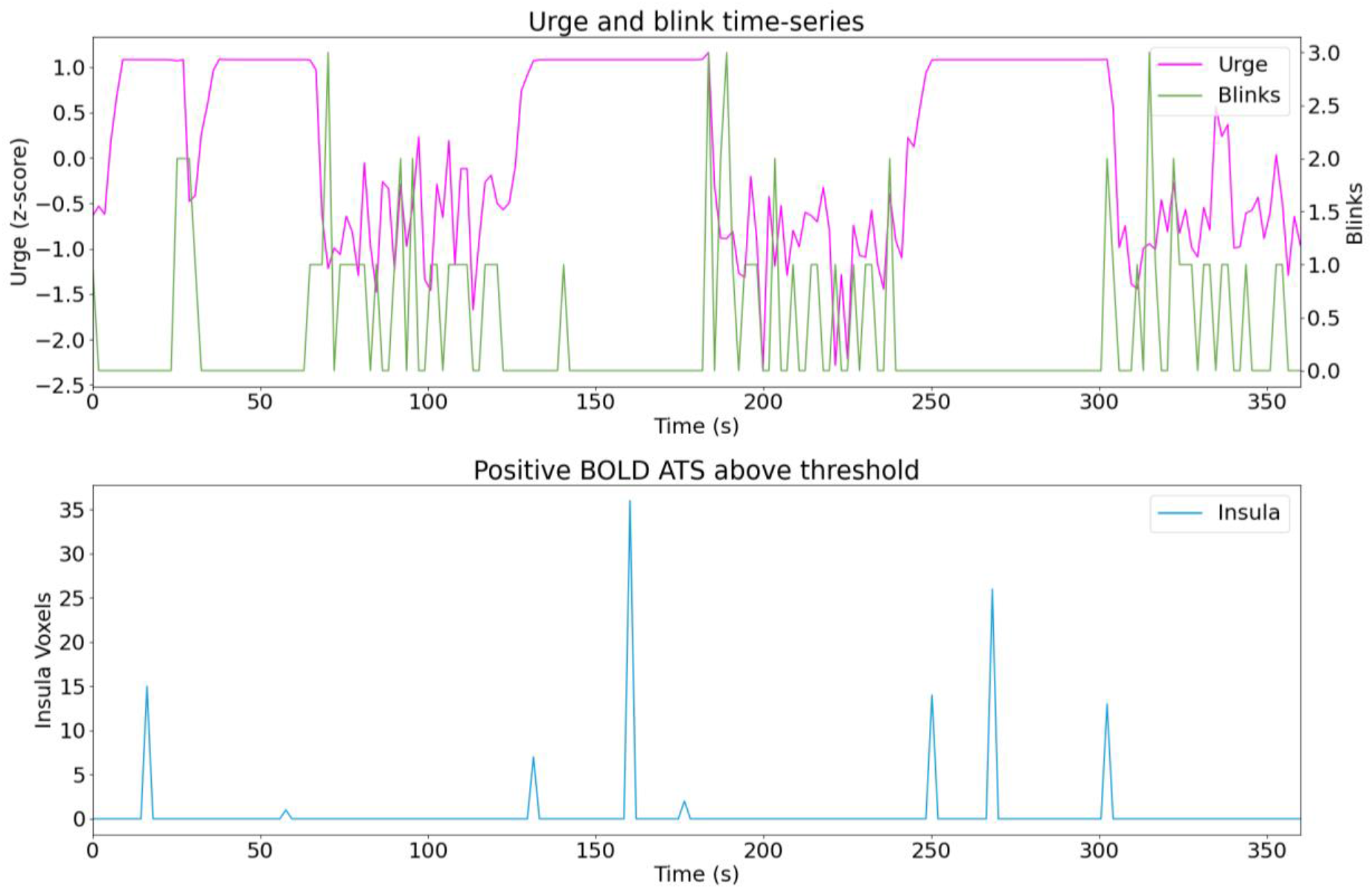
Sub16 run03.

**Figure C.38.**
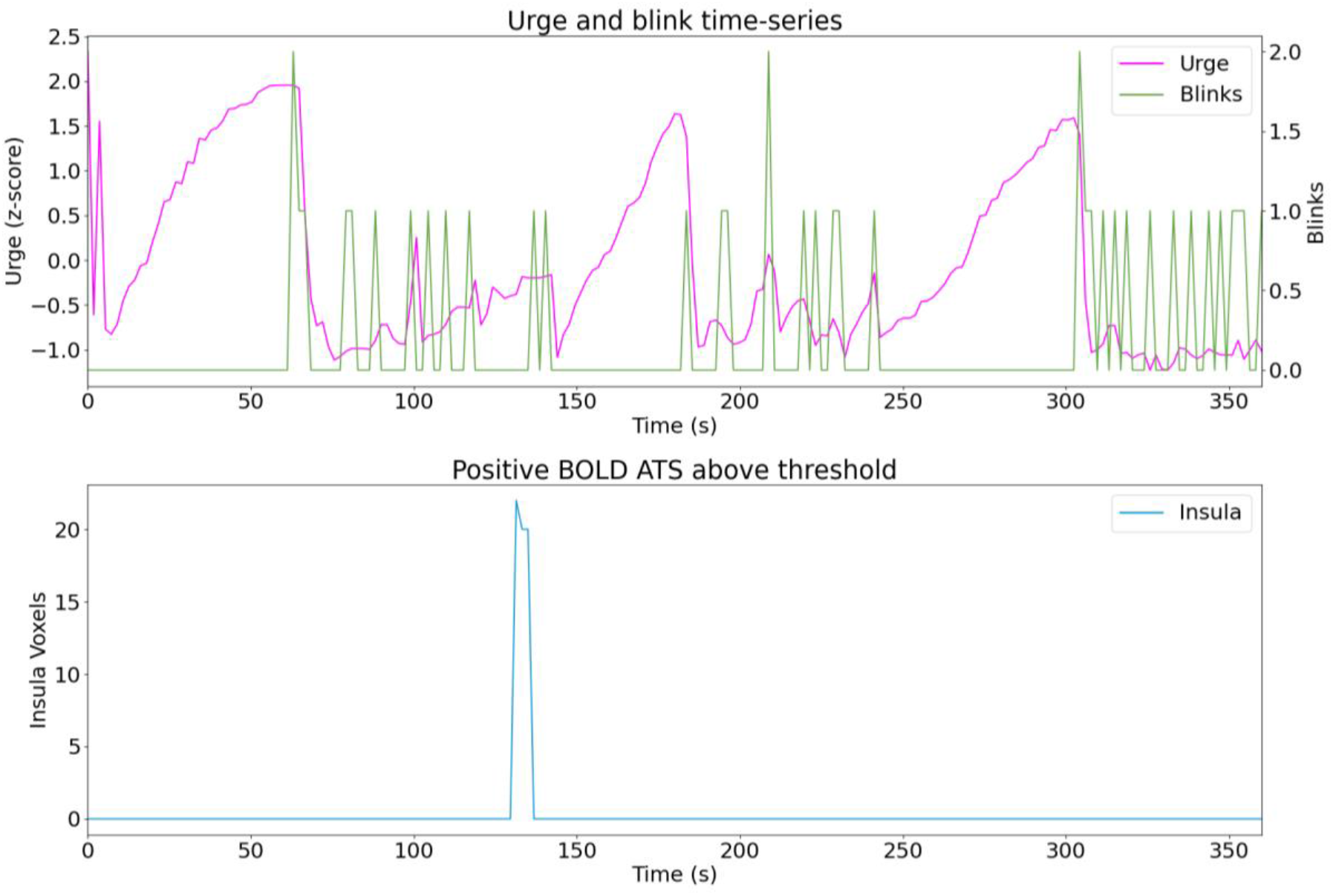
Sub17 run01.

**Figure C.39.**
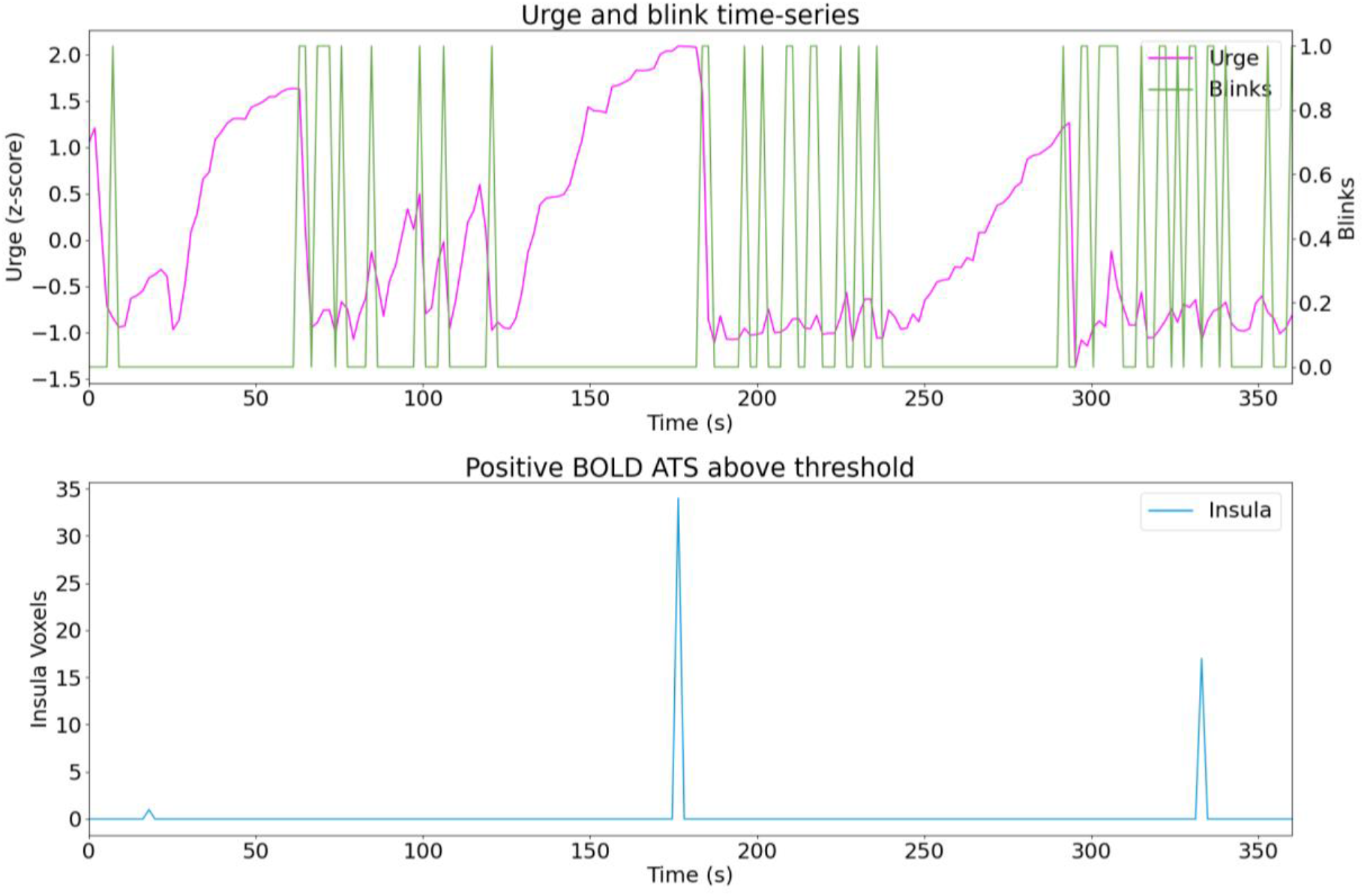
Sub17 run02.

**Figure C.40.**
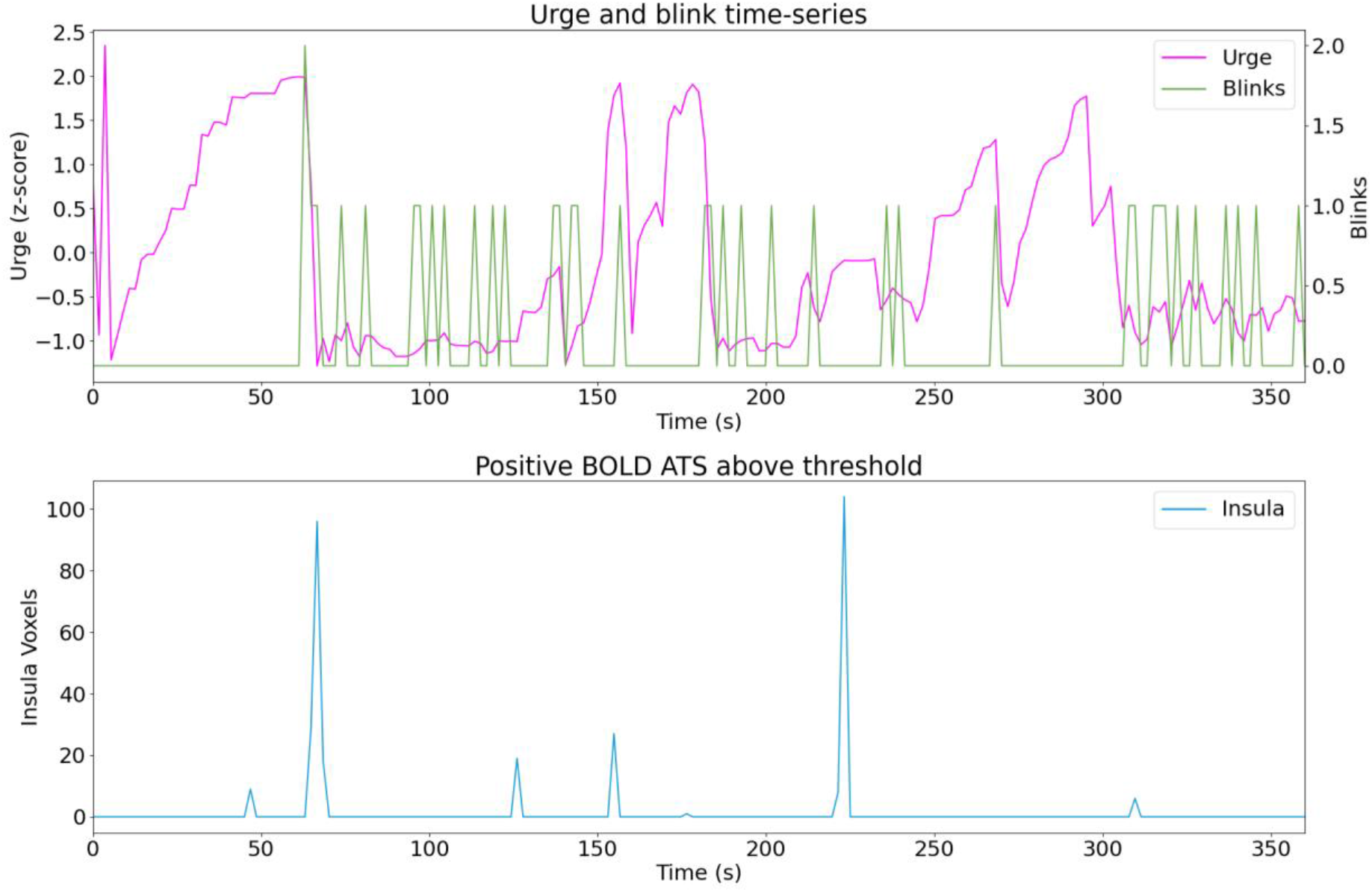
Sub17 run03.

**Figure C.41.**
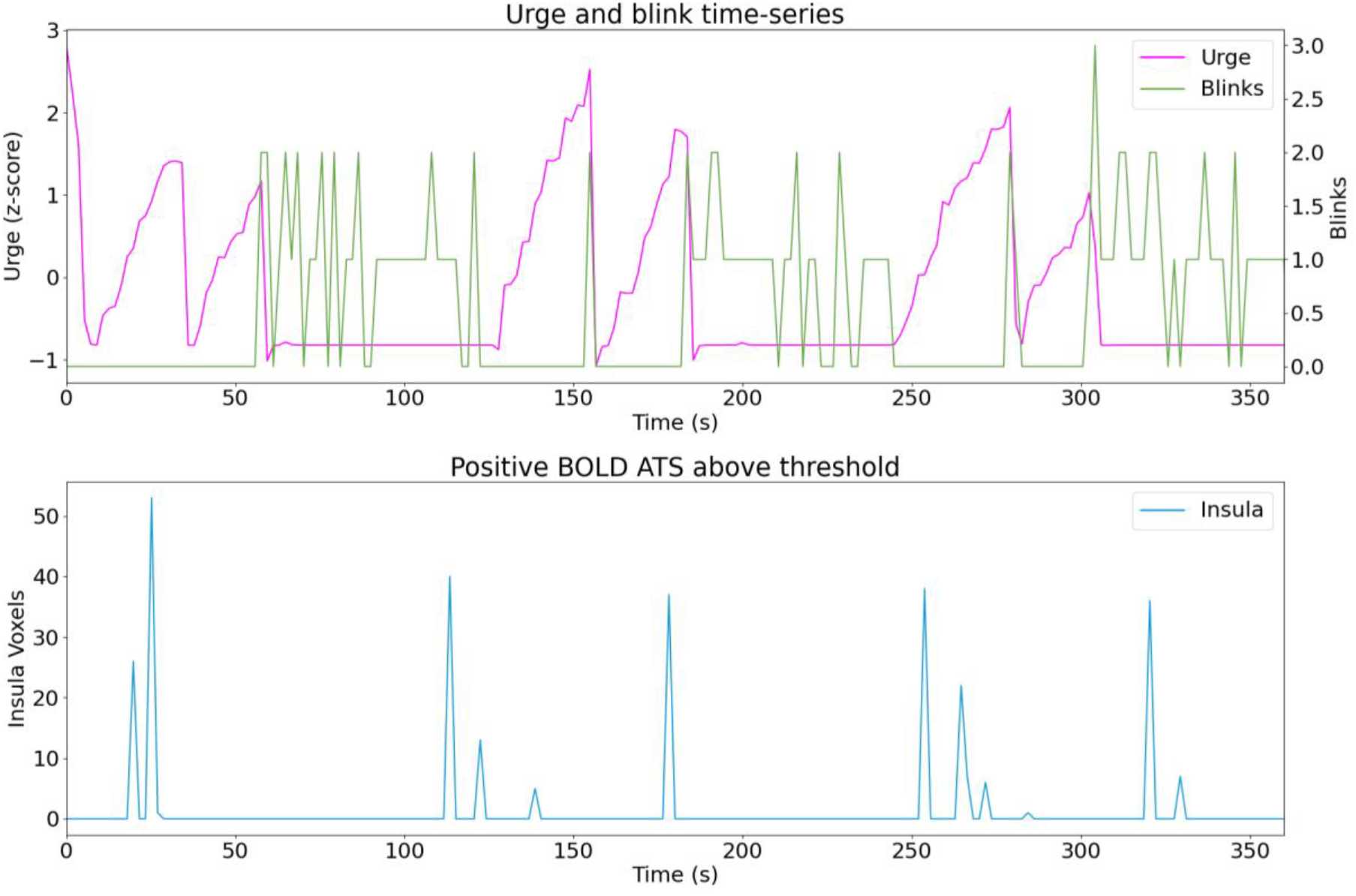
Sub18 run01.

**Figure C.42.**
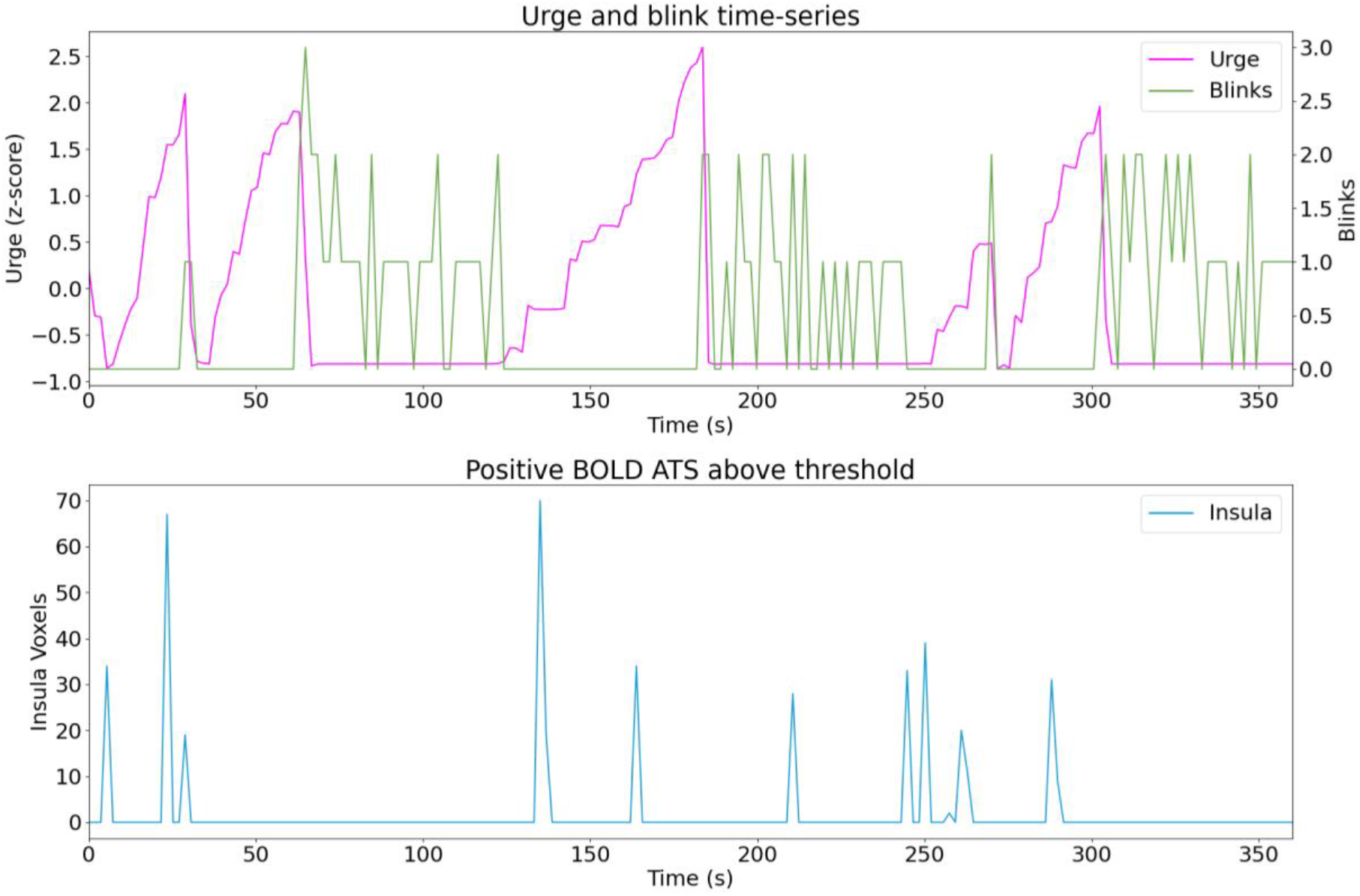
Sub18 run02.

**Figure C.43.**
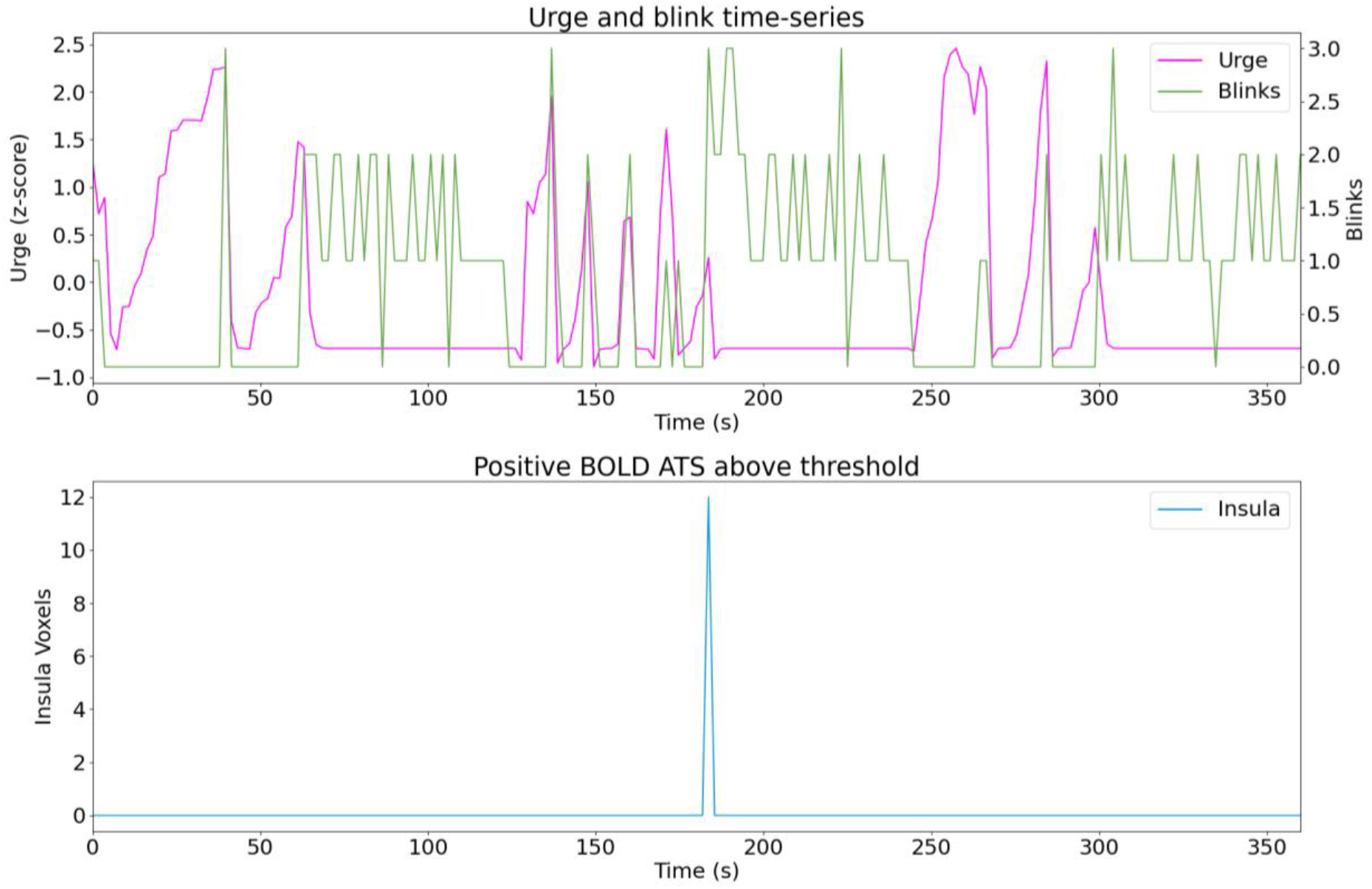
Sub18 run03.

**Figure C.44.**
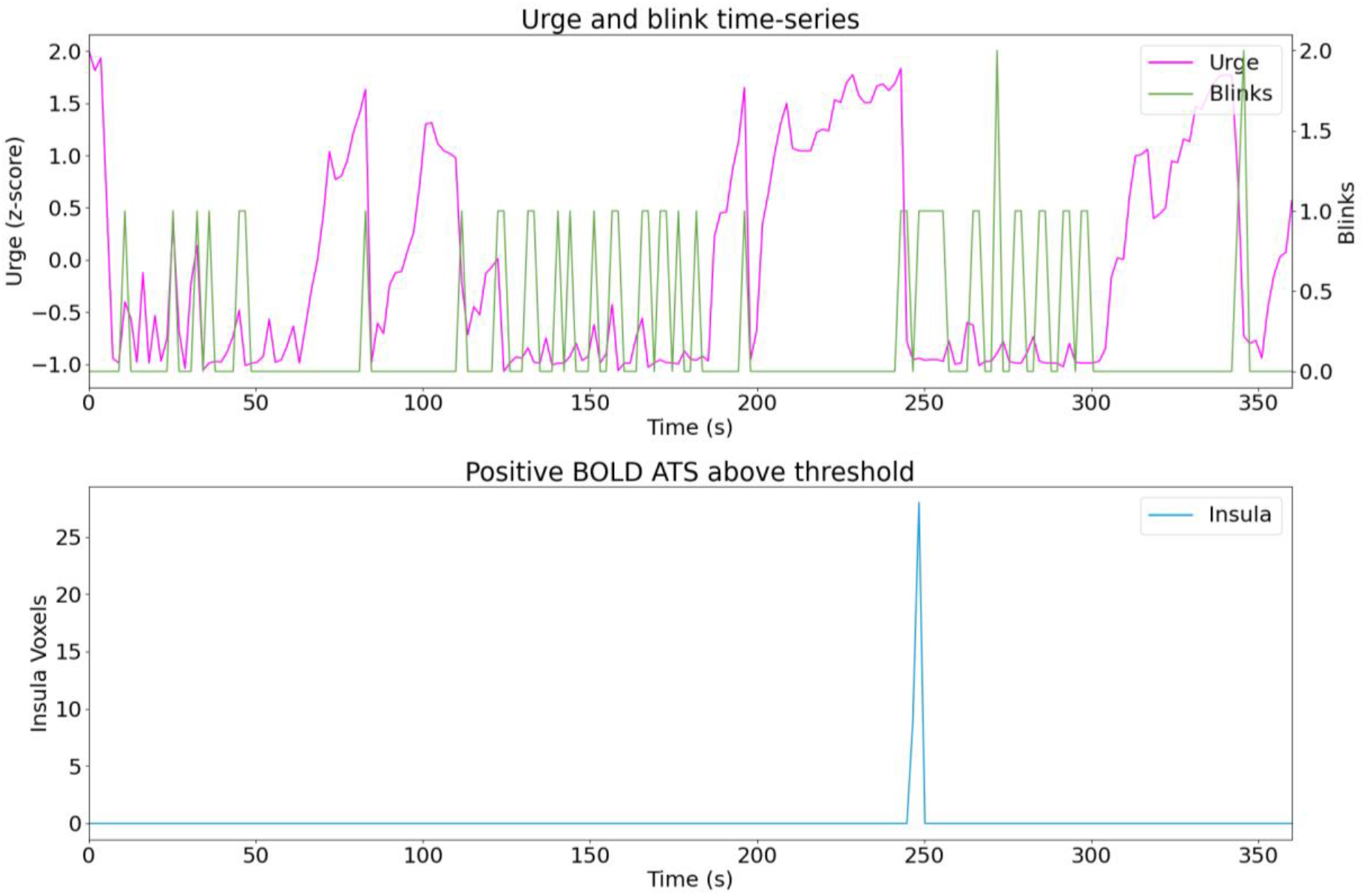
Sub19 run01.

**Figure C.45.**
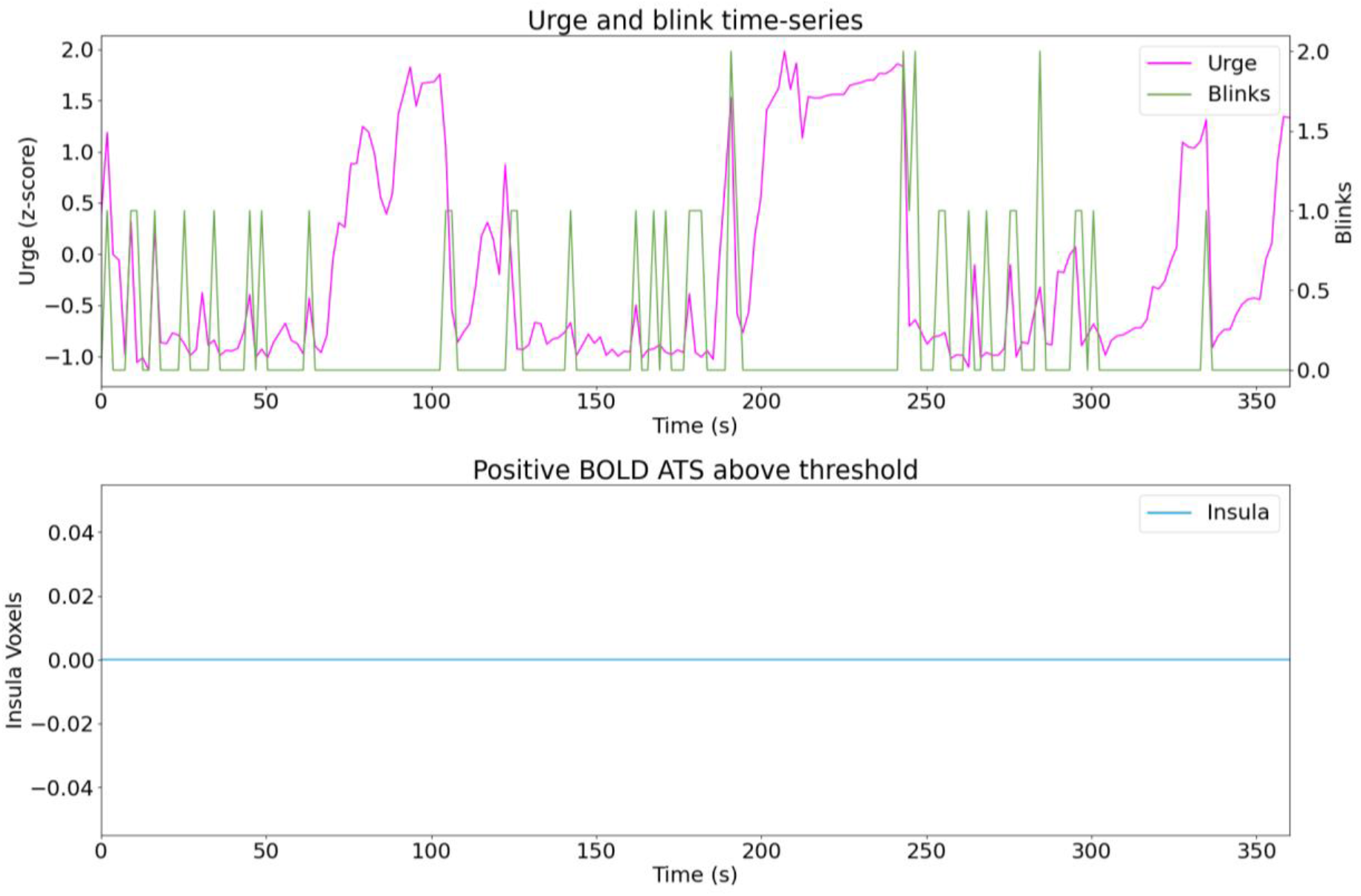
Sub19 run03.

**Figure C.46.**
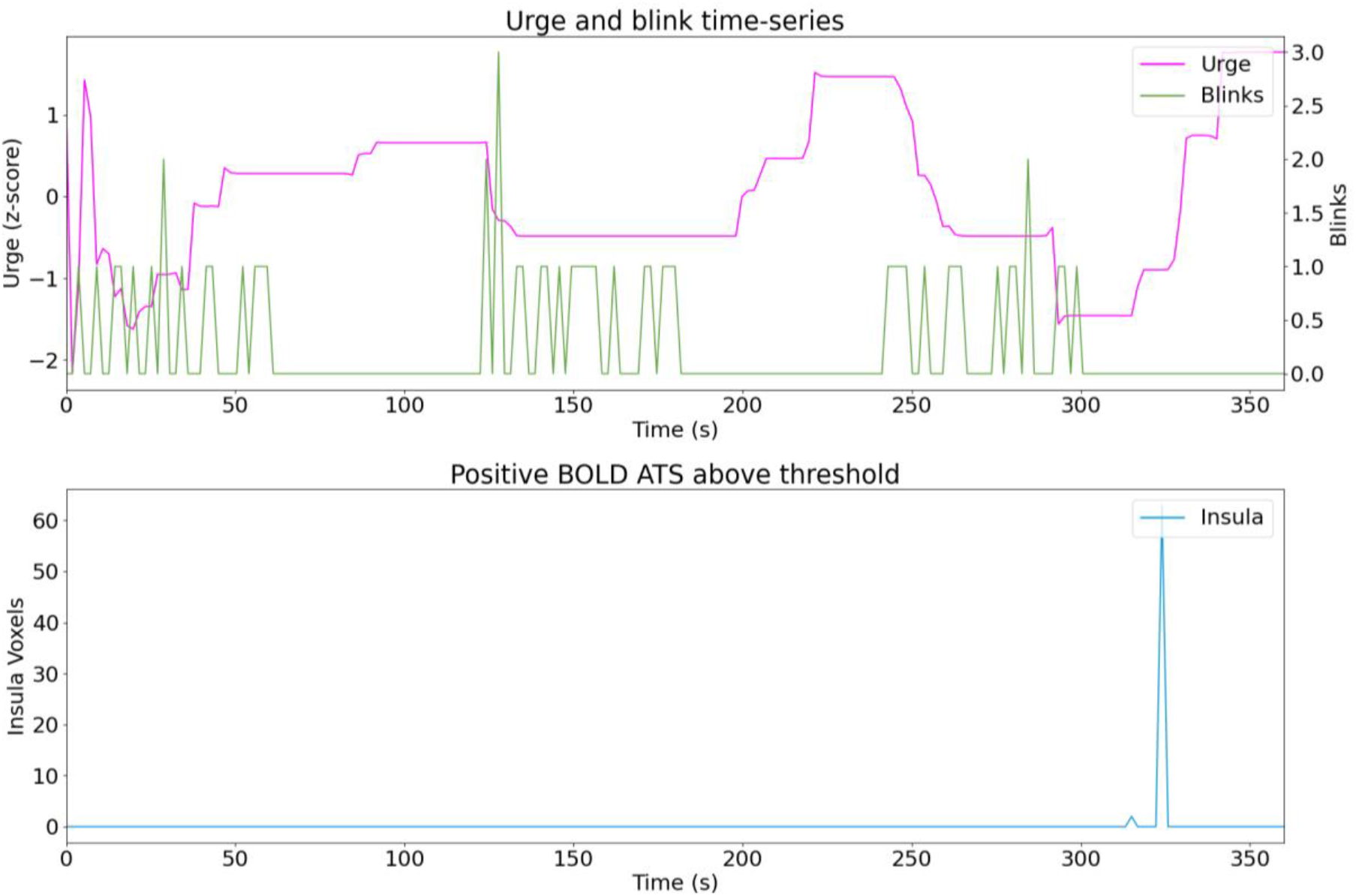
Sub20 run01.

**Figure C.47.**
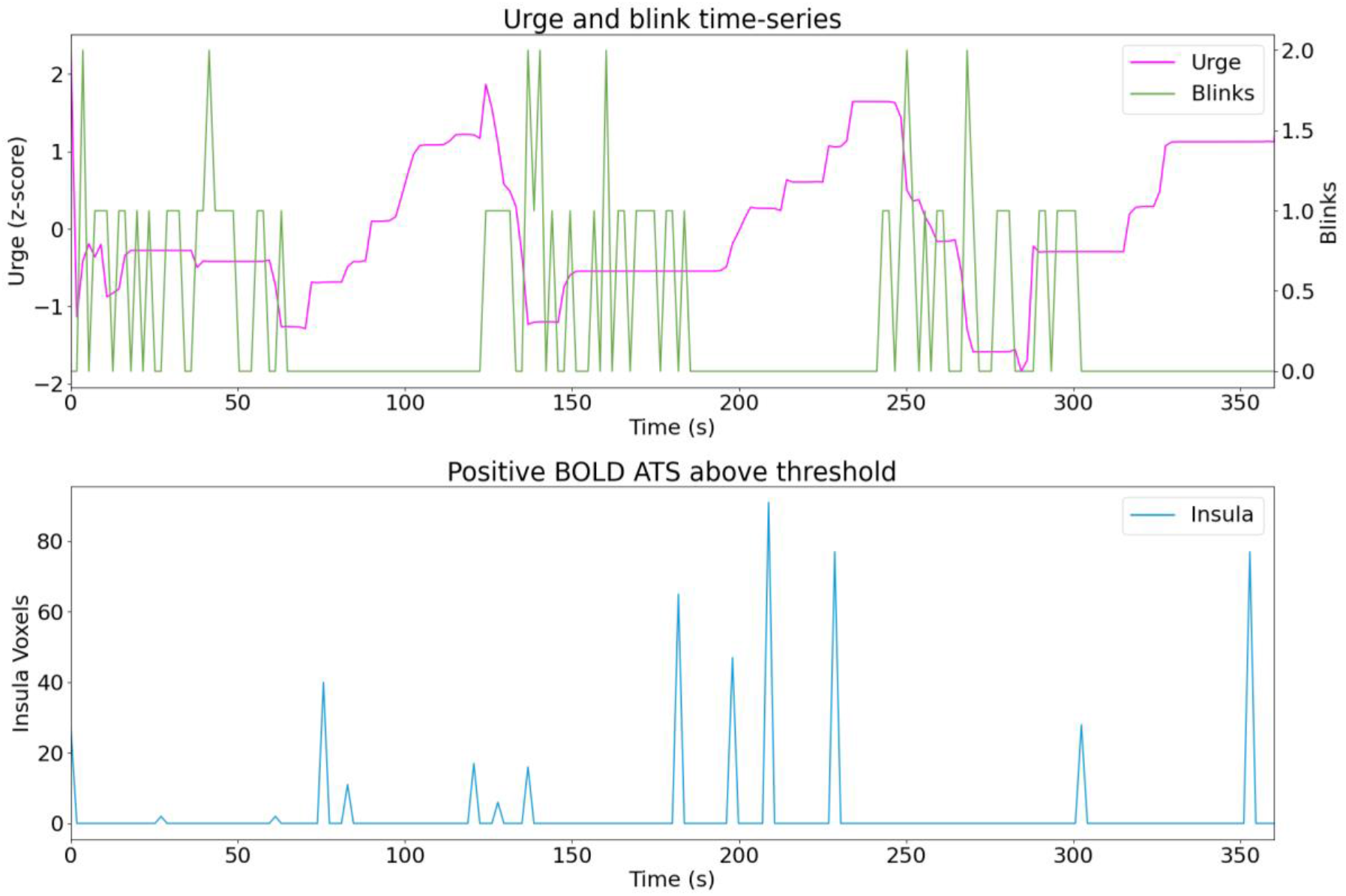
Sub20 run02.

**Figure C.48.**
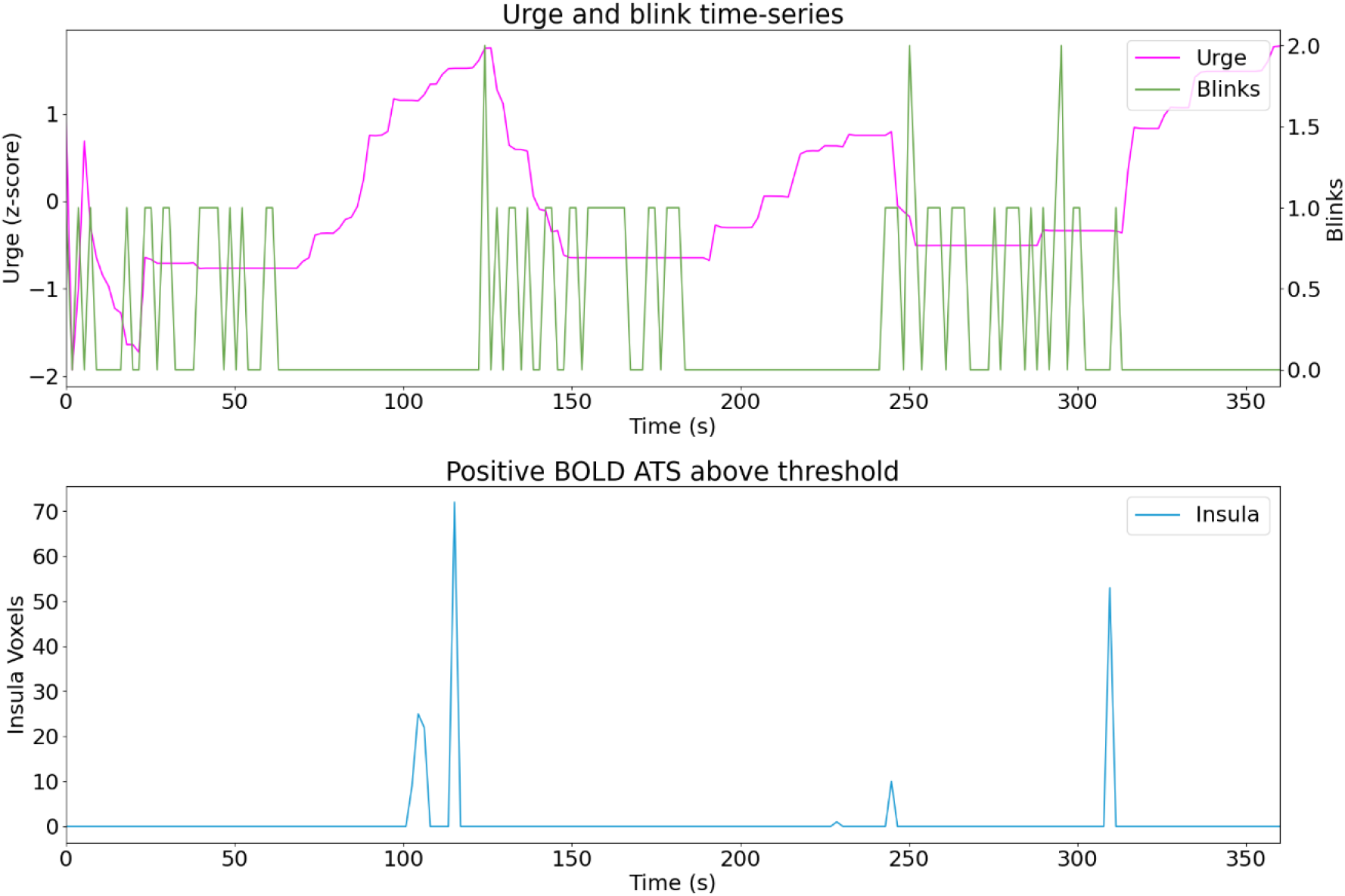
Sub20 run03.

**Figure C.49.**
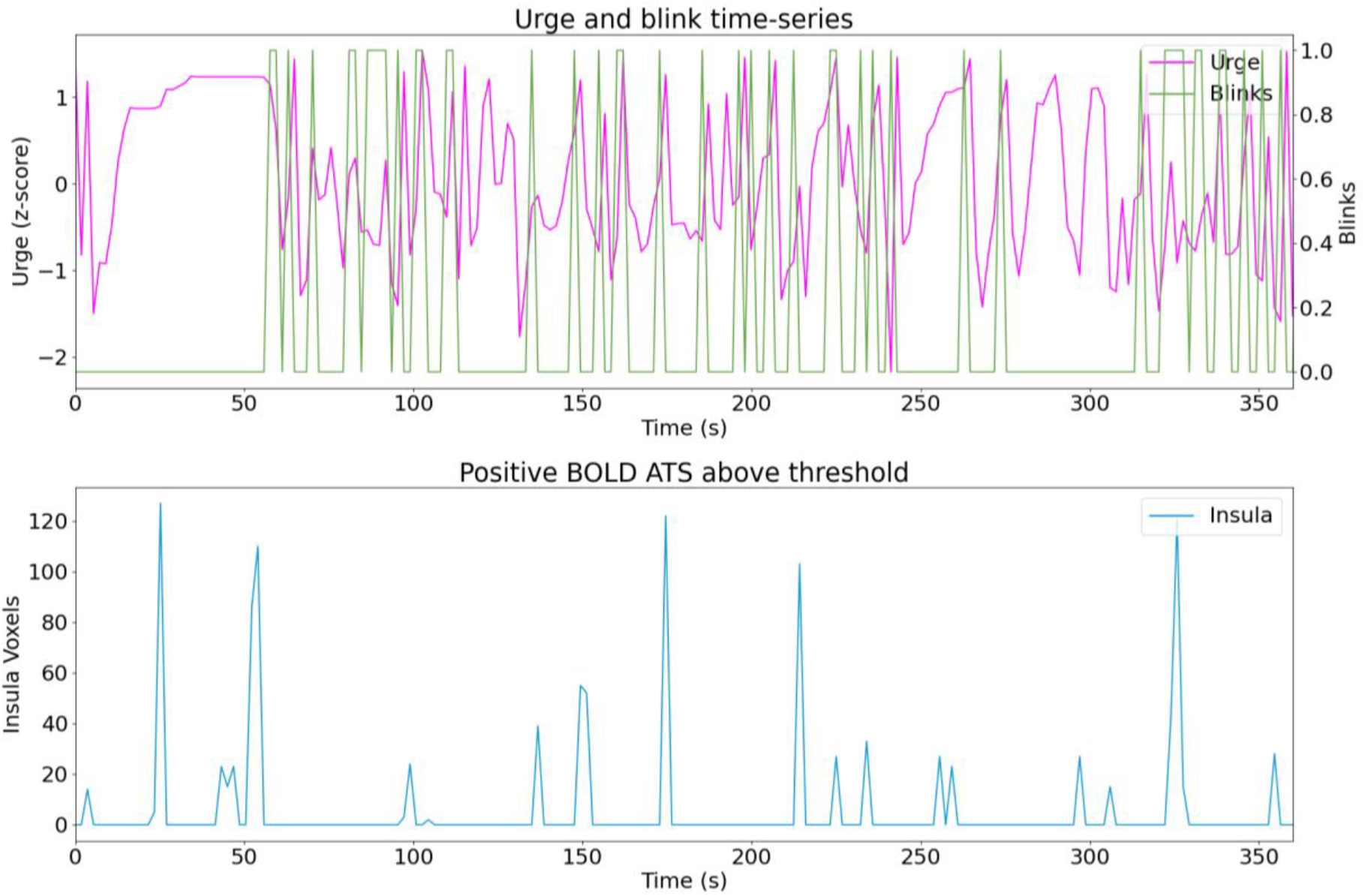
Sub21 run03.

**Figure C.50.**
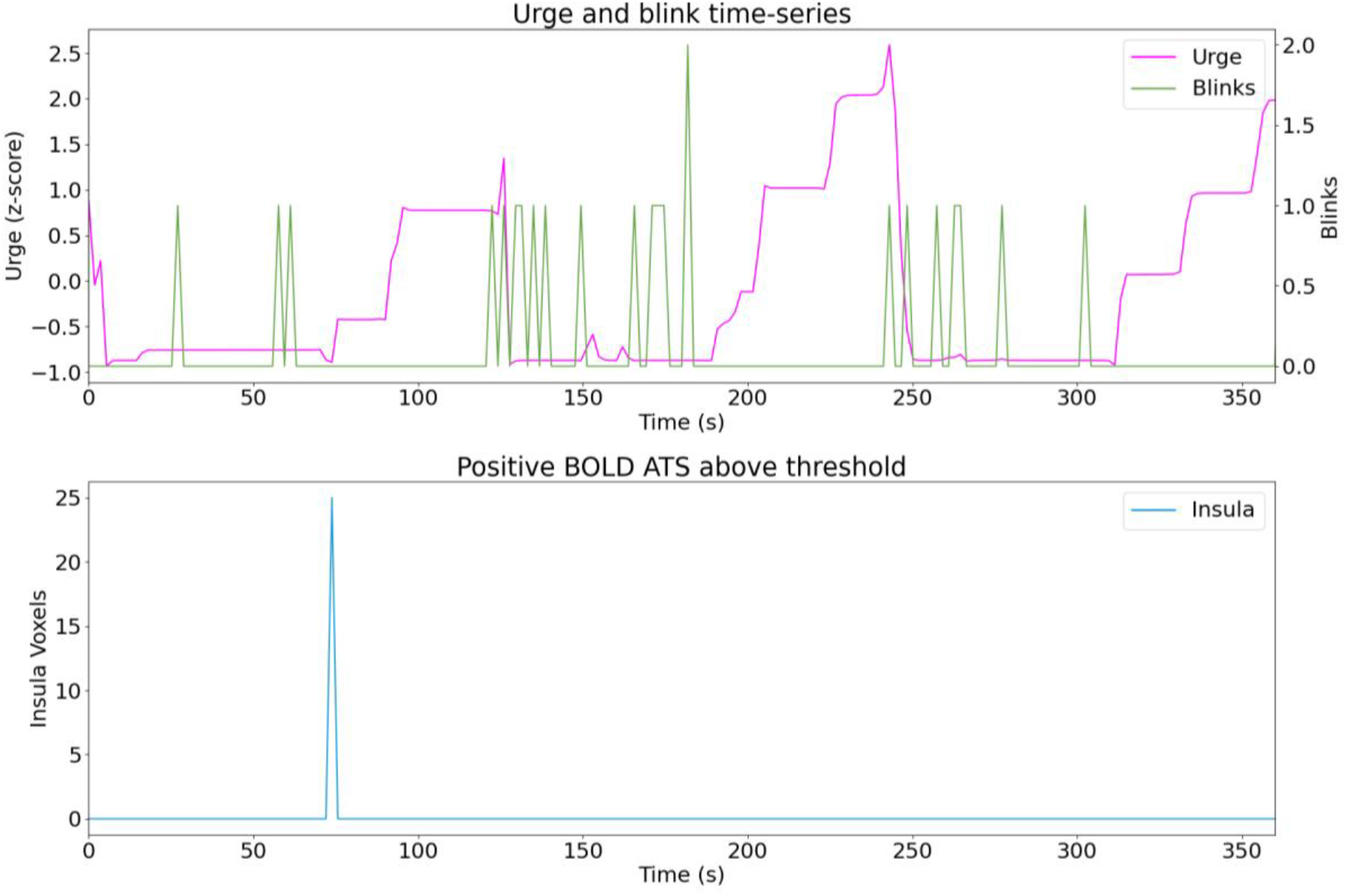
Sub22 run01.

**Figure C.51.**
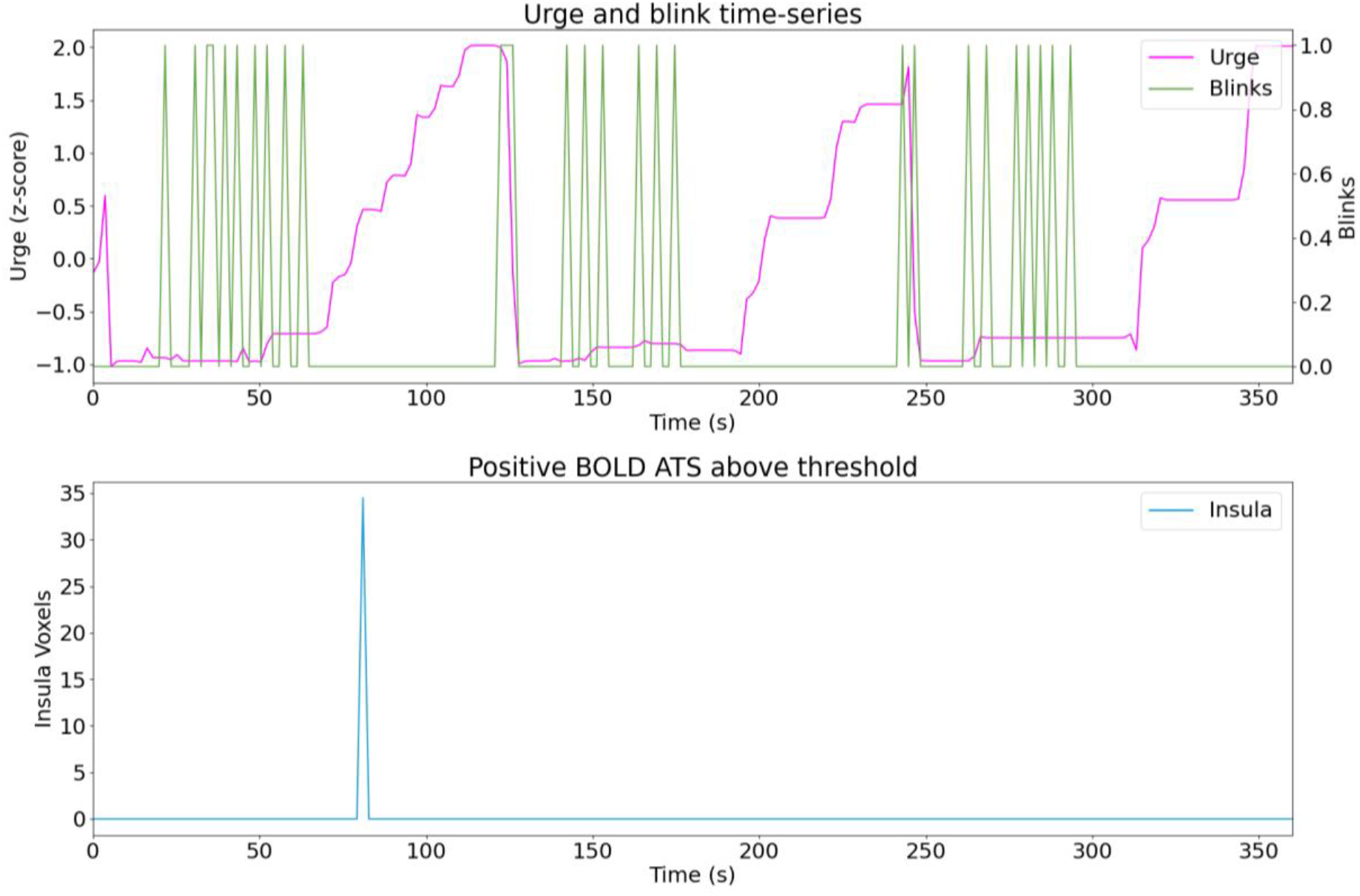
Sub22 run02.

**Figure C.52.**
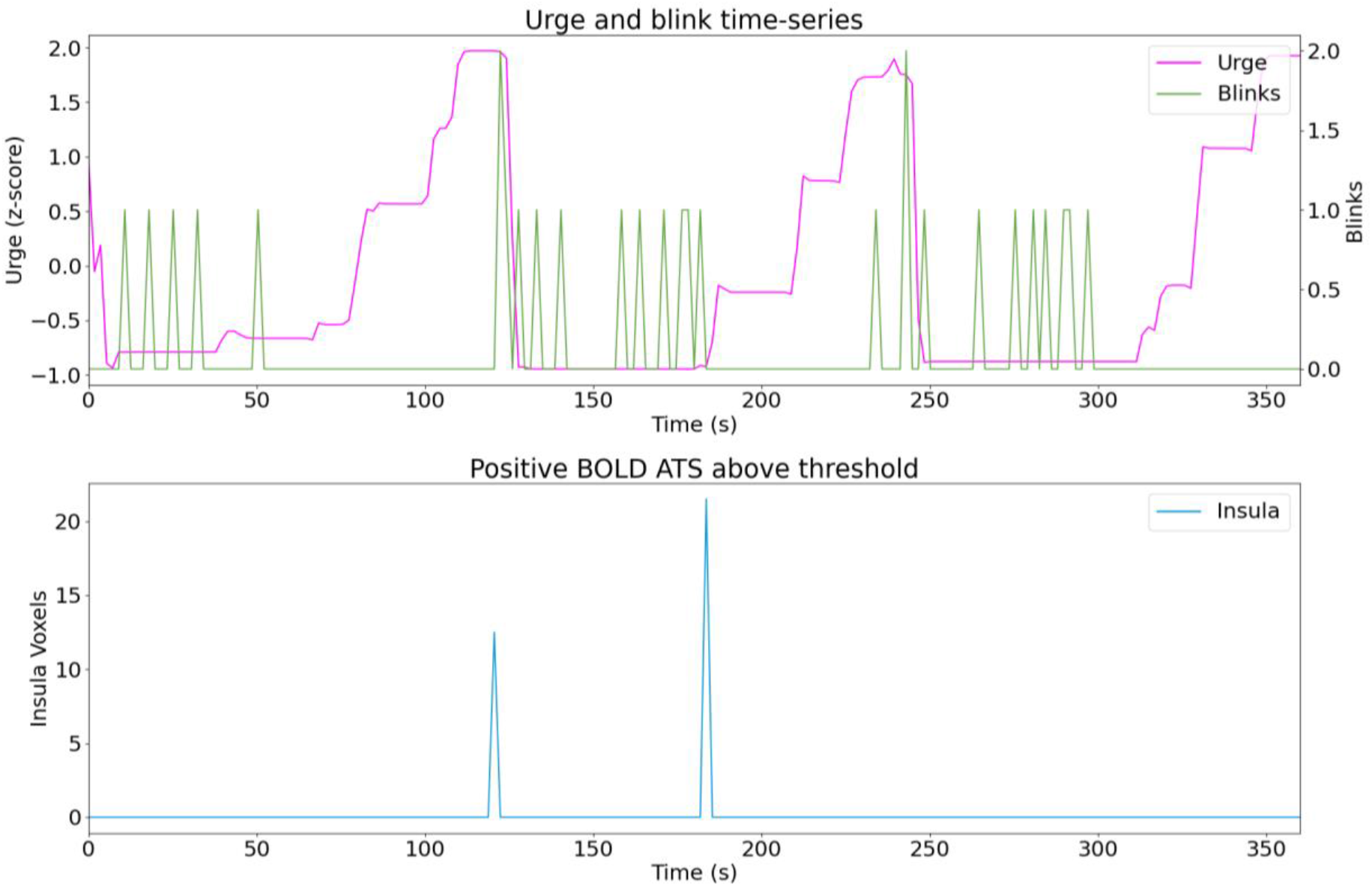
Sub22 run03.

